# Two functionally distinct heme/iron transport systems are virulence determinants of the fish pathogen *Flavobacterium psychrophilum*

**DOI:** 10.1101/2022.04.04.486927

**Authors:** Yueying Zhu, Delphine Lechardeur, Jean-François Bernardet, Brigitte Kerouault, Cyprien Guérin, Dimitri Rigaudeau, Pierre Nicolas, Eric Duchaud, Tatiana Rochat

## Abstract

Bacterial pathogens have a critical impact on aquaculture, a sector that accounts for half of the human fish consumption. *Flavobacterium psychrophilum* (phylum *Bacteroidetes*) is responsible for bacterial cold-water disease in salmonids worldwide. The molecular factors involved in host invasion, colonization and hemorrhagic septicaemia are mostly unknown. In this study, we identified two new TonB-dependent receptors, HfpR and BfpR, that are required for adaptation to iron conditions encountered during infection and for virulence in rainbow trout. Transcriptional analyses revealed that their expression is tightly controlled and upregulated under specific iron sources and concentrations. Characterization of deletion mutants showed that they act without redundancy: BfpR is required for optimal growth in the presence of high hemoglobin level, while HfpR confers the capacity to acquire nutrient iron from heme or hemoglobin under iron scarcity. The gene *hfpY*, co-transcribed with *hfpR*, encodes a protein related to the HmuY family. We demonstrated that HfpY binds heme and contributes significantly to host colonization and disease severity. Overall, these results are consistent with a model in which both BfpR and Hfp systems promote heme uptake and respond to distinct signals to adapt iron acquisition to the different stages of pathogenesis. Our findings give insight into the molecular basis of pathogenicity of a serious pathogen belonging to the understudied family *Flavobacteriaceae* and point to the newly identified heme receptors as promising targets for antibacterial development.

## Introduction

Aquaculture is a rapidly growing sector which now represents more than half of the human fish consumption in the world [1]. However, its sustainable development faces the major challenge of controlling infectious diseases caused by poorly known aquatic pathogens. These include several Gram-negative bacteria belonging to the family *Flavobacteriaceae* (phylum *Bacteroidetes*), of which a majority of members are commensals and saprophytes commonly isolated from environmental sources such as soil, sediments and water [2]. Among them, *Flavobacterium psychrophilum* infects salmonids in freshwater environments and is responsible for diseases known as rainbow trout fry syndrome and bacterial cold-water disease which constitute a major chronic health issue for the fish farming industry worldwide [3–8]. *F. psychrophilum* outbreaks cause complex rearing issues and substantial economic losses, impact animal welfare and -due to repeated use of antibiotics- the environment. A better understanding of the pathogenesis and biology of this pathogen is needed to enhance the development of alternative control strategies, such as efficient vaccines, bacteriophages therapy and selection of resistant fish lines [9–12].

Juveniles of rainbow trout (*Oncorhynchus mykiss*) are particularly susceptible. In fry, the disease results in septicaemia leading to mortalities rising up to 70% without appropriate antibiotic treatment. Infected fish present signs of tissue erosion and necrotic lesions, typically in the jaw, the caudal peduncle or the caudal fin [13]. The bacterium is mainly found in the damaged skin and muscular tissues surrounding the lesions, in the lymphoid organs, particularly in the spleen that exhibits hemorrhages and hypertrophy, and harbours a large number of bacteria [14–16]. *F. psychrophilum* cells are highly proteolytic, cytotoxic and possess adhesion abilities [17–20]. The Type IX secretion system (T9SS) is required for pathogenicity and several secreted adhesins, peptidases and other uncharacterized proteins likely play a role in adhesion to host cells, tissue degradation and pathogen spreading [21–23].

Within the host, *F. psychrophilum* cells face a strong inflammatory response [24, 25]. Host colonization relies on the resistance to stresses and the ability to efficiently retrieve nutrients such as amino acids, vitamins and essential metals. In vertebrates, iron is mainly sequestered in iron- or heme-binding proteins preventing toxic effects of free iron and limiting accessibility for bacterial growth. Pathogens have developed sophisticated mechanisms to fulfil their iron needs from various sources [26]. In Gram-negative bacteria, many of the iron acquisition systems rely on TonB-dependent receptors (TBDR) which mediate the active uptake of specific substrates across the outer membrane.

TBDRs are *β*-barrels outer membrane proteins forming pores that are occluded by a globular periplasmic domain (plug) in the absence of substrate. Substrate transport is energized by the inner membrane TonB-ExbBD complex that transduces energy derived from the proton motive force [27]. A large variety of iron uptake TBDRs have been described, involving either the direct binding of host iron- or heme-binding proteins by the receptor, or the indirect substrate capture by high-affinity siderophores or hemophores. Expression of these genes are typically tightly regulated in response to iron availability in order to meet bacterial needs for iron while avoiding toxicity of excessive intracellular iron concentrations [28].

Despite the importance of iron homeostasis, the genes involved in iron uptake remain uncharacterized in *F. psychrophilum*. *In vitro*, growth of the bacterium is strongly inhibited by addition of an iron chelator, but can be restored by the supply of hemoglobin or transferrin, suggesting that the bacterium is able to uptake iron from these compounds [29]. Production of siderophores, hemolysis activity and hemoglobin consumption have been reported [18,21,29]. In addition, the inactivation of *exbD2*, a subunit of the TonB system, results in impaired growth under iron restricted conditions [30], pointing out the probable existence of TonB-dependent iron uptake systems in *F. psychrophilum*.

Here, we took advantage of a large *in vitro* condition-dependent transcriptome dataset [31] to identify iron-regulated TBDR encoding genes. By combining reverse genetics, *in vitro* phenotyping of deletion mutants and experimental infection in rainbow trout, we demonstrate the importance of two heme/hemoglobin receptors for the virulence of *F. psychrophilum*. The first receptor, that we named BfpR, acts under high-level hemoglobin conditions. The second receptor, HfpR, acts under iron-restricted conditions in concert with its cognate hemophore HfpY, a new member of the HmuY family widespread in the phylum *Bacteroidetes*.

## Materials and Methods

### Bacterial strains and growth conditions

The strains used in this study are listed in T1 in *Supporting Information*. This study was performed using THCO2-90, a model strain for *F. psychrophilum* genetic manipulation [17, 32]. *F. psychrophilum* was routinely grown aerobically in tryptone yeast extract salts (TYES) broth [0.4 % (w/v) tryptone, 0.04 % yeast extract, 0.05 % (w/v) MgSO_4_ 7H_2_O, 0.02 % (w/v) CaCl_2_ 2H_2_O, 0.05 % (w/v) D-glucose, pH 7.2] or on TYES agar. When specified, media was supplemented with 5% fetal calf serum (FCS) for optimal growth. Growth experiments in liquid culture were carried out at 200 rpm and 18 °C, as follows: strains were streaked on TYES FCS agar from −80°C freezer stock and incubated 4 days at 18°C, a pre-culture was then prepared by inoculation of one colony into 5 ml of TYES broth and incubation during 36 h. Cultures were performed by inoculation of 10 ml of broth medium at an OD600 of 0.02 with the pre-culture and growth was evaluated by measuring OD600 using a Biochrom WPA CO8000 spectrophotometer. Iron-depleted cultures were performed in 10 ml of TYES broth supplemented with 20 µM ethylenediamine-*N*,*N′*-bis (2-hydroxyphenylacetic acid) (EDDHA) and 1.25 µM hemoglobin, 5 µM hemin or 5 µM FeCl_3_ when specified. Stock solutions were prepared as follows: 2.5 mM lyophilized hemoglobin (Sigma #H2500) in 0.1 M Tris 0.14 M NaCl, 10 mM hemin chloride (Fe-PPIX) (Frontier Scientific) in 50 mM NaOH, 50 mM FeCl_3_ (Merck) in water and 25 mM EDDHA (Sigma) in 50 mM NaOH. Hemoglobin-enriched cultures were performed by inoculation of the pre-culture at an OD600 of 0.005 into 5 ml of TYES broth supplemented with 1% hemoglobin (w/v) prepared from a fresh 10% hemoglobin (Sigma #08449) stock solution. Growth curves were determined by enumeration of colony-forming units (CFU) by plating serial dilutions of the cultures on TYES FCS agar plates. The elastin degradation assay was performed using TYES agar supplemented with 0.75 % (w/v) elastin from bovine neck ligament (Sigma, E1625) as previously described [33]. *E. coli* strains were grown at 37°C in Luria Bertani broth or supplemented with 15 g/l of agar.

### Bioinformatic analysis

Comparison of the gene content between strains was performed on a selection of 22 genomes using the web interface MicroScope [34]. TBDR encoding genes were identified using Interpro predictions of the TonB-dependent receptor family (IPR000531) or the highly conserved structural beta-barrel (IPR036942) and plug (IPR037066) domains. Orthologs were identified using BlastP Bidirectional Best Hit and the MicroScope default parameters (i.e., >80% protein identity, >80% coverage). The presence of truncated TBDR genes was verified by PCR and Sanger sequencing. The structure-assisted multiple sequence alignments of HfpR (THC0290_1814) and BfpR (THC0290_1679) with other heme/hemoglobin receptors and of HfpY (THC0290_1812) with other HmuY-like proteins were constructed with PROMALS3D [35] webserver.

### Plasmid construction and conjugative transfer into F. psychrophilum

Plasmids (T1 in *Supporting Information*) were engineered by the Gibson assembly method using the NEBuilder® HiFi DNA Assembly Master Mix (New England Biolabs) with appropriate primers (T2 in *Supporting Information*). Plasmids were constructed in *E. coli* DH5*α* Z1, verified by PCR and DNA sequencing, transferred to *E. coli* MFD*pir* and subsequently introduced into *F. psychrophilum* OSU THCO2-90 by conjugation as previously described [31]. For selective growth of *E. coli* strains carrying plasmids, transformants were selected with 100 µg/ml of ampicillin. Cultures of *E. coli* MFD*pir* were supplemented with 0.3 mM diaminopimelic acid (Sigma-Aldrich Co.). Selection of *F. psychrophilum* transconjugants was carried out with 50 µg/ml of gentamycin and 10 µg/ml of erythromycin for the pCP*Gm*^r^- and pYT313-derivative plasmids, respectively.

### Construction of transcriptional reporter plasmids and promoter activity monitoring

pCP*Gm*^r^-P_less_-mCh derivative plasmids carrying the promoter region upstream of *hfpR* (240 bp) or *bfpR* (various length: 306, 609 and 950 bp) were constructed as previously described [31]. Briefly, the vector fragment was amplified using primers TRO460/TRO461 with pCP*Gm*^r^-P_less_-mCh as matrix and the promoter region using strain OSU THCO2-90 chromosomal DNA and either primers TRO462/TRO463 resulting in P*_hfpR_*-mCh fusion (pGi48), TRO464/TRO465 resulting in P*_bfpR_1_*-mCh (306 bp, pGi49), TRO789/TRO790 resulting in P*_bfpR_2_*-mCh (609 bp, pGi74) or TRO824/TRO790 resulting in P*_bfpR_3_*-mCh (950 bp, pGi80) fusions. For promoter activity monitoring of P*_hfpR_*-mCh fusion, log-phase cultures (OD600 of 0.4) of the strain carrying the pGi47 plasmid performed in TYES broth supplemented with 20 µg/l gentamycin were distributed into 96-well microtiter plate, cultures were then supplemented with 20 µM of EDDHA and various iron sources (62.5 µM hemoglobin, 250 µM hemin, 5 to 250 µM FeCl_3_) and incubated with continuous shaking at 18°C during 5 h. Cultures of strain carrying the pCP*Gm*^r^-P_less_-mCh plasmid were used as negative control. Promoter activity monitoring of P*_bfp_*-mCh fusions was performed using bacterial colonies grown during 48 h on TYES agar supplemented with 1 % hemoglobin or not. Colonies were scraped from the plates, then resuspended in TYES broth at a final OD600 of 1 for fluorescence measurement. Expression of the *mcherry* gene was monitored using whole-cell fluorescence with a Tecan Microplate Reader (Infinite 200 PRO). Excitation and emission wavelengths were set at 535 nm and 610 nm, respectively. Promoter activity (in arbitrary units) was estimated by dividing fluorescence intensity by the OD600.

### Construction of F. psychrophilum mutants

Three suicide vectors pGi27, pGi29 and pGi38 allowing, respectively, *hfpR*, *hfpY* and *bfpR* deletion were constructed using the pYT313 plasmid [36] as previously described (M1 in *Supporting Information*)[31]. Each resulting plasmid was introduced by conjugation in strain OSU THCO2-90 and colonies selected for double crossing-over were screened by PCR using specific primers (T2 in *Supporting Information*) in order to identify those carrying the deletion genotype. The THC0290_1813 transposition mutant was produced as part of a Tn*4351* mutant library as previously described [23, 37]. Briefly, the pEP4351 suicide plasmid carrying Tn*4351* was introduced by conjugation in strain OSU THCO2-90 and clones were selected on TYES agar supplemented with 10 µg/l erythromycin. Identification of the Tn*4351* interrupted locus was performed by digestion of genomic DNA by HindIII, ligation, inverse PCR of the resulting circular molecules using primers targeting Tn*4351* and subsequent sequencing of the amplicons. Sequences were used to locate the transposon insertion site into the genomic sequence.

### Construction of the F. psychrophilum chromosomal expression platform

Several suicide plasmids allowing chromosomal recombination and gene insertion at a neutral intergenic region (between genomic positions 1121879 and 1124880 for strain OSU THCO2-90) were constructed using pYT313 as backbone (T1 in *Supporting Information*). pGi39 was designed to carry the two DNA fragments upstream and downstream of the insertion site flanking an expression platform composed of a constitutive promoter P*_rpsL_*, a multiple cloning site and an intrinsic terminator, and pGi42 to carry the elastase encoding gene FP0506 under the control of P*_rpsL_* (M2 in *Supporting Information*).

### Complementation of the deletion mutants

The chromosomal ectopic platform was used for reintroduction of deleted genes into the mutant strains. A set of pGi39-derivative plasmids (pGi44, pGi45, pGi46 and pGi55; T1 in *Supporting Information*) was constructed and introduced by conjugation in the respective mutant strains, resulting in the chromosomal insertion of a single copy gene under the control of native promoter at the neutral intergenic region (M2 and F2 in *Supporting Information*).

### Recombinant MBP-HfpY purification and heme binding assays

Recombinant HfpY was purified as a MBP-HfpY fusion protein from the pMBP-HfpY plasmid in *E. coli* Top10. pMBP-HfpY was constructed by gene synthesis (Proteogenix). MBP-HfpY was purified by affinity chromatography on amylose resin as reported previously [38]. Briefly, bacteria were grown in 1 L LB broth to OD600 = 0.6 and expression was induced with 0.5 mM isopropyl 1-thio-β-D-galactopyranoside (IPTG) overnight at RT. Cells were pelleted at 3500 g for 10 min, resuspended in 20 mM Hepes pH 7.5, 300 mM NaCl, containing 1 mM EDTA (binding buffer), and disrupted with glass beads (FastPrep, MP Biomedicals). Cell debris were removed by centrifugation at 18,000 g for 15 min at 4°C. MBP-tagged proteins contained in the supernatant were purified by amylose affinity chromatography (New England Biolabs) following manufacturer’s recommendations: the soluble fraction was mixed with amylose resin and incubated on a spinning wheel at 4°C for 1 h. The resin was then centrifuged, and washed 3 times with binding buffer. Purified proteins were eluted in binding buffer containing 10 mM maltose, and dialyzed against 20 mM Hepes pH 7.5, 300 mM NaCl. Quantity and purity of the eluted fraction were determined with the Lowry Method (Biorad) and by SDS-PAGE followed by Coomassie staining, respectively. Heme binding was studied by UV-visible spectroscopy of MBP-HfpY in the presence of hemin in a microplate spectrophotometer (Spark, Tecan).

### Titration of MBP-HfpY with hemin

Hemin binding affinity and stoichiometry to MBP-HfpY were determined by adding 0.5 to 1 µl increments of a 200 µM hemin solution to cuvettes containing 100 µl test samples of 20 µM MBP-HfpY in 20 mM Hepes pH 7.5, 300 mM NaCl, or reference sample without protein. Spectra were measured from 230 nm to 750 nm in a UV-visible spectrophotometer Tecan (Spark). OD_398nm_ was plotted against hemin concentration and data were fitted to a one-binding site model (with calculated extinction coefficient of bound hemin of ε398 = 40 mM^-1^.cm^-1^) as described [38].

### Total RNA extraction

Bacterial cells from 2 mL cultures were collected by centrifugation for 3 min after addition of ½ volume of frozen killing buffer (20 mM Tris-HCl pH 7.5, 5 mM MgCl_2_, 20 mM NaN_3_) to the culture sample. The cell pellets were frozen in liquid nitrogen and stored at −80 °C. For experiments performed in iron-depleted conditions, RNAs were extracted from pellets harvested at an OD600 of 0.8. For experiments performed in 1% hemoglobin-enriched conditions, RNAs were extracted from pellets harvested when bacterial concentration reached 10^9^ CFU/ml. Experiments were carried out from three independent cultures per biological condition. The ZymoBIOMICS RNA Miniprep Kit (Zymo Research) was used for total RNAs extraction. 5 µg of RNA extracts were treated using DNase I (Qiagen) to remove residual genomic DNA then purified using the ZymoBIOMICS RNA Miniprep Kit. The RNA concentration was measured using a NanoDrop 1000 Spectrophotometer (NanoDrop Technologies, Inc.) and quality was assessed by electrophoresis on denaturing formaldehyde agarose gel.

### Reverse Transcription PCR

Transcriptional structure of the *bfpR* locus was determined by overlapping RT-PCR experiments. Total RNA was extracted from bacterial pellets harvested from cultures performed in TYES 1 % hemoglobin. cDNA synthesis was carried out on 500 ng of DNase-treated RNA using the SuperScript™ II Reverse Transcriptase (Thermo Fisher) with random primers. Negative control reaction was run under similar conditions except that the reverse transcriptase was omitted (no RT). Polycistronic structure was detected by PCR amplification using cDNA as matrix and several primer sets designed to overlap adjacent genes (TRO677/TRO676 between the upstream region of *bfpR* and its coding sequence, TRO674/TRO675 for the junction between THC0290_1678 and *bfpR,* and TRO672/TRO673 for the junction *asd*-THC0290_1678; T2 in *Supporting Information*). Negative and positive controls were run using no RT cDNA reaction and genomic DNA as matrix, respectively, and results were visualized by electrophoresis of PCR products on 1% agarose gel.

### Real Time quantitative PCR

cDNA was generated from 500 ng of total RNA in a thermal cycler (Eppendorf) using the iScript™ Advanced cDNA Synthesis Kit for RT-qPCR (BIO-RAD) as recommended by suppliers. cDNA was diluted 1:25 in water and mixed with iTaq Universal SYBR Green Supermix (BIO-RAD) and forward and reverse primers with final concentration at 0.5 µM each. qPCR was performed using a Realplex Mastercycler (Eppendorf) or CFX system (BIO-RAD) with initial denaturation at 95°C for 3 min followed by 40 cycles of amplification as following: denaturation at 95°C for 5 s, annealing and extension at 60°C for 30 s. For each biological replicate, mean Ct values were calculated based on technical triplicate reactions and then normalized using the geomean of Ct values of two reference genes (*rpsA* and *frr*). Relative quantification of mRNA was expressed as 2^-ΔΔCt^ using TYES as a reference sample.

### Circular Rapid Amplification of cDNA ends (RACE)

RACE was carried out using a general four-step protocol which enables the simultaneous cloning of 5’ and 3’ ends of RNA, as previously described [39]. Two RNA samples extracted from culture of the wild-type strain performed in TYES broth with 20 µM EDDHA or 1% hemoglobin were used respectively for *hfp* or *bfpR* mRNAs RACE experiments. Briefly, 8 µg of DNase-treated RNA samples were treated by 80 units of RppH (New England Biolabs) at 37°C for 1 hour to transform 5’-triphosphate groups carried by primary transcripts into 5’-monophosphate groups, then purified using the ZymoBIOMICS RNA Miniprep Kit. Circular ligation was performed by adding 80 units of T4 RNA ligase 1 (New England Biolabs) to 4 µg of the RppH-treated sample, the mix was incubated at 16°C overnight then heat inactivated. A control sample was prepared following the same protocol except that RppH was omitted in the reaction. First strand cDNA synthesis was performed using the SuperScript™ II Reverse Transcriptase (Thermo Fisher) with 400 ng of the ligation product as the matrix and a specific primer for each 3’-5’ cDNA. 5’-3’ ends junctions were then amplified by PCR using cDNA as the matrix and specific primers (T2 and F4 in *Supporting Information*). PCR products were cloned into the pJET1.2 vector using the CloneJET PCR Cloning Kit (Thermo Fisher) then introduced in *E. coli*. Colonies were screened by PCR prior Sanger sequencing of the insert. Three 3’-5’ cDNAs carrying *hfpR*, *hfpY* or *bfpR* were produced using the gene-specific primers TRO806, TRO811 or TRO814, respectively. Detection of mRNA boundaries was performed as follows: for the *hfpR* cDNA using primers TRO806/TRO807 amplifying *hfpR*-THC0290_1813-*hfpY* mRNAs, for the *hfpY* cDNA using primers TRO812/TRO807 amplifying all *hfpY* mRNAs and for the *bfpR* cDNA using three sets of primers amplifying either all *bfpR* mRNAs (TRO815/TRO816), THC0290_1678-*bfpR* mRNAs (TRO815/TRO825) or *asd*-THC0290_1678-*bfpR* mRNAs (TRO815/TRO827). For *bfpR* RACE experiments, the first PCRs were followed by a nested PCR using TRO817/TRO818, TRO817/TRO826 or TRO817/TRO828, respectively, and the resulting PCR products were cloned into the pJET1.2 vector for Sanger sequencing.

### Rainbow trout experimental infections

Three experimental infections models were used in this study differing in the infection route (intramuscular injection *versus* immersion), the rainbow trout lines or the size of fish (fry *versus* juvenile fish). Fish were reared at 10°C in recycling aquaculture system (RAS) with dechlorinated water, then transferred to BSL2 zone in tanks in flow water (1 renewal per hour) for infection experiments.

For the intramuscular injection model at the fry stage, fish (average body weight of 5 g) of the rainbow trout standard line (Sy*Aut) selected by INRAE were used. Six groups of 10 fish were anesthetized and challenged with the wild-type strain at different doses, as follows: 6 bacterial suspensions were prepared by ten-fold serial dilution of a culture at an OD600 of 1 performed in TYES broth, and 50 µl of each suspension was injected per fish, corresponding to 50 CFU to 5 x 10^6^ CFU. LD50 of the wild-type was estimated at 5 x 10^6^ CFU. A similar procedure was used for the 8 deletion mutants and complemented strains except that only 3 doses were tested, corresponding to 5 x 10^4^ CFU, 5 x 10^5^ CFU and 5 x 10^6^ CFU per fish. *F. psychrophilum* bacterial counts were determined by plating serial dilutions of bacterial suspensions on TYES agar supplemented with FCS. After injection, fish were maintained in flow-through water at 10°C and mortalities were recorded twice a day for two weeks. The ability of bacterial strains to colonize rainbow trout at the fry stage (average body weight of 2 g) was compared using the bath infection model on the rainbow trout homozygous line A36, which is highly susceptible to *F. psychrophilum* infection [10,23,40]. Groups of 40 fish in duplicates were used for each strain. Bacterial cultures were prepared as follows: two independent cultures of the wild-type, deletion and complemented mutant strains were grown in TYES broth until late-exponential phase (OD600 of 1). The bacterial cultures were directly diluted (200-fold) into 10 l of aquarium water. Bacteria were maintained in contact with fish for 24 h by stopping the water flow and were subsequently removed by restoring the flow. During the experiment, water was maintained at 10°C under continuous aeration and physical parameters (NH_4_^+^, NO_2_^-^) were monitored. Sterile TYES broth was used for the control group. *F. psychrophilum* bacterial counts were determined at the beginning and at the end of the immersion challenge by plating serial dilutions of water sample on TYES FCS agar. Five fish from each tank (n = 10 per strain) were randomly chosen and euthanatized (tricaine, 300 mg/l) at 6 h post-infection to sample gills and spleen. Organs were mechanically disrupted in tubes containing 400 µl of peptone (10 g/l water) and 1 mm ceramic beads (Mineralex SAS, Lyon, France) using a FastPrep-24™ 5G instrument at 6.0 m s-1 for two cycles of 20 s (MP Biomedicals), and bacterial loads were determined by plating serial dilutions of lysates on TYES FCS agar.

Virulence of the *bfpR* deletion mutant and the wild-type strain was evaluated by intramuscular injection at the juvenile stage (average body weight of 65 ± 9 g) using the rainbow trout line A36. Bacterial suspension was prepared by 10-fold dilution in sterile TYES of a culture at an OD600 of 1. For each strain, a group of 24 fish were anesthetized and inoculated with 100 µl of bacterial suspension then maintained in flow-through water at 10°C into two compartments: the first subset (n = 10) was used for survival estimation by recording mortalities during 2 weeks and the second subset (n = 12) for sampling of spleen, head kidney and blood at 4 days post-infection and bacterial loads determination. Following anaesthesia, blood (150 µL per fish) was collected from the caudal vein using syringes containing heparin (500 units/ml) and the bacteraemia was determined by plating fresh blood and serial dilutions on TYES FCS agar plates. Infectious doses and bacterial loads in organs were determined as described above.

Statistical differences of bacterial loads between groups were analysed using the Mann-Whitney test, and survival curves were determined using the Kaplan-Meier method and compared using the Mantel-Cox log-rank test with GraphPad Prism 8.2.0 (GraphPad Software, San Diego, CA, USA).

### Ethics statements

The infection challenges were conducted at the INRAE-IERP Fish facilities of Jouy-en-Josas (building agreement n°C78-720), performed in accordance with the European Directive 2010/2063/UE regarding animal experiments, and approved by the institutional review ethics committee, COMETHEA, of the INRAE Center in Jouy-en-Josas, France. Authorizations were approved by the Direction of the Veterinary Services of Versailles (authorization number #19210-2015100215242446).

## Results

### Identification of two TonB-dependent receptors with potential function in heme utilization

The distribution of TBDRs was analysed in a collection of 22 genomes of *F. psychrophilum* isolates representative of a variety of host species, geographical origins and clonal complexes [41]. A total of 33 TBDR-encoding genes were retrieved. With an average number of 25 per genome, most (21) of these predicted TBDRs were conserved in all isolates, though 3 were pseudogenes in one isolate (Figure 1). Only a few TBDRs have predicted substrates, among which are the cobalamin transporter BtuB and 2 putative receptors homologous to the ferric hydroxamate receptor FhuA and the ferric citrate transporter FecA [42]. Among TBDR genes found only in a subset of the isolates, 3 may be associated to a particular host fish species. The first one, THC0290_1129 in strain OSU THCO2-90, is only present in the genomes of CC-ST9 isolates infecting coho salmon. The 2 others, FP2456 and FP1882 in strain JIP 02/86, are present in all isolates belonging to the clonal complexes associated with rainbow trout (CC-ST10 and CC-ST90), but truncated and absent in the other genomes, respectively.

**Figure 1.**
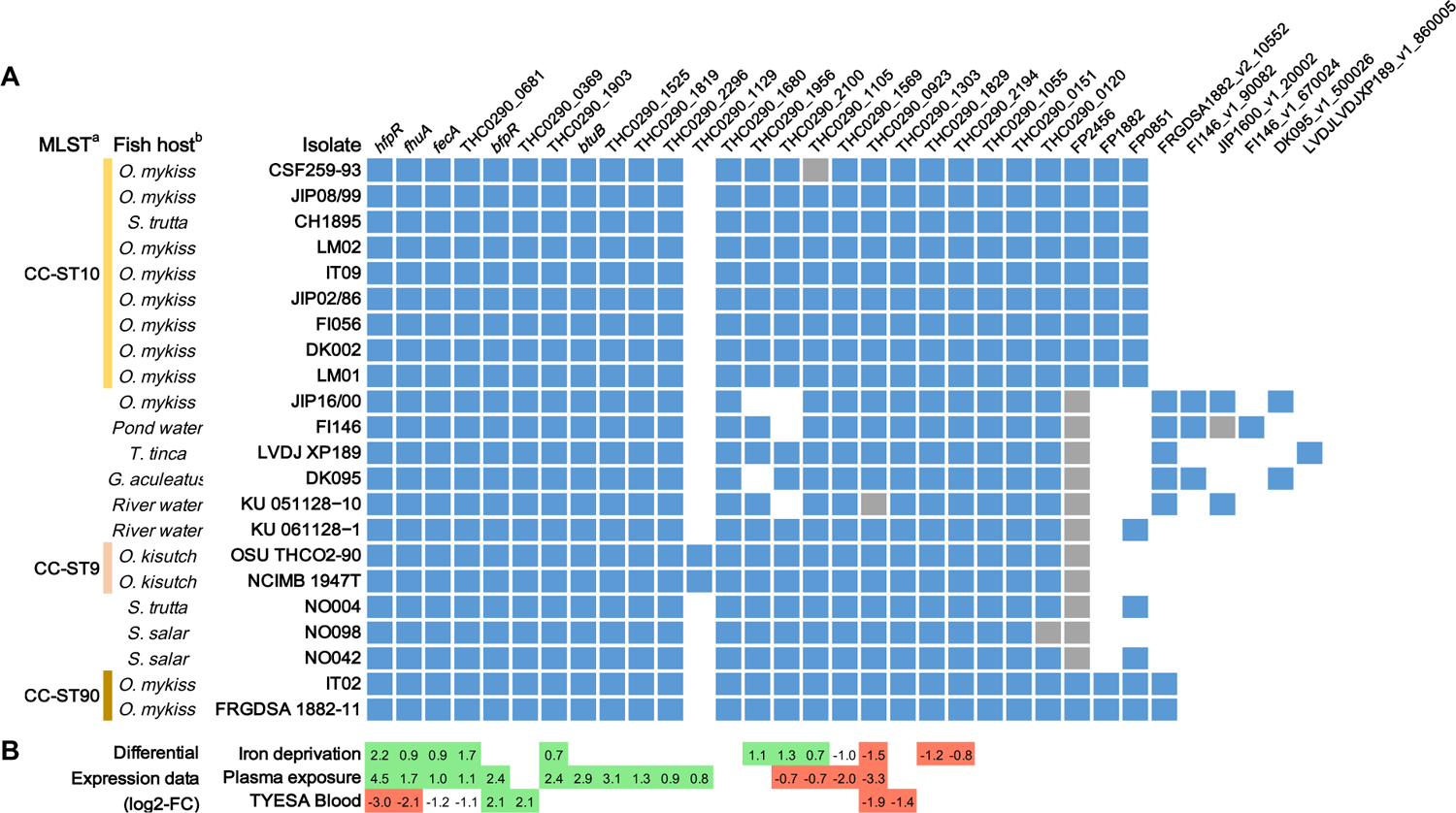
Conservation and gene expression of the 24 TonB-dependent receptors encoded in the genome of strain OSU THCO2-90. (A) *In silico* prediction of TBDR encoding genes. The list of TBDR encoding genes was retrieved by *in silico* prediction of the plug and ß-barrel domains. Conservation in a selection of 22 isolates: gene present (blue), truncated (grey), absent (white). ^(a)^MLST and genomic data are from Duchaud *et al* [41]. CC-ST: clonal complex-sequence type. ^(b)^Fish host: *Oncorhynchus mykiss*, *Salmo trutta*, *Tinca tinca*, *Gasterosteus aculeatus*, *Plecoglossus altivelis*, *Oncorhynchus kitsutch*, *Salmo salar.* (B) TBDR-encoding genes significantly upregulated (green) and down-regulated (red) in strain OSU THCO2-90 in iron-limited condition (TYES + 25 µM 2,2’-dipyridyl), under rainbow trout plasma exposure or in the presence of blood (TYES agar supplemented with 10% horse blood). Differential expression is expressed as log2-fold change values retrieved from Guérin *et al* [31](https://fpeb.migale.inrae.fr).

In order to further identify, among these TBDRs, those that are involved in iron acquisition, we searched for genes whose expression profile indicated specific regulation under blood supplementation, fish plasma exposure or conditions of iron limitation in the condition-dependent transcriptome dataset previously established for strain OSU THCO2-90 [31]. Two TBDR genes were up-regulated in the presence of blood: (i) THC0290_0369 without predicted substrates, and (ii) THC0290_1679, a member of the heme/hemoglobin receptor family (TIGR01785), hereafter named *bfpR* (for <u>b</u>lood-induced *<u>F</u>lavobacterium <u>p</u>sychrophilum* <u>r</u>eceptor). *bfpR* was also overexpressed in the presence of fish plasma but was not modulated by iron limitation. Four TBDR-encoding genes were up-regulated in cells submitted to low-iron conditions, but down-regulated when blood was supplied (Figure 1 and F1 in *Supporting Information*): (i-ii) FhuA and FecA siderophore receptors homologs; (iii) THC0290_0681 with unknown substrate; and (iv) THC0290_1814, the most highly deregulated gene, hereafter named *hfpR* (for <u>h</u>eme *<u>F</u>. <u>p</u>sychrophilum* <u>r</u>eceptor). *hfpR* is conserved in synteny with a gene encoding a HmuY-like hemophore protein (hereafter named *hfpY*), suggesting functional similarity with the heme utilization system Hmu characterized in other *Bacteroidetes* [43–46].

Here, we further characterized the BfpR and Hfp systems predicted to transport heme. Expression profiles suggesting a distinct regulation of the corresponding genes (F1 in *Supporting Information*), we thus examined their transcriptional structure and regulation (see below). To analyse their respective role in bacterial growth, we also constructed the single and double deletion mutants *ΔbfpR*, *ΔhfpR*, *ΔhfpY* and *ΔhfpRΔbfpR*, by allelic replacement in *F. psychrophilum* OSU THCO2-90 (T1 in *Supporting Information*). The ectopic complementation of these mutants was performed by inserting a single gene copy under the control of the native promoter at a different chromosomal position than the endogenous deleted gene copy. For *ΔhfpRΔbfpR*, the 2 genes were inserted arranged in tandem. For this purpose, a new plasmid was designed allowing chromosomal gene insertion at a neutral site conserved within the *F. psychrophilum* species and the efficacy of this expression platform was validated using reporter genes (F2 and F3 in *Supporting Information*). The growth of the mutants was analysed *in vitro* under different conditions of iron and *in vivo* in the natural host.

### High levels of hemoglobin promote the accumulation of monocistronic bfpR mRNA

BfpR is encoded downstream of the aspartate-semialdehyde dehydrogenase (*asd*) and a hypothetical protein (THC0290_1678; Figure 2A). Structure of the *bfpR* transcript is uncertain: expression level across biological conditions differs between *bfpR, asd* and THC0290_1678, and the increase of mRNA level in colonies grown on blood-supplemented TYES agar occurs upstream *bfpR*, suggesting that *bfpR* is transcribed as a monocistronic mRNA. However, no transcription start site (TSS) was detected in the corresponding region (Figure 2A)[31].

**Figure 2.**
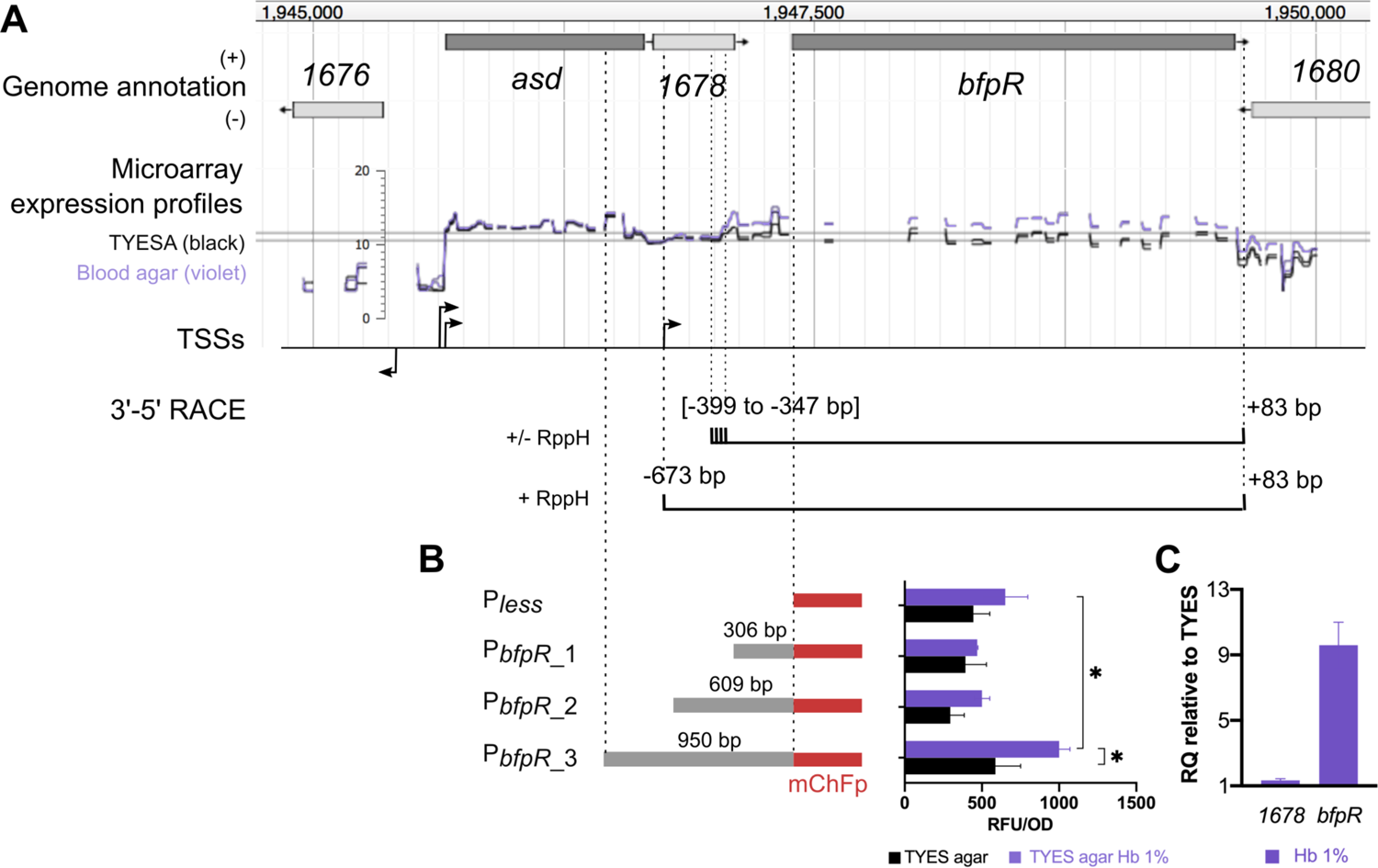
Transcriptional organization of the *bfpR* locus and its regulation by hemoglobin concentration. (A) Genomic view of the *bfpR* locus, microarray expression profiles and putative transcription start sites (TSS) from Guérin *et al* [31]. Expression profiles (log2-expression signal) show the upregulation of *bfpR* in the presence of blood: bacterial colonies grown on TYES agar (TYESA) covered by 10% defibrinated horse blood (violet) or not (black). 3’-5’ RACE: Schematic representation of 5’ and 3’ ends identified by circular 3’-5’ RACE experiments. Positions of the 5’ and 3’ ends are relative to the start and stop positions of *bfpR* CDS, respectively. (B) Assessment of transcriptional initiation upstream of *bfpR* using transcriptional fusions. Promoter activity was measured using whole-cell fluorescence of *F. psychrophilum* cells carrying the reporter plasmid pCPGm^r^-P*less*-mCh with various DNA fragments (Pless: empty plasmid; P*bfpR*_1: 306 bp, P*bfpR*_2: 609 bp, P*bfpR*_3: 950 bp). Bacterial colonies were grown on TYES agar supplemented with 1% hemoglobin (equivalent of 620 µM heme molecules) or not. Values represent the mean and standard deviation of three independent experiments. (*) indicates significant difference identified in a two-way ANOVA analysis (Bonferroni adjusted p-value < 0.05). (C) RT-qPCR measurement of *bfpR* expression. mRNA level was quantified using RNA extracted from cultures performed in TYES broth supplemented with 1% hemoglobin (Hb 1%) or not. Ct values of genes were normalized using the geomean of two reference genes (*rpsA* and *frr)*. RQ: Relative quantification of mRNA was expressed as 2^-ΔΔCt^ using TYES as reference sample. Values are the mean and standard deviation of three independent experiments.

Several methods were combined to define the structure of the *bfpR* transcript. First, a putative polycistronic mRNA encompassing *asd*, THC0290_1678 and *bfpR* was predicted by RT-PCR using overlapping primers. Circular 3’-5’ RACE experiments were then run in an attempt to identify the most abundant *bfpR* transcript and to distinguish between primary and processed RNAs (Figure 2A; F4 in *Supporting Information*). *bfpR* amplification from a primer within *asd* was unsuccessful, attesting of the very low quantity of the polycistronic mRNA predicted by overlapping RT-PCR. In contrast, cDNA was successfully amplified using primers within *bfpR*. The majority of transcripts ended at +83 bp relative to the *bfpR* stop codon, while no discrete 5’-end position was identified. Instead, most transcripts started within a ∼50 bp-long region spanning from −399 to −347 bp relative to the *bfpR* start codon, which coincides with the upshift of expression signal detected by microarrays in blood supply conditions. By using a primer within THC0290_1678, transcripts were detected by nested PCR only and started mainly at −673 (±1) bp relative to *bfpR* start codon. This 5’ end is 22 bp downstream the start position of THC0290_1678. However, the location of translation initiation is difficult to predict in *Bacteroidetes* [47] and another ATG codon located 38 bp downstream the 5’ end may be the correct start. Various fragments (306, 609 and 950 bp) of the upstream region of *bfpR* were cloned into a transcriptional reporter plasmid and a significant promoter activity was detected only for the one carrying the longest fragment when colonies were grown in TYES agar supplemented with 1% hemoglobin (Figure 2B). These results indicate that the majority of *bfpR* mRNAs are monocistronic and may originate from processing of a longer transcript starting 673 bp upstream of *bfpR*.

We therefore compared the effect of hemoglobin supply on *bfpR* mRNA level by RT-qPCR assays (Figure 2C). Addition of large amounts of hemoglobin in TYES broth led to 10-fold increase of *bfpR* mRNA level. This increase was not observed for the upstream gene THC0290_1678. Altogether, we concluded that a primary transcript encompassing THC0290_1678 and *bfpR* is synthetized then probably subject to maturation, resulting in the *bfpR* mRNA that accumulates in presence of large amounts of hemoglobin.

### BfpR contributes to growth in the presence of high amounts of hemoglobin

On the basis of its expression profile, we analyzed the role of BfpR on the growth of *F. psychrophilum* under high-hemoglobin condition. The wild-type strain exhibited a slight delay in the presence of 1% hemoglobin (equivalent of 620 µM heme), but final viable cell counts were higher than in TYES broth (F5 in *Supporting Information*), indicating that hemoglobin availability can enhance bacterial growth. In these conditions, the growth rate of *ΔbfpR* was reduced compared to that of the wild-type and partially restored in the complemented *ΔbfpR* strain (Figure 3). RT-qPCR showed that ectopic expression of *bfpR* in this complemented *ΔbfpR* strain resulted in reduced mRNA level compared to the wild-type (F6 in *Supporting Information*), indicating that adequate regulation of *bfpR* expression is important for growth in these conditions. Growth curves of *ΔhfpRΔbfpR* and *ΔbfpR* were similar. These results showed that among the two TBDRs, only BfpR is required for optimal growth in the presence of high amounts of hemoglobin.

**Figure 3.**
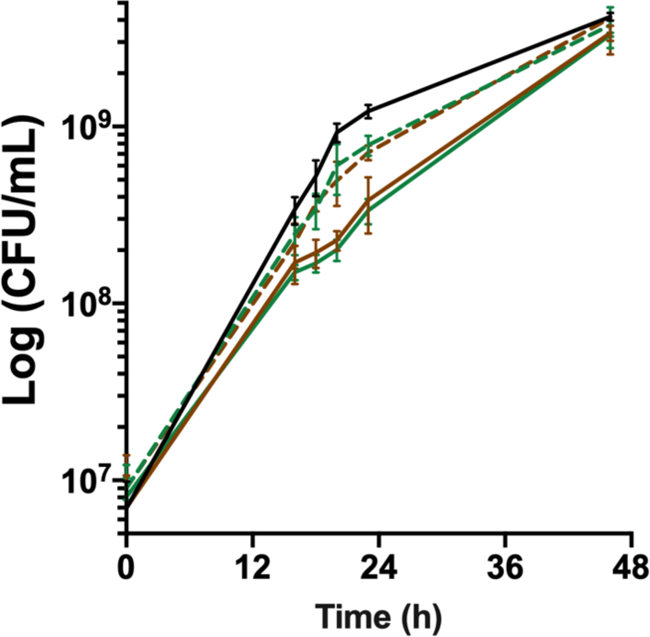
*bfpR* is required for *in vitro* growth under high hemoglobin concentration. Growth under high hemoglobin concentration determined by viable cell measurements (CFU/mL) of cultures in TYES broth supplemented with 1% hemoglobin. Plain lines: wild-type (black) and deletion mutants (*ΔbfpR*: green; *ΔhfpRΔbfpR*: brown), dashed lines: ectopic complemented mutants. Values represent the mean and standard deviation of four independent experiments.

### The hfp operon is transcribed in response to iron scarcity

The Hmu systems characterized so far in *Porphyromonas gingivalis* and other *Bacteroidetes* are transcribed in a polycistronic transcript encoding at least 1 HmuY-like hemophore, 1 TBDR and 4 uncharacterized proteins (a putative CobN/magnesium chelatase and 3 putative membrane proteins) [45, 46]. In *F. psychrophilum*, only the genes encoding a TBDR (*hfpR*) and the HmuY-like protein (*hfpY*) are present. They are located within a 19 kb region containing other iron-responsive transcripts encoding proteins that have a putative role in iron assimilation or transmembrane electron transfer (T3 in *Supporting Information*)[31, 48]. The upregulated polycistronic mRNA deduced from transcriptomic data is bound by a putative TSS and a predicted intrinsic terminator and encodes, from 5’ to 3’: (i) HfpR; (ii) THC0290_1813, a predicted 18 kDa inner membrane protein of unknown function belonging to the DoxX family (IPR032808); and (iii) HfpY (Figure 4A). Expression signal measured by high-density microarrays was higher in the *hfpY* part of the operon for all biological conditions suggesting that a monocistronic *hfpY* mRNA may also be produced (F7 in *Supporting Information*). We confirmed this transcriptional structure by 3’-5’ RACE experiments: two 5’-ends and one unique 3’-end were identified (Figure 4A; F4B in *Supporting Information*). RppH enzymatic treatment increased cDNA amplification for the polycistronic form only indicating a 5’-triphosphorylated RNA. These results confirmed the TSS position and the existence of 2 mRNAs, while suggesting that the *hfpY* mRNA originates from a post-transcriptional maturation event.

**Figure 4.**
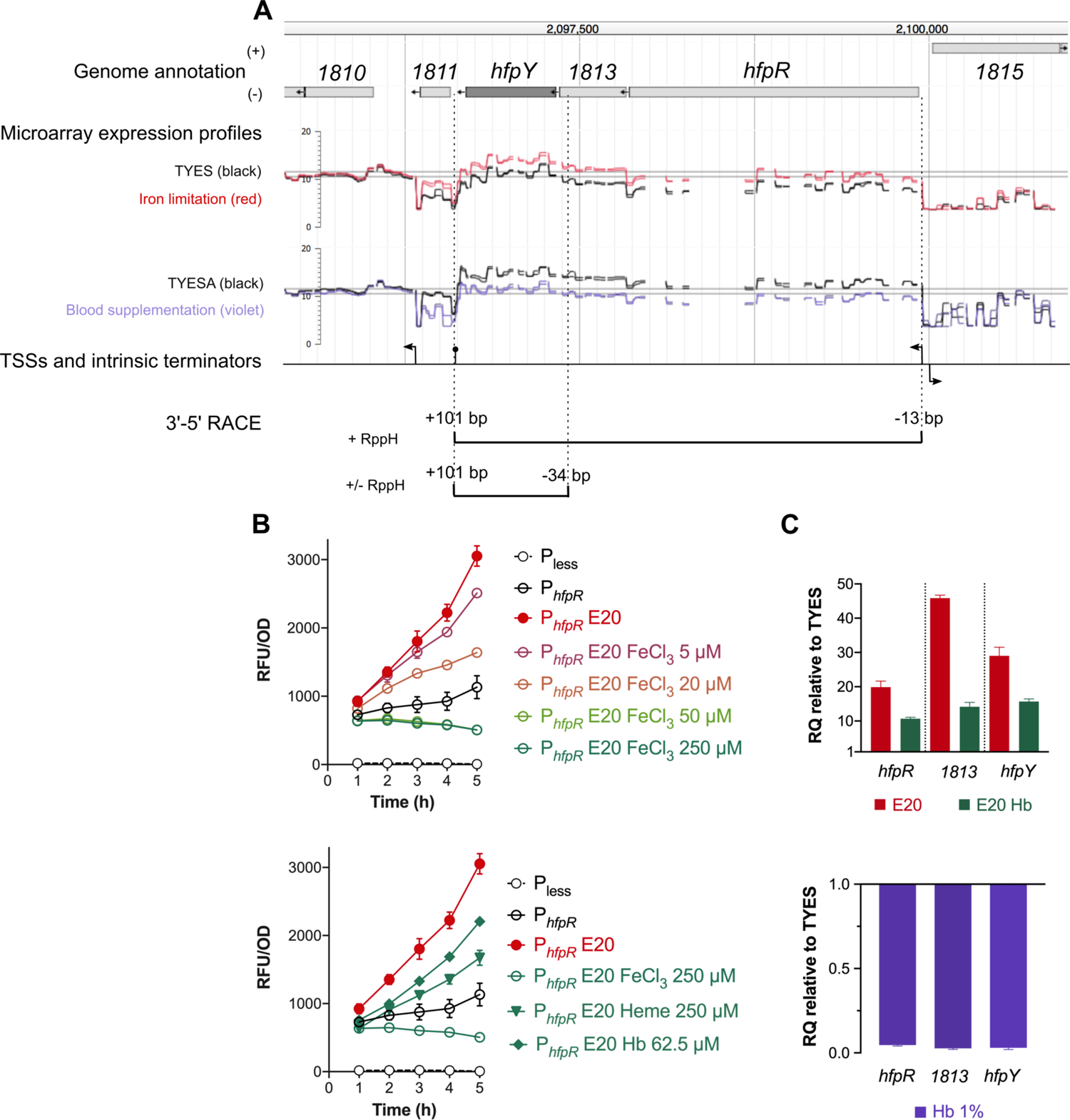
Transcriptional organization of the *hfp* locus and its regulation by iron availability. (A) Genomic view of the *hfpR*-THC0290_1813-*hfpY* locus, microarray expression profiles and putative TSSs from Guérin *et al* [31] are shown (see legend of Figure 2). For iron limitation, cells were grown in TYES broth supplemented with 25 µM 2,2’-dipyridyl (red) or not (black); for blood supplementation, cells were from colonies grown on TYES agar (TYESA) covered by 10% defibrinated horse blood (violet) or not (black). 3’-5’ RACE: Schematic representation of 5’ and 3’ ends identified by circular 3’-5’ RACE experiments. Positions of the two 5’ ends and of the 3’ end are relative respectively to start positions of *hfpR*, *hfpY*, and stop position of *hfpY*. (B) Transcriptional regulation of the *hfpR* promoter by iron availability. Promoter activity was measured using whole-cell fluorescence of the wild-type strain carrying the reporter plasmid with 240 bp DNA upstream of *hfpR* (P*hfpR*). Bacteria were grown in iron depleted TYES broth (E20, 20 µM EDDHA) supplemented with a range of FeCl3 (upper panel) or with different iron sources (lower panel) as indicated. Cultures were inoculated at OD600 0.4 and fluorescence was measured during 5 h as described in Materials and Methods. Strain carrying the empty plasmid (Pless) was used as control. Values represent the mean and standard deviation of three independent experiments. (C) *hfpR*, *1813* and *hfpY* mRNA levels correlate with iron availability. RT-qPCR measurements were performed under iron depletion (upper panel) or high concentration of hemoglobin (lower panel) as described in Figure 2 using the wild-type strain grown in TYES broth, iron-depleted TYES (E20) supplemented with 1.25 µM hemoglobin (E20 Hb) or in TYES supplemented with 1% hemoglobin (Hb 1%).

Expression of the *hfp* locus was then monitored under iron-depleted and -replete conditions using a transcriptional reporter (P*_hfpR_*-mCh) and RT-qPCR assays. Addition of the iron-chelator ethylenediamine-*N*,*N′*-bis (2-hydroxyphenylacetic acid) (EDDHA) in TYES broth resulted in a strong increase of promoter activity, which was countered by FeCl_3_ supply in a dose-dependent manner (Figure 4B). Promoter activity was also limited by heme or hemoglobin supply, although with a lower efficiency than when FeCl_3_ was used (Figure 4B). RT-qPCR confirmed up-regulation of the 3 genes composing the *hfp* operon under iron limitation and the effect of hemoglobin supply (Figure 4C). Moreover, a 5-fold down-regulation was measured in the presence of high level of hemoglobin in iron-replete TYES broth (Figure 4C), which is consistent with the down-regulation previously observed by microarrays in the presence of blood [31].

We concluded from these results that 2 mRNAs are produced at the *hfp* locus: a primary transcript encompassing *hfpR*, THC0290_1813 and *hfpY,* and a matured mRNA carrying *hfpY* only, resulting in more abundant mRNA for *hfpY* than for *hfpR*. The expression level of these 2 RNAs is tightly controlled in response to iron availability through the regulation of the *hfpR* promoter which is activated by iron scarcity and downregulated in presence of iron sources.

### HfpR and HfpY are required for heme utilization under iron scarcity

The effect of iron limitation on growth was first analysed on the wild-type strain. Addition of EDDHA, a chelator of ferric iron Fe(III), to TYES broth resulted in a strong growth inhibition, confirming that iron is an essential element for *F. psychrophilum*. Growth was restored by the supply of hemoglobin, hemin or FeCl_3_, indicating that the bacterium is able to scavenge these different iron sources from the environment (Figure 5).

**Figure 5.**
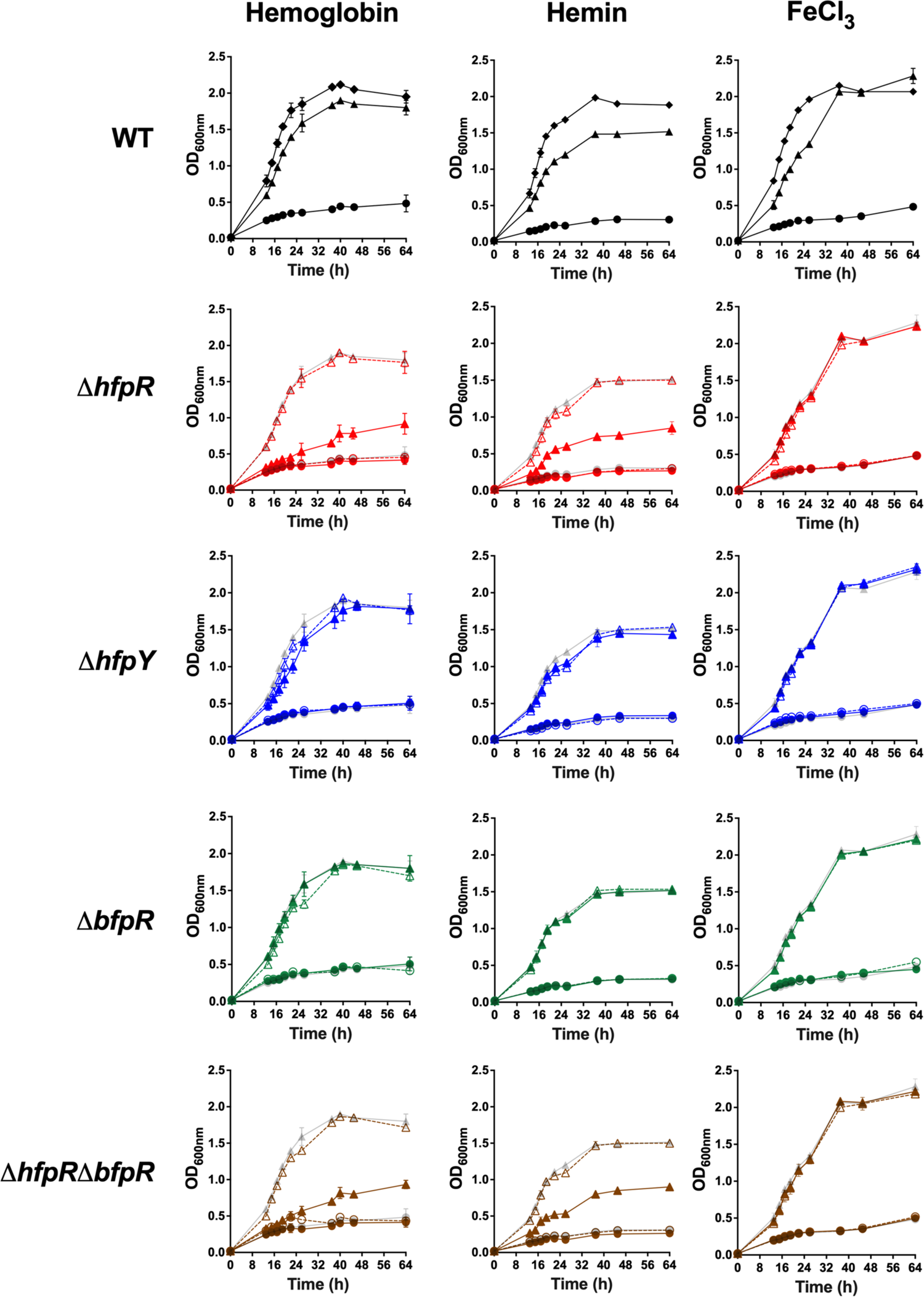
*hfpR* is required for *in vitro* growth under iron scarcity. Growth measured by optical density at 600nm of the wild-type, mutants and complemented strains (rows) under various conditions of iron availability (columns). Cultures were performed in TYES broth containing the iron chelator EDDHA 20 µM only (circle) or containing EDDHA and an iron source (triangle): 1.25 µM hemoglobin (left); 5 µM hemin (center) and 5 µM FeCl3 (right). Growth curves in TYES broth were similar for all strains and are plotted for the WT panel only (diamond). Plain lines: wild-type and deletion mutants (*ΔhfpR*: red, *ΔhfpY*: blue, *ΔbfpR*: green, *ΔhfpRΔbfpR*: brown), dashed lines: ectopic complemented mutants. For mutants, growth curves of wild-type are in light grey for comparison purpose. Values represent the mean and standard deviation of three independent experiments. Growth with different iron sources were from independent experiments.

We then analysed the role of HfpR and HfpY by comparing the growth of deletion mutants and respective complemented strains under iron scarcity (Figure 5). Supplementation by FeCl_3_ fully restored the growth of all strains (Figure 5). In contrast, growth inhibition was only partially restored by hemoglobin and hemin supplementation in mutant *ΔhfpR* which still exhibited a severe growth delay compared to complemented *ΔhfpR* and wild-type strains. *ΔhfpY* was by far less affected but its growth rate still significantly reduced compared to the wild-type in hemoglobin supplemented medium containing EDDHA, and this growth defect was partially restored in the complemented *ΔhfpY* strain. In contrast, no growth defect was observed in *ΔhfpY* when hemin was supplied instead of hemoglobin (Figure 5). Partial growth restoration in the ectopic complemented *ΔhfpY* strain is likely due to its reduced *hfpY* mRNA level compared to the wild-type as measured by RT-qPCR (F6 in *Supporting Information*).

We also evaluated the growth on hemoglobin of a transposition mutant inactivated in the unknown function gene lying between *hfpR* and *hfpY* (THC0290_1813); no difference was observed compared to the wild-type, indicating that this gene is dispensable for heme utilization under iron-restricted conditions (F8 in *Supporting Information*).

Finally, the behaviour of *ΔbfpR* was analysed. In contrast to *ΔhfpR* and *ΔhfpY*, no growth defect was observed under iron limitation when hemoglobin, hemin or FeCl_3_ was supplied, and growth of *ΔhfpRΔbfpR* was indistinguishable from that of *ΔhfpR* (Figure 5). We concluded that HfpR and HfpY are important for utilization of heme compounds under iron scarcity while BfpR is dispensable, and that there is no functional compensation by other genes for the lack of HfpR under these conditions.

### F. psychrophilum HfpY is a heme-binding protein

HfpY exhibits only low level of similarity with other HmuY-like hemophores (below 26% identity in amino-acids sequence; F9 in *Supporting Information*); we thus investigated the ability of HfpY to bind heme *in vitro*. Recombinant HfpY was purified as a fusion to the maltose-binding protein (MBP) from *E. coli*. Spectroscopic monitoring showed that while the maximum absorbance of free hemin was located between 361 and 384 nm, addition of purified MBP-HfpY resulted in a Soret peak at *λ*_max_ 398 nm typical of heme coordination to protein (Figure 6A). Q bands were also visible at 503, 530 and 623 nm in the difference spectra. Heme titration assays indicated a stoichiometry of 1:1 for the holoprotein which is similar to the other characterized HmuY-like hemophores (Figure 6B). Based on these results, we concluded that HfpY of *F. psychrophilum* is a heme-binding protein.

**Figure 6.**
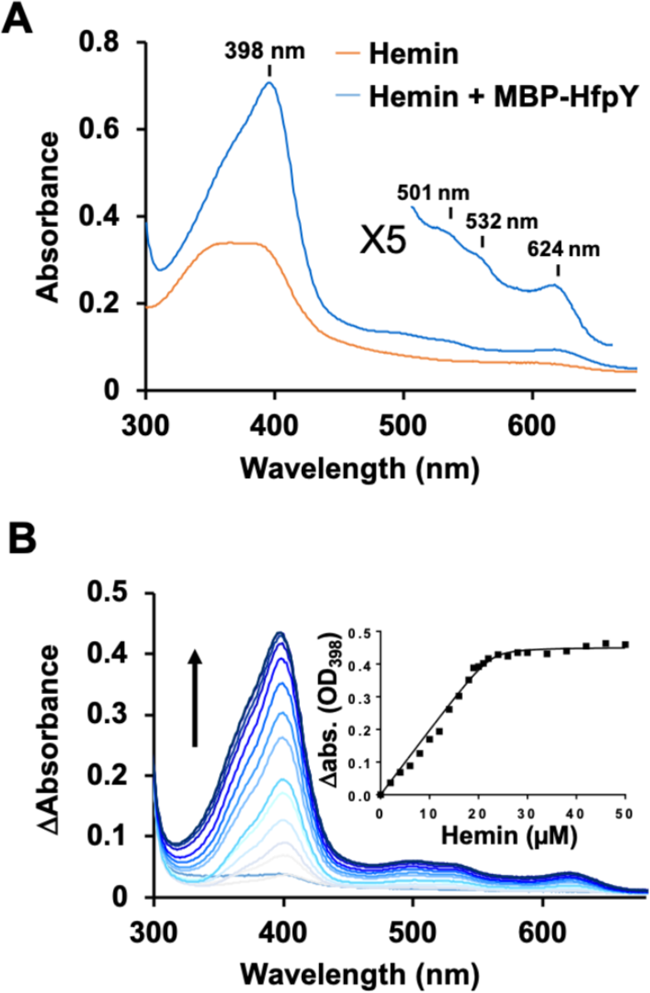
HfpY is a heme-binding protein. (A) UV-visible absorption spectra of MBP-HfpY complexed with hemin. 40 µM MBP-HfpY was purified from *E. coli* and mixed with equimolar concentration of hemin. UV-visible spectra (in 200 µl) of the complex and same concentration of hemin were obtained in a microplate spectrophotometer (Spark; Tecan). (Inset) 5X Magnification of the 500- to 700 nm region. Results are representative of three independent experiments. (B) Titration of MBP-HfpY with hemin followed by absorbance at 398 nm. 20 µM of MBP-HfpY were mixed with increasing concentrations of hemin as indicated. For each spectrum, the absorbance of the same concentration of hemin was subtracted and the difference absorption spectra are shown, from light to dark blue color as a function of hemin concentration. The absorption difference at 398 nm was plotted against hemin concentration (Inset). The curve is representative of 3 independent experiments and was fitted using the nonlinear regression function of GraphPad Prism 4 software, which determined that the stoichiometry of the HfpY-hemin complex was 1:1.

### The two TBDRs are required for virulence in rainbow trout

Wild-type, deletion mutants and complemented strains were analysed for virulence in rainbow trout using three experimental infection models that differ by the infection route and fish size. The roles of *hfpR, hfpY* and *bfpR* were first evaluated by measuring bacterial loads in organs after immersion challenge in fry (2 g), a model that mimics the natural route of infection. At 6 h post-infection, *F. psychrophilum* was detected in the gill of fish from all infected groups at a similar concentration (average of 9 x 10^3^ CFU/gill). In contrast, colonization of spleen was significantly lower for groups infected with *ΔhfpR, ΔhfpY* or *ΔhfpRΔbfpR* relative to the wild-type (Figure 7A). No significant difference was observed between bacterial loads of fish infected with *ΔbfpR* or the wild-type, though values were more scattered with *ΔbfpR*. Bacterial colonization of the spleen was fully restored by complementation of the *ΔhfpRΔbfpR* double mutant compared to the wild-type, but only partially restored in the complemented *ΔhfpY* strain.

**Figure 7.**
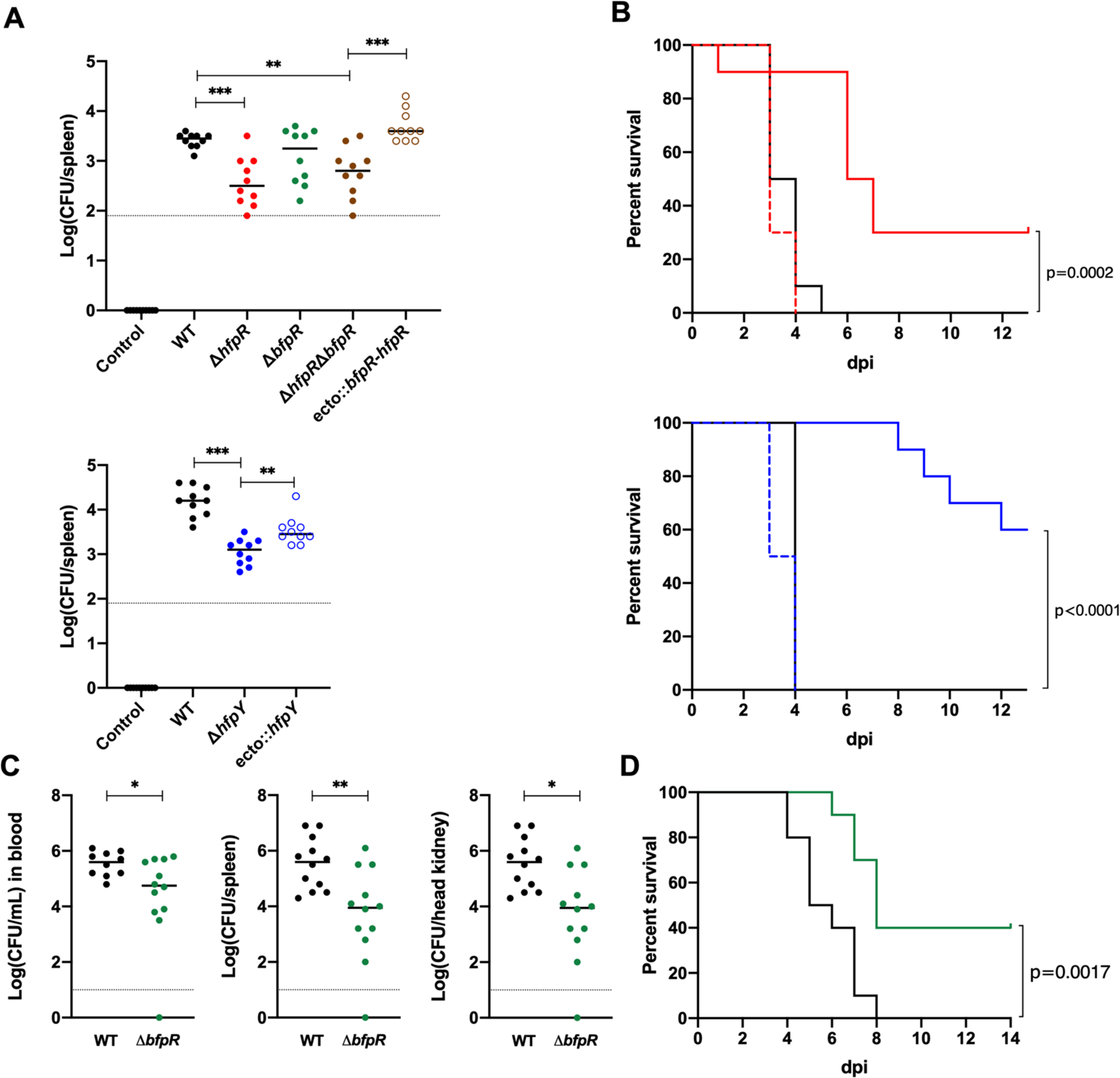
Evaluation of the virulence of wild-type and deletion mutant strains in rainbow trout. (A) Bacterial loads in the spleen of rainbow trout fry after immersion challenge (A36 line, 2 g average weight). Each tank was inoculated at a bacterial concentration of 1 (±0.3) x 10^7^ CFU/mL and fish were maintained in contact with bacteria for 24 h at 10°C under optimal oxygenation conditions. At the end of the challenge, bacterial concentration in water was 2 (±0.7) x 10^7^ CFU/mL. Ten fish from each group were sacrificed at 6 h post-infection and bacterial loads in the spleen were determined. Horizontal dashed line indicates the detection threshold. Values are significantly lower in the spleen of groups infected with the *ΔhfpR* and *ΔhfpY* mutants compared to the wild-type and complemented strains (Mann-Whitney: *, 0.033; **, 0.002; ***, <0.001). (B) Kaplan-Meier survival curves of rainbow trout fry (Sy*Aut line, 5 g average weight) after intramuscular injection. Each group (n=10) was challenged at a dose corresponding to 10 x LD50 of the wild-type strain. Upper panel: 3 x 10^6^ CFU for the wild-type (black), 2 x 10^6^ CFU for *ΔhfpR* (red plain line) and ecto::*hfpR* (red dashed line); lower panel: 6 x 10^6^ CFU for the wild-type, 7 x 10^6^ CFU for *ΔhfpY* (blue plain line) and ecto::*hfpY* (blue dashed line). The survival curves for fish inoculated with either the wild-type or the ectopic complemented mutants were significantly different from those inoculated with the *ΔhfpR* or *ΔhfpY* mutants (p-value of the Mantel-Cox log-rank test is indicated). (C-D) Experimental infection of juvenile rainbow trout (A36 line, 65 g average weight) after intramuscular injection. Fish were inoculated with 1 x 10^7^ CFU for the wild-type (black) or *ΔbfpR* mutant (green) strains; a subset (n = 12) was used for bacterial loads determination in blood, spleen and head kidney at 4 days post-infection (C) and another subset (n = 12) for survival estimation (D). Significant differences were observed between both the Kaplan-Meier survival curves (Mantel-Cox log-rank test p-value) and the bacterial loads (Mann-Whitney test).

Survival curves were then compared in fry (mean weight, 5 g) after intramuscular injection to quantity the virulence. At an infection dose corresponding to 10 LD50 of the wild-type strain, 100% of fish died rapidly in groups infected with the wild-type or complemented strains whereas mortality was delayed and survival was significantly increased for fish infected with either *ΔhfpR* or *ΔhfpY* (Figure 7B). A similar difference between strains was also observed using a lower infectious dose (F10 in *Supporting Information*). Using this infection model, the survival curves of fish inoculated with *ΔbfpR* or the wild-type were similar. In addition, the levels of mortality observed with *ΔhfpRΔbfpR* and *ΔhfpR* were similar (F10 in *Supporting Information*).

Since massive hemolysis may not be encountered in the fry infection model, we also investigated the role of BfpR in virulence on larger fish. Two groups of juvenile rainbow trout (mean weight, 65 g) were infected by intramuscular injection and the survival curves or bacterial loads in blood, spleen and kidney 4-days post-infection were determined. In this model, bacteraemia and bacterial loads in spleen and kidney were consistently significantly lower in the group infected with *ΔbfpR* relative to the wild-type (Figure 7C). The fish groups infected with *ΔbfpR* displayed a higher survival rate compared to those infected with the wild-type strain (Figure 7D).

Altogether, the results showed that both Hfp and BfpR systems are necessary for full virulence of *F. psychrophilum* in rainbow trout with no apparent functional redundancy.

## Discussion

Here, we report the identification of two heme/hemoglobin outer membrane receptors conserved in the species and demonstrate that they act differently to support bacterial growth, under high hemoglobin condition for BfpR and under iron scarcity for HfpR. We also show by experimental infection in a natural host, the rainbow trout, that both receptors are required for full virulence (Figure 8). The findings have relevance to other important fish pathogens such as *Flavobacterium columnare* and *Tenacibaculum maritimum* that possess homologs of the two uptake systems reported here.

**Figure 8.**
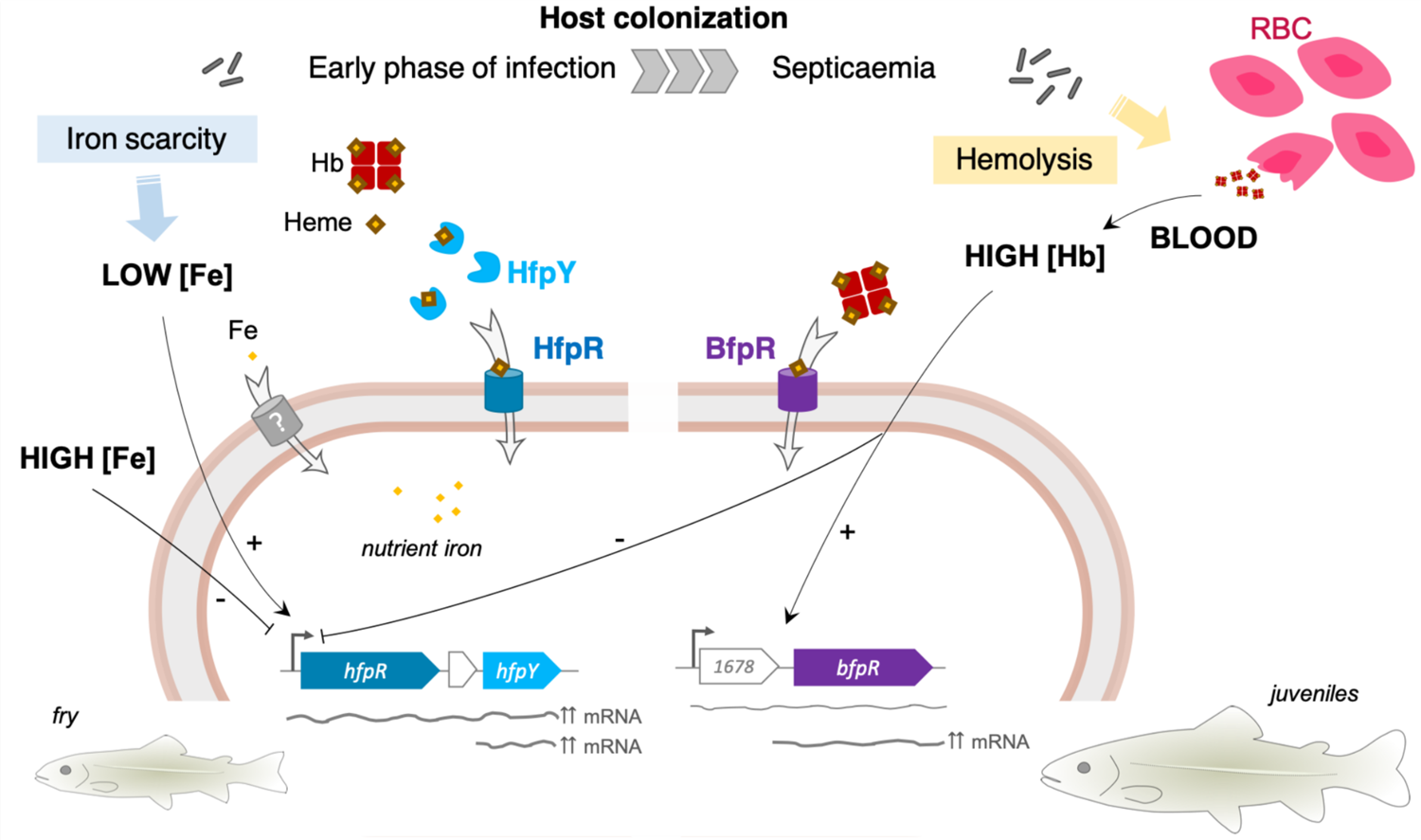
Graphical model of non-redundant heme/hemoglobin uptake systems of *F. psychrophilum.* This study showed that HfpR and BfpR outer membrane receptors both mediate heme uptake, but lack functional redundancy. We propose that HfpR and BfpR act in distinct environmental conditions during pathogenesis. During the early phase of infection, *F. psychrophilum* encounters low iron conditions due to the sequestration of essential nutrients by the host; iron is either bound to proteins or complexed in heme-binding proteins [26]. The Hfp system is upregulated in low iron and mediates iron acquisition from heme or hemoglobin (Hb); our results indicate that the HfpY hemophore identified here outcompetes host sequestration systems. In contrast, BfpR is up-regulated in the presence of high amounts of hemoglobin from blood, an iron source available upon massive lysis of erythrocytes (RBC) during late stages of septicaemia. In these conditions, the Hfp system is strongly repressed and BfpR likely mediate heme uptake by direct binding to hemoglobin. The importance of BfpR for full virulence was observed in juvenile rainbow trout but not in fry. Altogether, these results suggest that *F. psychrophilum* evolved two heme acquisition systems to adapt to the range of substrate concentrations encountered during the infection process as well as to the physiological changes occurring during the life cycle of fish.

### Expression of the hfp and bfpR loci is tightly regulated and indicates a role under distinct iron conditions

Complex regulatory mechanisms act in bacterial pathogens to maintain iron at intracellular levels needed for bacterial development and below its toxicity threshold [26, 49]. Here, we show that *hfpR* and *hfpY* are up-regulated in an iron-dependent manner. The promoter of the operon was induced under low-iron condition, whereas availability of ferric iron led to a dose-dependent transcriptional repression, which was also observed in presence of a high level of hemoglobin. In many bacteria, such regulation is mediated by Fur family iron-responsive regulators. Typically, under iron-replete condition the Fe^2+^-Fur holoprotein binds to the promoter of iron-regulated genes thus preventing transcription, while under low-iron apo-Fur is released and transcription occurs [50]*. F. psychrophilum* genome contains one member of the Fur family (THC0290_0812). Several attempts to construct a deletion mutant failed at the second recombination event, plasmid excision resulting only in wild-type genotype. This may be either due to the requirement of the Fur homolog for growth or to impaired expression of the downstream purine biosynthesis gene *purA* [31].

Several evidences indicate that post-transcriptional regulatory events occur in both *bfpR* and *hfp* mRNAs. In the wild-type strain, mRNA levels for *hfpY* were higher than for *hfpR* under all tested biological conditions (F7 in *Supporting Information*)[31]. In-depth analysis of the transcriptional structure of the *hfp* locus revealed the existence of two mRNAs (*hfpR*-*1813*-*hfpY* and *hfpY*) and only the polycistronic RNA was triphosphorylated at the 5’ terminus, indicating a single primary transcript. Moreover, no binding site of sigma factor was found in the sequence surrounding the 5’ end position of *hfpY* mRNA, which is consistent with a post-transcriptional regulation. We also showed that *hfpY* mRNA level was lower in the *ΔhfpY* strain carrying the chromosomal transcriptional fusion P*_hfpR_*-*hfpY* than in the wild-type, suggesting that the natural 32-nt 5’ untranslated region of *hfpY* mRNA is important for its stability. This lower expression was associated to only partial restoration of *in vitro* and *in vivo* phenotypic defects of *ΔhfpY*, indicating that accurate *hfpY* mRNA quantity is needed for full activity of the Hfp system. Moreover, a putative low permissive transcriptional state of the ectopic chromosomal site is unlikely as *hfpR* mRNA level in complemented *ΔhfpR* carrying the P*_hfpR_*-*hfpR* fusion was similar than in wild-type. It is noteworthy that several *hmuY* transcripts are also produced in *P. gingivalis,* the monocistronic form being more abundant than the bicistronic *hmuY-hmuR* one, suggesting a similar hemophore upregulation in the two species despite distinct operonic structure [45]. In bacteria, the stability of RNAs released by processing of a polycistronic RNA may differ, which affects the relative expression levels of genes within a same operon. Our hypothesis is that a stable *hfpY* transcript is released after maturation of *hfpR*-*1813*-*hfpY* mRNA, resulting in higher quantity of *hfpY* compared to *hfpR*, independently of iron level. A similar regulation was recently described in *Pseudomonas aeruginosa* for the *hasR*-*hasA* operon encoding the hemophore-dependent Has system [51]. Production of large amounts of hemophore likely provides a significant advantage to maintain effective concentration relative to its cognate receptor while diffusing into the surrounding environment.

In contrast to *hfp*, we showed that *bfpR* mRNA accumulated in the presence of hemoglobin under iron-replete condition. The upregulated RNA was monocistronic and likely originated from maturation of a longer RNA starting at −673 bp from the *bfpR* start codon and carrying a putative hypothetical gene (THC0290_1678). RNA level of the 5’ region of this long transcript was unchanged in the presence of hemoglobin. The underlying mechanism, presently not known, likely involves the condition-dependent modulation of *1678*-*bfpR* mRNA cleavage by a ribonuclease. Typically, such regulation can be mediated by structural change of the target RNA or by binding of a small regulatory RNA, this interaction acting positively or negatively on the stability and translation of the target [52]. For instance, a sRNA-dependent mRNA cleavage of the 5’ UTR of *colA* mRNA, which encodes the collagenase in *Clostridium perfringens*, releases a more stable transcript allowing overexpression of the toxin [53]. Several *F. psychrophilum* sRNAs were shown to accumulate in the presence of blood [31] and are regulator candidates for the *bfpR* post-transcriptional regulation. It is also noteworthy that the regulatory event leading to *bfpR* accumulation occurs also under specific conditions without hemoglobin such as in biofilm [54] or in stationary phase [31].

### Role of the two heme/hemoglobin receptors in heme acquisition and host colonization

For vertebrate pathogens, the most abundant form of iron is bound within heme, and mainly sequestered into hemoglobin in erythrocytes and into other host proteins. In rainbow trout, serum albumin, hemopexin-like proteins and hemoglobin are among the most abundant proteins in plasma [55].

*In vitro*, when ferric iron was deprived and heme or hemoglobin was the sole iron source, the growth of *F. psychrophilum* was strongly impaired in the absence of HfpR, but only slowed in the absence of HfpY. The importance of the hemophore was nevertheless obvious *in vivo*, the lack of HfpY resulting in reduced mortality and bacterial loads in spleen. These results indicate that the outer membrane receptor is able to independently capture and transport heme, although cooperation with its cognate hemophore is needed *in vivo*, likely to outcompete the host sequestration systems (Figure 8). The effectiveness of hemophore-based receptors to capture very low levels of heme was exemplified for several acquisition systems, such as the Has system found in *Serratia marcescens* and other Gram-negative bacteria: the affinity of HasR for the hemophore was 100-fold higher than for free heme [56, 57].

Direct binding of various substrates to the HfpR receptor was not assayed here, but meaningful information can be deduced from phenotypic results. The Hfp system was able to provide nutrient iron by the uptake of heme from hemoglobin. Previously, it was established that *F. psychrophilum* cells are highly proteolytic and that a T9SS-deficient mutant is unable to consume hemoglobin [21]. This indicates that the Hfp system may require additional factors to efficiently retrieve heme from hemoproteins. We propose that hemoglobin cleavage by exopeptidases contribute to release heme molecules which are then captured by the hemophore HfpY for delivery to the outer membrane receptor. Other hemoproteins such as serum albumin or hemopexin may similarly serve as alternative heme source after their proteolysis by secreted peptidases.

The expression profile of *bfpR* and the phenotypic characterization of the corresponding deletion mutant suggested a role in adaptation to hemoglobin overload, a stress condition that the pathogen likely undergoes during its lifecycle (Figure 8). Indeed, infected fish develop hemorrhagic septicaemia [13]. *F. psychrophilum* cells are hemolytic and high bacterial load in blood could result in the release of large amounts of hemoglobin. Bacterial cells also accumulate at high level in spleen and kidney, two hematopoietic tissues in teleost fish. In particular, the spleen acts as a filter for blood-derived substances and plays a major role for pathogens clearing and turnover of erythrocytes through phagocytosis by melanomacrophages [58]. This infection niche thus accumulates large amount of heme. We assume that the hemoglobin concentration (1%) used for *in vitro* experiments properly mimics massive hemolysis, as total hemoglobin levels of 6 to 12 % (g/dl) in blood of healthy rainbow trout were reported [59]. The results showed that the presence of 1% hemoglobin did not affect the viability of *F. psychrophilum*, rather, the growth of wild-type was only slightly reduced and the final biomass was increased. In contrast, the lack of BfpR resulted in a greatly reduced growth rate. Under these conditions, the Hfp uptake system was strongly repressed, though free iron was available as nutrient iron source. We concluded that *F. psychrophilum* cells can sustain high concentration of hemoglobin (equivalent to 620 µM heme) without toxicity and that BfpR is involved in this adaptation. Several Gram-negative pathogenic bacteria are reported to accumulate heme in their membrane with no apparent toxicity, whereas others are highly susceptible [60]. Additional experiments are ongoing in order to better characterize the function of BfpR in the physiology of *F. psychrophilum* under hemoglobin overload.

Experimental infection in rainbow trout fry using intramuscular injection or immersion models highlighted the major contribution of the Hfp system to virulence and host colonization. In contrast, the lack of BfpR had no significant impact on these experiments. It is noteworthy that experimental infection of rainbow trout at the early stage (∼1 to 5 g) by strain OSU THCO2-90 induced acute mortality 2- to 3-days post-infection, this tiny window making monitoring of the infectious process difficult. We thus infected juvenile rainbow trout (∼65 g) which immune system is expected to be more mature, allowing a better resistance against infection. Mortalities occurred between 4- and 8-days post infection with the wild-type, whereas the lack of BfpR resulted both in reduced bacterial loads in blood, spleen and kidney and in increased fish survival. This confirms that BfpR contributes to *in vivo* fitness in *F. psychrophilum*. Overall, we propose that both BfpR and Hfp systems support heme uptake with non-redundant functions and act at distinct substrate concentration ranges to adapt the physiology of *F. psychrophilum* to iron sources available in the surrounding environment. Many pathogenic bacteria evolved to produce multiple heme receptors, each of them contributing to host colonization. For instance, *P. aeruginosa* produces three outer membrane heme receptors: the hemophore-dependent systems HxuA and Has act mostly as sensors of extracellular heme, while the hemophore-independent PhuR receptor ensures the uptake of high amounts of heme providing nutrient iron for growth [61, 62].

### Sequence properties of the BfpR and HfpR receptors and the HfpY hemophore

Distinct TBDRs typically display similar 3D-structural organization but low-sequence similarity (∼20%). Predicting the substrate of TBDRs is challenging as specificity resides more in a few residues located in external loops and internal surface of the channel than in the global sequence conservation of the beta-barrel domain. Despite the high diversity between sequences, conserved amino acid motifs have been identified in heme/hemoglobin receptors characterized in other Gram-negative bacteria [63–65]. Multiple sequence alignments showed that the FRAP/NxxL motifs and the histidine residue located within the external loop L7 in other hemoglobin/heme receptors were conserved in BfpR (F11A in *Supporting Information*), suggesting that it interacts directly with hemoglobin. In the case of HfpR, the loop L7 was extended (78 AA instead of 34 AA for *P. gingivalis* HmuR) and motif conservation was only partial. No obvious conservation of the FRAP/NxxL motifs was found for the two other TBDRs (THC0290_0681 and THC0290_0369) that are overexpressed by low-iron conditions and in the presence of blood, respectively (Figure 1, F11B in *Supporting Information*). These FRAP/NxxL motifs and histidine residue are thought to mediate hemoglobin binding and heme utilization, although mutagenesis studies of *P. gingivalis* HmuR and *Haemophilus ducreyi* HgbA showed a different behaviour and other residues can be involved in substrate interaction [63–65].

We noticed that, compared to other TonB-dependent hemoglobin receptors, BfpR possesses an additional domain called NTE for N-terminal extension (Carboxypeptidase D regulatory-like, IPR008969) which is frequently found in TBDRs (*i.e.* 16 out of 23 TBDRs in strain OSU THCO2-90). This domain of unknown function may contribute to substrate uptake as recently exemplified for the levan acquisition system in *Bacteroides thetaiotaomicron* [66].

The low sequence similarity of BfpR and HfpR with other characterized heme/hemoglobin receptors makes the transfer of knowledge hazardous and future investigations are needed to decipher the heme transport mediated by BfpR and HfpR at the molecular level. Moreover, the very low sequence conservation of HfpY with other members of the HmuY family makes it probable that the hemophore of *F. psychrophilum* functions using an original heme-binding mechanism. Indeed, despite a general conservation of heme binding ability in all five HmuY-like proteins described so far in strict anaerobes (*P. gingivalis, Tannerella forsythia*, *Prevotella intermedia* and *Bacteroides vulgatus*), in depth studies revealed important differences in their mode of interaction with heme; for instance, heme coordination involved His134/His166 in *P. gingivalis* HmuY but Met140/Met169 in *T. forthysia* Tfo [44, 67].

### Other iron acquisition systems are predicted in the genome of F. psychrophilum

The majority of TBDRs encoded in the *F. psychrophilum* genome have unknown functions. Based on the expression profiles in plasma or blood, several other candidates may also contribute to iron acquisition from heme or other iron sources. Occurrence analysis of TBDR encoding genes in *F. psychrophilum* genomes identified several receptors present only in genomes from specific lineages. Molecular epidemiology studies of *F. psychrophilum* outbreaks revealed an epidemic population structure with limited number of clonal complexes that are associated with host fish species [6, 41]. Phenotypic diversity between isolates was reported and a few related genetic determinants identified so far by comparative genomics [33,68–73]. Here, we found a gene cluster containing an uncharacterized TBDR-encoding gene (FP2456 in strain JIP 02/86) present only in genomes belonging to the two predominant clonal complexes infecting rainbow trout (*i.e.,* CC-ST10 and CC-ST-90). The FP2456 protein belongs to the TonB-dependent siderophore receptor family (TIGR01783) and genes conserved in synteny encode two putative siderophore biosynthesis enzymes and a MFS-type efflux transporter. Further studies aiming to characterize the role of this predicted siderophore system in iron acquisition and *in vivo* fitness are of great interest but are currently limited by the lack of efficient mutagenesis tools to manipulate these isolates.

## Conclusion

This study provides molecular knowledge on three novel molecular factors involved in the infection process. The results will help to determine future directions of study for many aspects of heme acquisition. A brief list of interesting topics would be the characterization of regulatory events that coordinate expression of HfpR and BfpR receptors in response to environmental changes and identification of additional proteins involved in the process such as exopeptidases releasing heme from hemoproteins, putative heme oxygenase degrading heme to free ferrous iron, or inner membrane transporter mediating transfer to the cytoplasm. *F. psychrophilum* likely evolved several acquisition strategies to take advantage of the diversity of iron sources encountered during its lifecycle. Mutagenesis of other putative iron acquisition systems may also unveil additional factors favouring host colonisation and disease development. *F. psychrophilum* infections are a major threat for salmonid farming worldwide and innovative control strategies are needed to limit their impact. This study shows that heme outer membrane receptors have a major contribution to bacterial fitness in the fish host and are promising targets for innovative antibacterial strategies [74].

## Acknowledgements

The authors are grateful to the staff of the fish facilities (IERP, INRAE, 2018. Infectiology of fishes and rodent facility, doi: 10.15454/1.5572427140471238E12 and PEIMA, INRAE, Fish Farming Systems Experimental Facility, doi: 10.15454/1.5572329612068406E12), to Edwige Quillet and Nicolas Dechamp (INRAE GABI, France) for supplying fish and advice, and to Benjamin Fradet and Claire Gautier-Brossois for technical assistance. We thank Elise Borezée-Durant, Alexandra Gruss, Rodrigo Coronel-Tellez, Philippe Bouloc and Mirjam Czjzek for helpful discussions. This work has benefited from the MicroScope annotation platform for computational resources.

## Author contributions

Author contributions following the CRediT taxonomy (https://casrai.org/credit/) are as follows: Conceptualization: TR, DL; Data curation: ED, CG; Formal analysis: YZ, DL, TR; Funding acquisition: TR; Investigation: YZ, JFB, BK, TR; Project administration: TR; Resources: DR, CG, PN; Software: CG, PN; Supervision: DL, DR, ED, TR; Visualization: YZ, DL, TR; Writing – original draft: TR; Writing – review & editing: YZ, DL, JFB, BK, CG, DR, PN, ED, TR.

## Supporting information

**Supporting Information.pdf** This file contains Supplementary Methods (M1 to M3), Supplementary Figures (F1 to F11) and Supplementary Tables (T1 to T3).

## REFERENCES

1. FAO, editor The state of world fisheries and aquaculture 2020. Sustainability in action. 2020; Rome, Italy: Food and Agriculture Organization (FAO); (The State of World Fisheries and Aquaculture (SOFIA).

2. Bernardet JF. Flavobacteriaceae. Bergey’s Manual of Systematics of Archaea and Bacteria 2015.

3. Nematollahi A, Decostere A, Pasmans F, et al. *Flavobacterium psychrophilum* infections in salmonid fish [Research Support, Non-U.S. Gov’t Review]. Journal of fish diseases. 2003 Oct;26(10):563–74.

4. Avendano-Herrera R, Tapia-Cammas D, Duchaud E, et al. Serological diversity in *Flavobacterium psychrophilum*: A critical update using isolates retrieved from Chilean salmon farms. Journal of fish diseases. 2020 Aug;43(8):877–888.

5. Fujiwara-Nagata E, Chantry-Darmon C, Bernardet JF, et al. Population structure of the fish pathogen *Flavobacterium psychrophilum* at whole-country and model river levels in Japan [Research Support, Non-U.S. Gov’t]. Veterinary research. 2013;44:34.

6. Nicolas P, Mondot S, Achaz G, et al. Population structure of the fish-pathogenic bacterium *Flavobacterium psychrophilum* [Research Support, Non-U.S. Gov’t]. Applied and environmental microbiology. 2008 Jun;74(12):3702–9.

7. Nilsen H, Sundell K, Duchaud E, et al. Multilocus sequence typing identifies epidemic clones of *Flavobacterium psychrophilum* in Nordic countries [Research Support, Non-U.S. Gov’t]. Applied and environmental microbiology. 2014 May;80(9):2728–36.

8. Knupp C, Wiens GD, Faisal M, et al. Large-scale Analysis of *Flavobacterium psychrophilum* MLST Genotypes Recovered From North American Salmonids Indicates Both Newly Identified and Recurrent Clonal Complexes are Associated with Disease. Applied and environmental microbiology. 2019 Jan 18.

9. Castillo D, Christiansen RH, Dalsgaard I, et al. Bacteriophage resistance mechanisms in the fish pathogen *Flavobacterium psychrophilum*: linking genomic mutations to changes in bacterial virulence factors [Research Support, Non-U.S. Gov’t]. Applied and environmental microbiology. 2015 Feb;81(3):1157–67.

10. Fraslin C, Quillet E, Rochat T, et al. Combining Multiple Approaches and Models to Dissect the Genetic Architecture of Resistance to Infections in Fish. Frontiers in genetics. 2020;11:677.

11. Gomez E, Mendez J, Cascales D, et al. *Flavobacterium psychrophilum* vaccine development: a difficult task [Research Support, Non-U.S. Gov’t Review]. Microbial biotechnology. 2014 Sep;7(5):414–23.

12. Wiens GD, Lapatra SE, Welch T, et al. On-farm performance of rainbow trout (*Oncorhynchus mykiss*) selectively bred for resistance to bacterial cold water disease: Effect of rearing environment on survival phenotype. Aquaculture. 2013;388–391:128-136.

13. Barnes ME, Brown ML. A review of *Flavobacterium psychrophilum* biology, clinical signs, and Bacterial Cold Water Disease prevention and treatment. The Open Fish Science Journal. 2011 2011;4:1–9.

14. Decostere A, D’Haese E, Lammens M, et al. *In vivo* study of phagocytosis, intracellular survival and multiplication of *Flavobacterium psychrophilum* in rainbow trout, *Oncorhynchus mykiss* (Walbaum), spleen phagocytes. Journal of fish diseases. 2001 2001;24:481–487.

15. Evensen O, Lorenzen E. An immunohistochemical study of *Flexibacter psychrophilus* infection in experimentally and naturally infected rainbow trout (*Oncorhynchus mykiss*) fry. Diseases of aquatic organisms. 1996;25(1):53–61.

16. Wiklund T, Dalsgaard I. Association of *Flavobacterium psychrophilum* with rainbow trout (*Oncorhynchus mykiss*) kidney phagocytes *in vitro*. Fish & shellfish immunology. 2003 Nov;15(5):387–95.

17. Bertolini JM, Wakabayashi H, Watral VG, et al. Electrophoretic detection of proteases from selected strains of *Flexibacter psychrophilus* and assesment of their variability. Journal of aquatic animal health. 1994;6:224–233.

18. Hogfors-Ronnholm E, Wiklund T. Hemolytic activity in *Flavobacterium psychrophilum* is a contact-dependent, two-step mechanism and differently expressed in smooth and rough phenotypes. Microbial pathogenesis. 2010 Dec;49(6):369–75.

19. Otis EJ. Lesions of cold-water disease in steelhead trout (Salmo gairdneri): the role of Cytophaga psychrophila extracellular products. [MSc thesis]: University of Rhode Island, Kingston; 1984.

20. Papadopoulou A, Dalsgaard I, Linden A, et al. *In vivo* adherence of *Flavobacterium psychrophilum* to mucosal external surfaces of rainbow trout (*Oncorhynchus mykiss*) fry. Journal of fish diseases. 2017 Oct;40(10):1309–1320.

21. Barbier P, Rochat T, Mohammed HH, et al. The type IX secretion system is required for virulence of the fish pathogen *Flavobacterium psychrophilum*. Applied and environmental microbiology. 2020 Jun 12;86(16).

22. Duchaud E, Boussaha M, Loux V, et al. Complete genome sequence of the fish pathogen *Flavobacterium psychrophilum* [Research Support, Non-U.S. Gov’t]. Nature biotechnology. 2007 Jul;25(7):763–9.

23. Pérez-Pascual D, Rochat T, Kerouault B, et al. More than gliding: Involvement of GldD and GldG in the virulence of *Flavobacterium psychrophilum*. Frontiers in microbiology. 2017;8:2168.

24. Langevin C, Blanco M, Martin SA, et al. Transcriptional responses of resistant and susceptible fish clones to the bacterial pathogen *Flavobacterium psychrophilum* [Research Support, Non-U.S. Gov’t]. PloS one. 2012;7(6):e39126.

25. Marancik D, Gao G, Paneru B, et al. Whole-body transcriptome of selectively bred, resistant-, control-, and susceptible-line rainbow trout following experimental challenge with *Flavobacterium psychrophilum*. Frontiers in genetics. 2014;5:453.

26. Skaar EP. The battle for iron between bacterial pathogens and their vertebrate hosts. PLoS pathogens. 2010 Aug 12;6(8):e1000949.

27. Schauer K, Rodionov DA, de Reuse H. New substrates for TonB-dependent transport: do we only see the ‘tip of the iceberg’? Trends Biochem Sci. 2008 Jul;33(7):330–8.

28. Andrews SC, Robinson AK, Rodriguez-Quinones F. Bacterial iron homeostasis. FEMS Microbiol Rev. 2003 Jun;27(2-3):215–37.

29. Moller JD, Ellis AE, Barnes AC, et al. Iron acquisition mechanisms of *Flavobacterium psychrophilum*. Journal of fish diseases. 2005 Jul;28(7):391–8.

30. Alvarez B, Alvarez J, Menendez A, et al. A mutant in one of two *exbD* loci of a TonB system in *Flavobacterium psychrophilum* shows attenuated virulence and confers protection against cold water disease. Microbiology (Reading, England). 2008 Apr;154(Pt 4):1144–51.

31. Guérin C, Lee B-H, Fradet B, et al. Transcriptome architecture and regulation at environmental transitions in flavobacteria: the case of an important fish pathogen. ISME Communications. 2021 2021/07/07;1(1):33.

32. Rochat T, Barbier P, Nicolas P, et al. Complete genome sequence of *Flavobacterium psychrophilum* strain OSU THCO2-90, used for functional genetic analysis. Genome announcements. 2017 Feb 23;5(8).

33. Rochat T, Pérez-Pascual D, Nilsen H, et al. Identification of a novel elastin-degrading enzyme from the fish pathogen *Flavobacterium psychrophilum*. Applied and environmental microbiology. 2019 Jan 11.

34. Vallenet D, Calteau A, Cruveiller S, et al. MicroScope in 2017: an expanding and evolving integrated resource for community expertise of microbial genomes. Nucleic acids research. 2017 Jan 04;45(D1):D517-D528.

35. Pei J, Kim BH, Grishin NV. PROMALS3D: a tool for multiple protein sequence and structure alignments. Nucleic acids research. 2008 Apr;36(7):2295–300.

36. Zhu Y, Thomas F, Larocque R, et al. Genetic analyses unravel the crucial role of a horizontally acquired alginate lyase for brown algal biomass degradation by *Zobellia galactanivorans*. Environ Microbiol. 2017 Jun;19(6):2164–2181.

37. Alvarez B, Secades P, McBride MJ, et al. Development of genetic techniques for the psychrotrophic fish pathogen *Flavobacterium psychrophilum*. Applied and environmental microbiology. 2004 Jan;70(1):581–7.

38. Lechardeur D, Cesselin B, Liebl U, et al. Discovery of intracellular heme-binding protein HrtR, which controls heme efflux by the conserved HrtB-HrtA transporter in *Lactococcus lactis*. The Journal of biological chemistry. 2012 Feb 10;287(7):4752–8.

39. Mandl CW, Heinz FX, Puchhammer-Stockl E, et al. Sequencing the termini of capped viral RNA by 5’-3’ ligation and PCR. Biotechniques. 1991 Apr;10(4):484, 486.

40. Quillet E, Dorson M, Le Guillou S, et al. Wide range of susceptibility to rhabdoviruses in homozygous clones of rainbow trout. Fish & shellfish immunology. 2007;22(5):510–519.

41. Duchaud E, Rochat T, Habib C, et al. Genomic diversity and evolution of the fish pathogen *Flavobacterium psychrophilum*. Frontiers in microbiology. 2018;9:138.

42. Krewulak KD, Vogel HJ. Structural biology of bacterial iron uptake. Biochim Biophys Acta. 2008 Sep;1778(9):1781–804.

43. Bielecki M, Antonyuk S, Strange RW, et al. *Prevotella intermedia* produces two proteins homologous to *Porphyromonas gingivalis* HmuY but with different heme coordination mode. Biochem J. 2020 Jan 31;477(2):381–405.

44. Bielecki M, Antonyuk S, Strange RW, et al. *Tannerella forsythia* Tfo belongs to *Porphyromonas gingivalis* HmuY-like family of proteins but differs in heme-binding properties. Biosci Rep. 2018 Oct 31;38(5).

45. Olczak T, Sroka A, Potempa J, et al. *Porphyromonas gingivalis* HmuY and HmuR: further characterization of a novel mechanism of heme utilization. Arch Microbiol. 2008 Mar;189(3):197–210.

46. Sieminska K, Cierpisz P, Smiga M, et al. *Porphyromonas gingivalis* HmuY and *Bacteroides vulgatus* BvuA Novel Competitive Heme Acquisition Strategy. Int J Mol Sci. 2021 Feb 24;22(5).

47. Baez WD, Roy B, McNutt ZA, et al. Global analysis of protein synthesis in *Flavobacterium johnsoniae* reveals the use of Kozak-like sequences in diverse bacteria. Nucleic acids research. 2019 Nov 18;47(20):10477–10488.

48. Meheust R, Huang S, Rivera-Lugo R, et al. Post-translational flavinylation is associated with diverse extracytosolic redox functionalities throughout bacterial life. Elife. 2021 May 25;10.

49. Chandrangsu P, Rensing C, Helmann JD. Metal homeostasis and resistance in bacteria. Nature reviews Microbiology. 2017 Jun;15(6):338–350.

50. Troxell B, Hassan HM. Transcriptional regulation by Ferric Uptake Regulator (Fur) in pathogenic bacteria. Front Cell Infect Microbiol. 2013;3:59.

51. Dent AT, Mourino S, Huang W, et al. Post-transcriptional regulation of the *Pseudomonas aeruginosa* heme assimilation system (Has) fine-tunes extracellular heme sensing. The Journal of biological chemistry. 2019 Feb 22;294(8):2771–2785.

52. Rochat T, Bouloc P, Repoila F. Gene expression control by selective RNA processing and stabilization in bacteria. FEMS microbiology letters. 2013 Jul;344(2):104–13.

53. Obana N, Shirahama Y, Abe K, et al. Stabilization of *Clostridium perfringens* collagenase mRNA by VR-RNA-dependent cleavage in 5’ leader sequence. Mol Microbiol. 2010 Sep;77(6):1416–28.

54. Levipan HA, Avendano-Herrera R. Different phenotypes of mature biofilm in *Flavobacterium psychrophilum* share a potential for virulence that differs from planktonic state. Front Cell Infect Microbiol. 2017;7:76.

55. Morro B, Doherty MK, Balseiro P, et al. Plasma proteome profiling of freshwater and seawater life stages of rainbow trout (*Oncorhynchus mykiss*). PloS one. 2020;15(1):e0227003.

56. Ascenzi P, di Masi A, Leboffe L, et al. Structural Biology of Bacterial Haemophores. Adv Microb Physiol. 2015;67:127–76.

57. Izadi-Pruneyre N, Huche F, Lukat-Rodgers GS, et al. The heme transfer from the soluble HasA hemophore to its membrane-bound receptor HasR is driven by protein-protein interaction from a high to a lower affinity binding site. The Journal of biological chemistry. 2006 Sep 1;281(35):25541–50.

58. Press CM, Evensen Ø. The morphology of the immune system in teleost fishes. Fish & shellfish immunology. 1999 1999/05/01/;9(4):309–318.

59. Martinez FJ, Garcia-Riera MP, Ganteras M, et al. Blood parameters in rainbow trout (*Oncorhynchus mykiss*): Simultaneous influence of various factors. Comparative Biochemistry and Physiology Part A: Physiology. 1994 1994/01/01/;107(1):95-100.

60. Anzaldi LL, Skaar EP. Overcoming the heme paradox: heme toxicity and tolerance in bacterial pathogens. Infection and immunity. 2010 Dec;78(12):4977–89.

61. Otero-Asman JR, Garcia-Garcia AI, Civantos C, et al. *Pseudomonas aeruginosa* possesses three distinct systems for sensing and using the host molecule haem. Environ Microbiol. 2019 Dec;21(12):4629–4647.

62. Smith AD, Wilks A. Differential contributions of the outer membrane receptors PhuR and HasR to heme acquisition in *Pseudomonas aeruginosa*. The Journal of biological chemistry. 2015 Mar 20;290(12):7756–66.

63. Bracken CS, Baer MT, Abdur-Rashid A, et al. Use of heme-protein complexes by the *Yersinia enterocolitica* HemR receptor: histidine residues are essential for receptor function. Journal of bacteriology. 1999 Oct;181(19):6063–72.

64. Fusco WG, Choudhary NR, Council SE, et al. Mutational analysis of hemoglobin binding and heme utilization by a bacterial hemoglobin receptor. Journal of bacteriology. 2013 Jul;195(13):3115–23.

65. Liu X, Olczak T, Guo HC, et al. Identification of amino acid residues involved in heme binding and hemoprotein utilization in the *Porphyromonas gingivalis* heme receptor HmuR. Infection and immunity. 2006 Feb;74(2):1222–32.

66. Gray DA, White JBR, Oluwole AO, et al. Insights into SusCD-mediated glycan import by a prominent gut symbiont. Nature communications. 2021 Jan 4;12(1):44.

67. Wojtowicz H, Guevara T, Tallant C, et al. Unique structure and stability of HmuY, a novel heme-binding protein of *Porphyromonas gingivalis*. PLoS pathogens. 2009 May;5(5):e1000419.

68. Castillo D, Jorgensen J, Sundell K, et al. Genome-informed approach to identify genetic determinants of *Flavobacterium psychrophilum* phage susceptibility. Environ Microbiol. 2021 May 14.

69. Cisar JO, Bush CA, Wiens GD. Comparative Structural and Antigenic Characterization of Genetically Distinct *Flavobacterium psychrophilum* O-Polysaccharides. Frontiers in microbiology. 2019;10:1041.

70. Madsen L, Dalsgaard I. Comparative studies of Danish Flavobacterium psychrophilum isolates: ribotypes, plasmid profiles, serotypes and virulence. 2000.

71. Ngo TP, Bartie KL, Thompson KD, et al. Genetic and serological diversity of *Flavobacterium psychrophilum* isolates from salmonids in United Kingdom. Veterinary microbiology. 2017 Mar;201:216–224.

72. Rochat T, Fujiwara-Nagata E, Calvez S, et al. Genomic Characterization of *Flavobacterium psychrophilum* Serotypes and Development of a Multiplex PCR-Based Serotyping Scheme. Frontiers in microbiology. 2017;8:1752.

73. Sundell K, Landor L, Nicolas P, et al. Phenotypic and Genetic Predictors of Pathogenicity and Virulence in *Flavobacterium psychrophilum*. Frontiers in microbiology. 2019;10:1711.

74. Shisaka Y, Iwai Y, Yamada S, et al. Hijacking the heme acquisition system of *Pseudomonas aeruginosa* for the delivery of phthalocyanine as an antimicrobial. ACS Chem Biol. 2019 Jul 19;14(7):1637–1642.

