## Supporting Information for "Two functionally distinct heme/iron transport systems are virulence determinants of the fish pathogen *Flavobacterium psychrophilum*"

### *Supplementary Methods*

**M1. Construction of the Δ*bfpR*, Δ*hfpR*, Δ*hfpY* and Δ*hfpR*Δ*bfpR* strains**

Three suicide vectors pGi27, pGi29 and pGi38 allowing *hfpR*, *hfpY* and *bfpR* deletion respectively, were constructed using pYT313 plasmid [1] by three-fragments Gibson assembly using oligonucleotides listed in **Table T2**. For pGi27, three DNA fragments were amplified by PCR as follows: the plasmid fragment using pYT313 DNA as matrix and primers TRO186/TRO187, the 2,044-bp fragment upstream and 2,129-bp fragment downstream of *hfpR* with OSU THCO2-90 chromosomal DNA and primers TRO336/TRO337 and TRO338/TRO339, respectively. pGi29 and pGi38 were constructed similarly except that the 2-kb upstream and downstream fragments were amplified using primers TRO358/TRO359 and TRO360/TRO361 for *hfpY,* and TRO408/TRO409 and TRO410/TRO411 for *bfpR*.

Each resulting plasmid was introduced by conjugation in strain OSU THCO2-90 and colonies carrying chromosomal integration of the plasmid were selected using erythromycin resistance. The second recombination event (plasmid excision) was obtained by plating a stationary phase culture inoculated with a single colony in TYES broth without antibiotics on TYES supplemented with 5% sucrose. The chromosomal structure of erythromycin-sensitive clones was determined by PCR in order to identify the deletion mutant. The chromosomal structure of clones selected after double crossing over was analyzed by PCR using primers TRO341/TRO342 and TRO341/TRO325 for Δ*hfpR*, TRO379/TRO380 and TRO381/TRO382 for Δ*hfpY* and TRO417/TRO418 and TRO414/TRO415 for Δ*bfpR*. The double mutant Δ*hfpR*Δ*bfpR* was constructed by introduction of pGi38 plasmid by conjugation in Δ*hfpR*.

**M2. Construction of pGi39 plasmid allowing chromosomal gene integration in *F. psychrophilum* and derivative reporter gene plasmids.**

Several suicide vectors allowing chromosomal recombination and gene insertion at the neutral intergenic region were constructed using pYT313 plasmid [1] by Gibson method using oligonucleotides listed in **Table T2** (**Figure F2**).

pGi37 was designed to carry the 2,021-bp fragment upstream and the 1,979-bp fragment downstream of the insertion site (between positions 1121879 and 1124880 of OSU THCO2-90 genome) and a K7 composed of the 160-bp promoter region of *rpsL* gene ($P_{rpsL}$), the *F. psychrophilum* codon adapted mCherry gene (mChFp; [2]) and an intrinsic terminator. Four DNA fragments were amplified by PCR as follows: the plasmid fragment using pYT313 DNA as matrix and primers TRO390/TRO419, the upstream and downstream fragments using OSU THCO2-90 chromosomal DNA and primers TRO420/TRO393 and TRO396/TRO397 respectively, and the $P_{rpsL}$-mChFp K7 with pGi19 DNA and primers TRO394/TRO395. pGi19 is a pCP23-derivative plasmid carrying the mChFp gene under the control of the $P_{rpsL}$ promoter. pGi19 was constructed by assembly of the pCP23 fragment amplified using the pCP23-$P_{rpsL}$ plasmid and primers TRO245/TRO246 and the mChFp gene amplified using pUC-mChFp as matrix [2] and primers TRO247/248. pCP23-$P_{rpsL}$ was constructed by assembly of a pCP23 fragment amplified using primers TRO125/TRO126 and the $P_{rpsL}$ fragment amplified using primers TRO108/TRO111.

pGi39 was designed to carry the upstream and downstream fragments of the insertion site flanking a K7 composed of $P_{rpsL}$, a multiple cloning site and an intrinsic terminator. pGi39 was constructed by Gibson assembly of the plasmid fragment amplified using pGi37 with primers TRO443/TRO444 and ssDNA primers TRO421/TRO422 (300 ng each) carrying the MCS sequence.

pGi42 was designed to allow chromosomal integration of the elastase gene FP0506 under the control of $P_{rpsL}$ at the insertion site. pGi42 was constructed by Gibson assembly of the plasmid fragment amplified using pGi37 DNA as matrix with primers TRO434/TRO435 and the FP0506 fragment amplified using JIP02/86 chromosomal DNA with primers TRO436/TRO437.

The chromosomal structure of clones selected after double crossing over was analyzed by PCR using primers TRO406/TRO407 and TRO402/TRO403.

**M3. Construction of pGi39 derivative plasmids allowing complementation of *hfp* and *bfpR* deletion mutants.**

Ectopic complementation of deletion mutants was performed by constructing a set of pGi39-derivative plasmids that allow gene insertion under the control of native promoter into the chromosome (**Figure F2**).

pGi44 that allows ectopic integration of the *hfpR* gene with the 240-bp upstream region was constructed as follows: the pGi39 fragment was amplified using pGi37 DNA as matrix without the $P_{rpsL}$-mCh sequence with primers TRO443/TRO445 and the $P_{hfpR}$-*hfpR* fragment using OSU THCO2-90 chromosomal DNA and primers TRO446/TRO447. Clones resulting from Gibson assembly were easily selected from false-positive clones (carrying pGi37 matrix) based on the absence of mCherry fluorescence signal.

pGi45 and pGi46 were designed to allow ectopic integration of the *bfpR* gene with the 306-bp upstream region ($P_{bfpR\_1}$-*bfpR*) and the two genes $P_{bfpR\_1}$-*bfpR*-$P_{hfpR}$-*hfpR* in tandem, respectively. They were constructed by assembly of the pGi39 fragment amplified using pGi37 DNA as matrix and primers TRO443/TRO445 and, for pGi45, the $P_{bfpR\_1}$-*bfpR* fragment amplified using primers TRO448/TRO449, or for pGi46, the $P_{bfpR\_1}$-*bfpR* and $P_{hfpR}$-*hfpR* fragments amplified using primers TRO448/TRO452 and TRO453/TRO447.

pGi55 was designed to allow ectopic integration of the *hfpY* gene under the control of $P_{hfpR}$ promoter and was constructed by Gibson assembly of a plasmid fragment carrying the $P_{hfpR}$ amplified using pGi44 DNA matrix and primers TRO519/TRO520 and the *hfpY* coding sequence fragment amplified using OSU THCO2-90 chromosomal DNA matrix and primers TRO521/TRO522.

### *Supplementary Figures*

**F1. Expression profiles of TonB-dependent receptors in *F. psychrophilum* OSU THCO2-90.**
Quantile-normalized log2-expression level of mRNAs encoding HfpR (top) and BfpR (bottom) across 32 biological conditions in duplicates as previously published in Guérin *et al* [2]. A full description of the biological conditions used in the condition-dependent transcriptome dataset is available at https://fpeb.migale.inrae.fr/. Briefly, iron deprivation (DIP) was generated by TYES broth supplementation with 25 µM 2,2'-dipyridyl. Plasma exposure was performed by incubating bacterial cells with 50% rainbow trout fish plasma during 5 min (Plasma). Blood conditions correspond to bacterial colonies grown either on TYES agar supplemented with 10% defibrinated horse blood (TYESA-Blood) or on 10% defibrinated horse blood agar only (Blood). Starvation conditions correspond to late stationary phase (T5) or colonies grown on TYES agar supplemented or not with fetal calf serum (TYESA-FCS; TYESA).

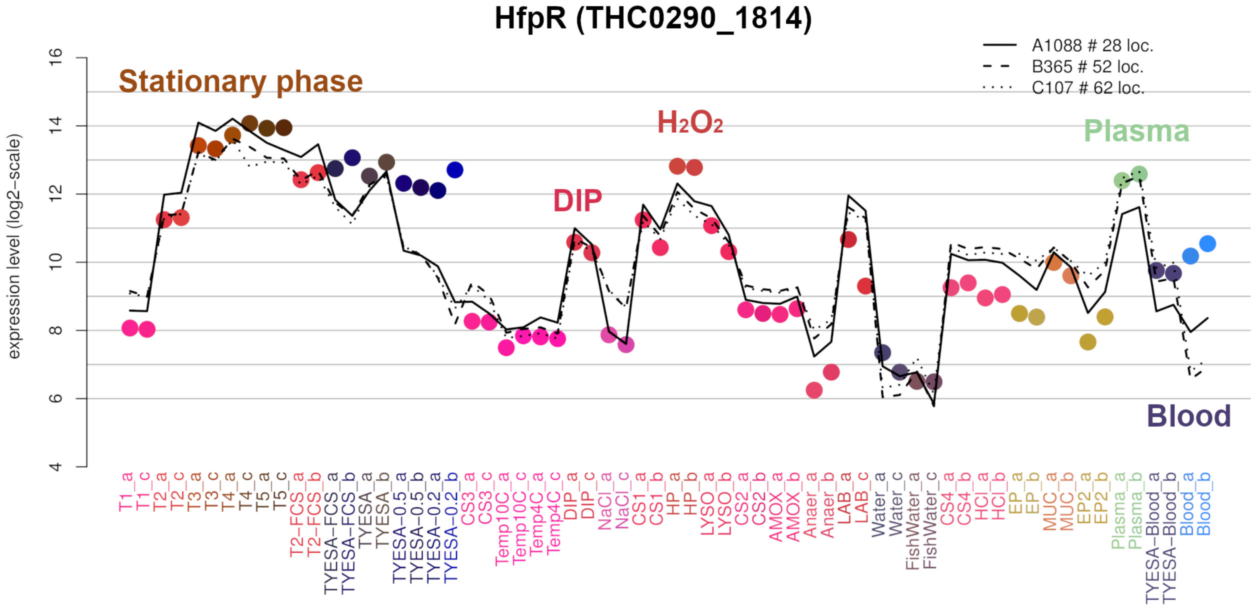

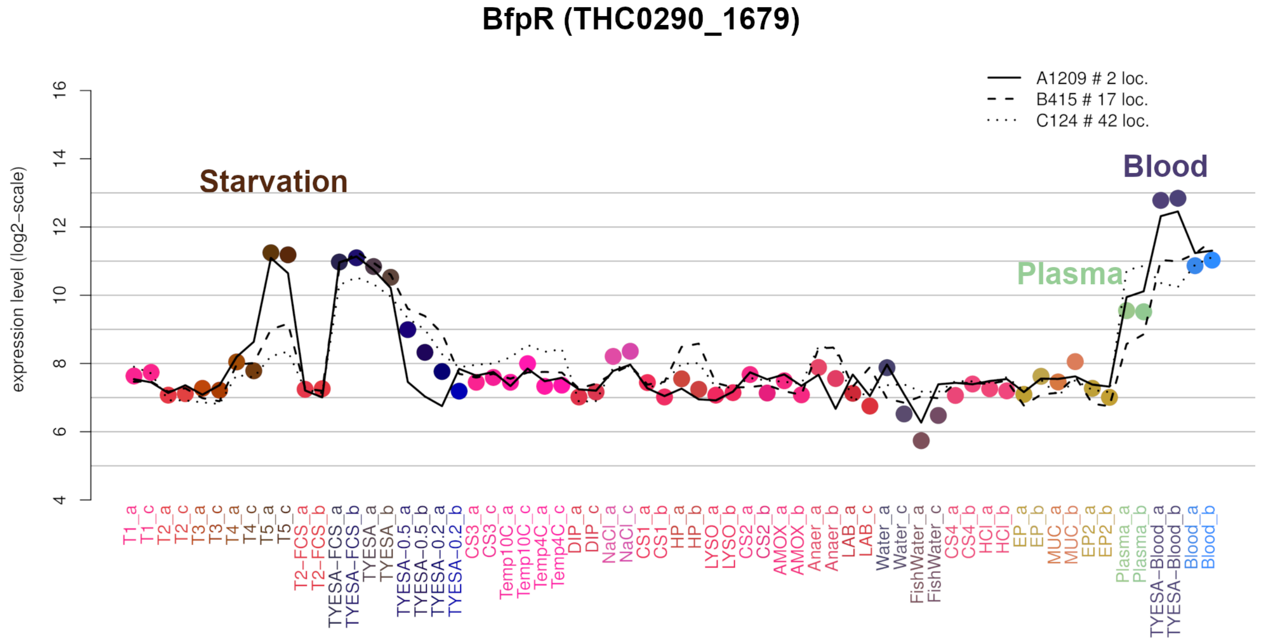

**F2. Schematic representation of the *F. psychrophilum* chromosomal gene expression platform and set of plasmids used for reporter genes expression and mutant complementation.**

The insertion site is within an intergenic region chosen for the absence of transcriptional signals and for its conservation within the *F. psychrophilum* species. pGi39 was constructed using pYT313 [1] as a backbone that carries, between homologous recombination regions, a chromosomal expression platform composed of a constitutive promoter ($P_{rpsL}$), a multiple cloning site and an intrinsic terminator. Gene inserted into the MCS of pGi39 is under the control of the constitutive promoter of *rpsL*. Restriction sites available in the multiple cloning site of pGi39 are: NotI, SpeI, SacI, SalI, XmaI (nucleotide sequence: gcggccgccgactagtgagctcgtcgacccggg). Excision of the DNA fragment carrying the $P_{rpsL}$, MCS and the intrinsic terminator can be achieved by double digestion of pGi39 with BamHI/PstI and allow cloning of a gene under its native expression signals using BamHI/PstI. Ectopic complementation of *hfpR*, *hfpY* and *bfpR* deletion mutants was performed by constructing a set of pGi39-derivative plasmids that allow gene insertion under the control of native promoter into the chromosome. Plasmids were constructed by Gibson assembly as described in **Methods M2**.

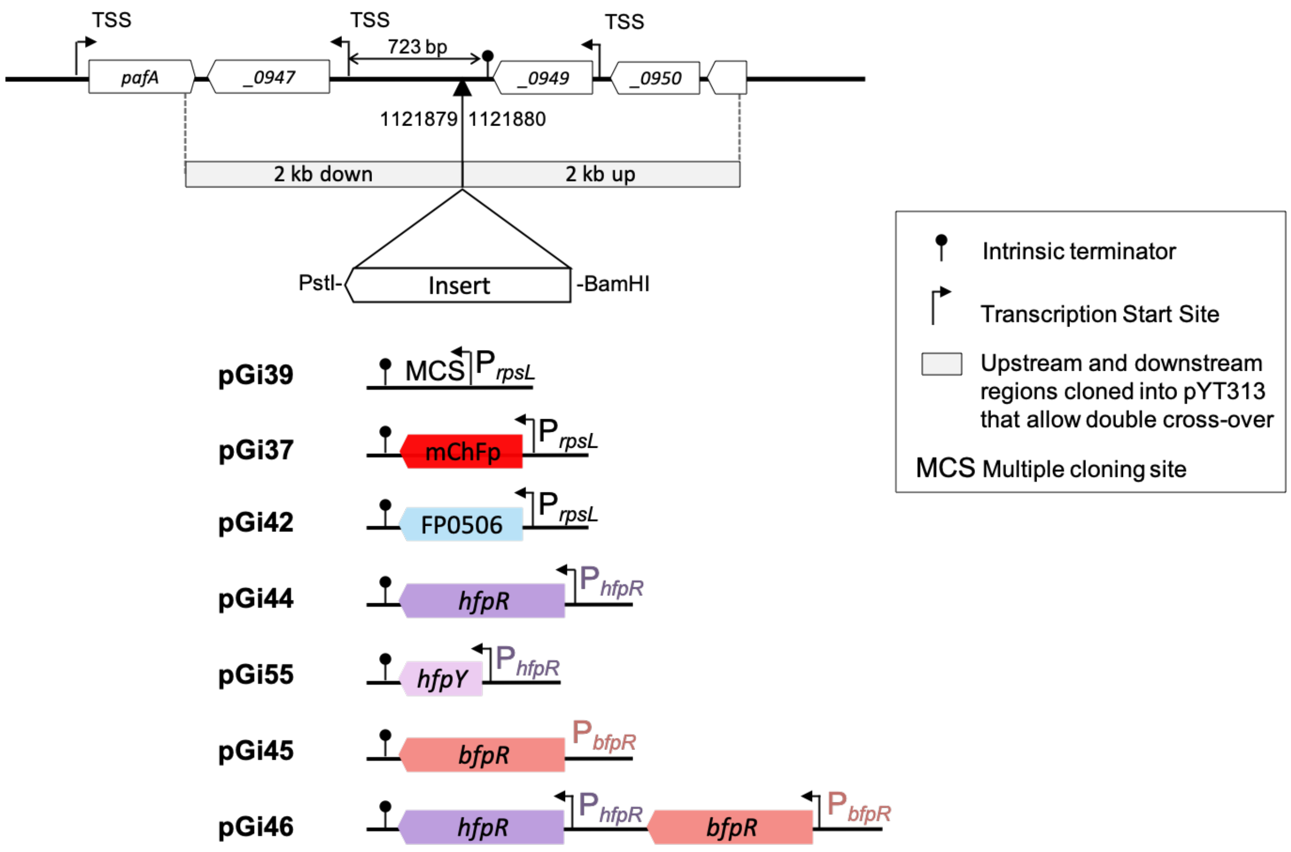

**F3. Validation of the chromosomal expression platform using reporter genes in strain OSU THCO2-90.**
**A.** mCherry expression of the strain carrying the $P_{rpsL}$-mChFp construct inserted into the chromosome after recombination with pGi37 plasmid. Expression of the *mCherry* gene was monitored in stationary phase cultures performed in TYES broth using whole-cell fluorescence with a Tecan Microplate Reader (Infinite 200 PRO). Excitation and emission wavelengths were set at 535 nm and 610 nm, respectively. mCherry expression (in arbitrary units) was estimated by dividing fluorescence intensity by $OD_{600}$. The mCherry expression was weak and near the background signal estimated using the wild-type strain, especially in the stationary phase. **B.** Elastinolytic activity of strain OSU THCO2-90 carrying the elastase-encoding gene FP0506 (from strain JIP 02/86)[3] at the chromosomal expression platform. Ten µL of stationary phase cultures performed were spotted on TYES agar supplemented with elastin (0.75%). Proteolytic activity was visualized as clearing zones around the bacterial growth after incubation at 18°C for 6 days. WT: OSU THCO2-90 wild-type; ecto::$P_{rpsL}$-mCh: insertion of pGi37 by homologous recombination at the insertion site (between positions 1121879 and 1121880); ecto::FP0506: insertion of pGi42 at the insertion site.

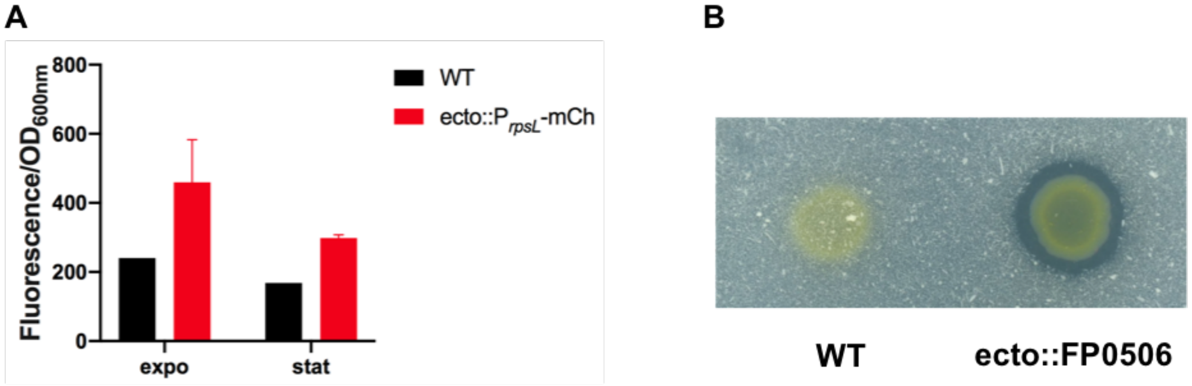

**F4. Transcriptional structure of *hfp* and *bfpR* loci analyzed by circular 3'-5′ RACE experiments.**
**(A)** Electrophoresis of amplicons from the first PCR (left) and the nested PCR (right) reactions of 3'-5′ RACE experiments. Total RNA treated by the pyrophosphohydrolase RppH (+RppH) or not (-RppH) was circularized, first-strand cDNA was synthetized using a primer specific of *bfpR* (TRO814), *hfpR* (TRO806) and *hfpY* (TRO811), then 3'-5' junctions were amplified using a set of primers, as follows: for *bfpR* cDNA, with internal *bfpR* primers (a/d, 1st PCR: TRO815/TRO816; nested PCR: TRO817/TRO818), *bfpR*-THC0290_1678 primers (b/d, 1st PCR: TRO815/TRO825; nested PCR: TRO817/TRO826) or *asd-bfpR* primers (c/d, 1st PCR: TRO815/TRO827; nested PCR: TRO817/TRO828); for *hfpR* cDNA with primers f/g (1st PCR: TRO807/TRO808; nested PCR: TRO809/TRO810) and for *hfpY* cDNA with primers e/g (1st PCR: TRO807/TRO812; nested PCR: TRO809/TRO813). **(B)** Determination of 5' and 3' ends of *bfpR* transcripts by TA-cloning and Sanger DNA sequencing of PCR products from +RppH treated samples. Boundaries of *bfpR* RNAs were identified by cloning 1st PCR products generated using a/d and nested PCR products generated using b/d (no amplification was obtained with c/d). 1, 2 and 3 indicate the most frequent positions identified for 5'ends and 3' end of *bfpR* mRNAs. **(C)** Determination of 5' and 3' ends of *hfpR and hfpY* transcripts by TA-cloning and DNA Sanger sequencing of PCR products from +RppH treated samples. Boundaries of *hfp* RNAs were identified by cloning 1st PCR products generated using f/g and e/g. 1, 2 and 3 indicate the most frequent positions identified for 5'ends and 3' end of *hfpR and hfpY* mRNAs. x-axis, genomic position in base pairs (bp) relative to the start position of downstream gene; y-axis, number of sequenced clones representative of each positions. Arrows indicate primers positions.

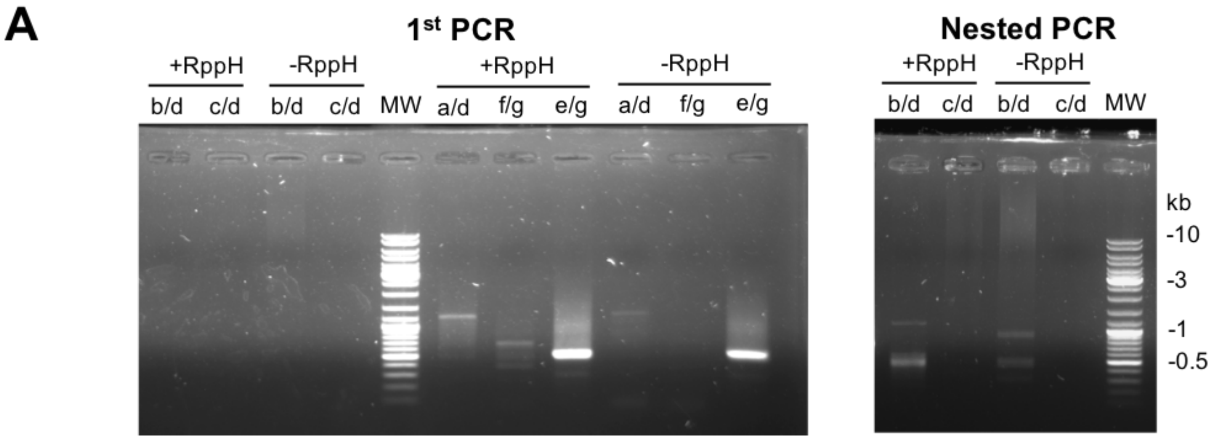

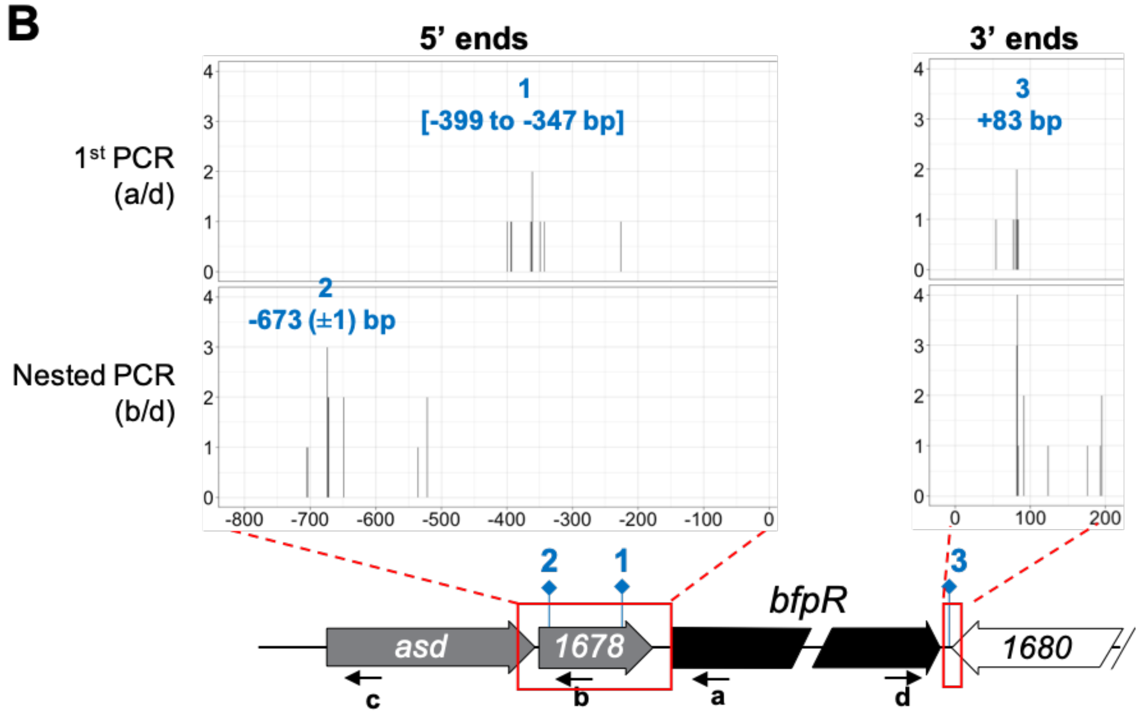

C

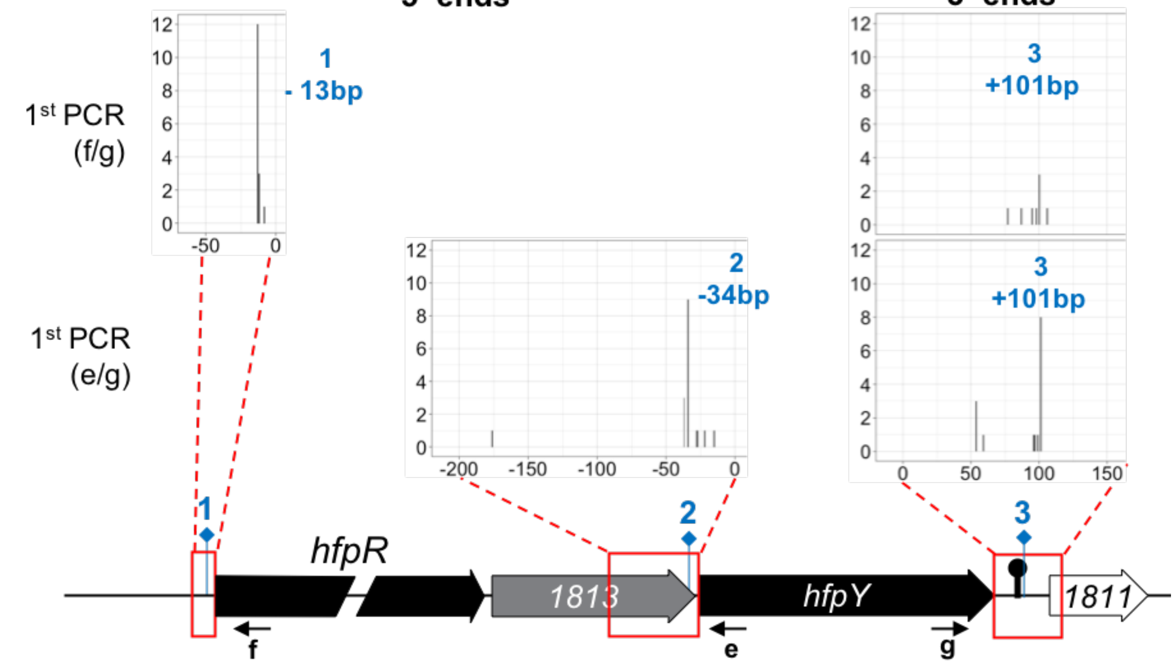

5' ends
3' ends
1st PCR (f/g)
1st PCR (e/g)
1
- 13bp
2
-34bp
3
+101bp
3
+101bp
hfpR
1813
hfpY
1811
1
2
3
f
e
g

**F5. Growth of *F. psychrophilum* OSU THCO2-90 at high concentration of hemoglobin.**
Growth of the wild-type strain at high hemoglobin concentration determined by viable cell measurements (CFU/mL) in TYES broth supplemented with 1% hemoglobin (purple) or not (black). Values represent the mean and standard deviation of three (A-C) or six (D) independent experiments.

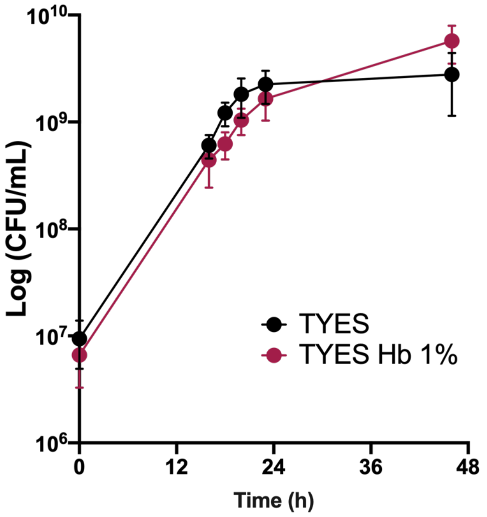

**F6. RT-qPCR measurement of gene expression in complemented deletion mutants compared to wild-type.**
mRNA level was quantified using RNA extracted from cultures of wild-type and deletion mutant carrying the corresponding ectopic gene copy (Figure F2). Cultures were performed in the biological condition corresponding to gene overexpression: TYES E20 Hb 1.25 µM for ecto::*hfpR* and ecto::*hfpY* strains, TYES Hb 1% for ecto::*bfpR* strain. Ct values of genes were normalized using the geomean of two reference genes (*rpsA* and *frr)*. RQ: Relative quantification of mRNA was expressed as $2^{-\Delta\Delta Ct}$ using wild-type as reference sample. Values are mean and standard deviation of three independent experiments. RT-qPCR showed that gene expression was restored in all complemented strains. Deletion of *hfpR* did not affect the mRNA level of *hfpY* and ectopic complementation resulted in similar *hfpR* mRNA level relative to wild-type. Deletion of *hfpY* did not affect the mRNA level of *hfpR*, however ectopic complementation resulted in lower *hfpY* mRNA level relative to wild-type (17%). Ectopic complementation of *bfpR* restored only partially the mRNA level of *bfpR* compared to wild-type (13%).

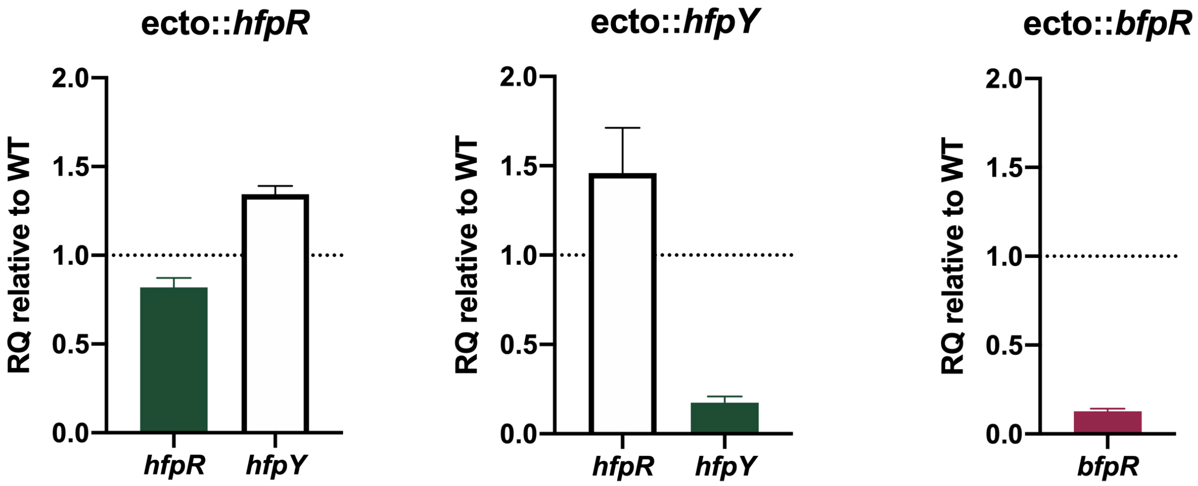

**F7. Distinct *hfpR* and *hfpY* mRNA levels across several biological conditions assessed by microarrays.** Quantile-normalized log2-expression level of microarray probes targeting *hfpR* or *hfpY* across several biological conditions. Hybridization values of 15 *hfpR*- and 11 *hfpY*-specific probes were retrieved from Guérin *et al* [2]. Total RNAs were extracted from cells from log to stationary phases (T1 to T5), under iron-depletion (DIP), peroxide stress ($H_2O_2$), oxygen limitation (Hypoxia), exposure to rainbow trout plasma (Plasma), and from colonies grown on TYES blood agar or blood agar.

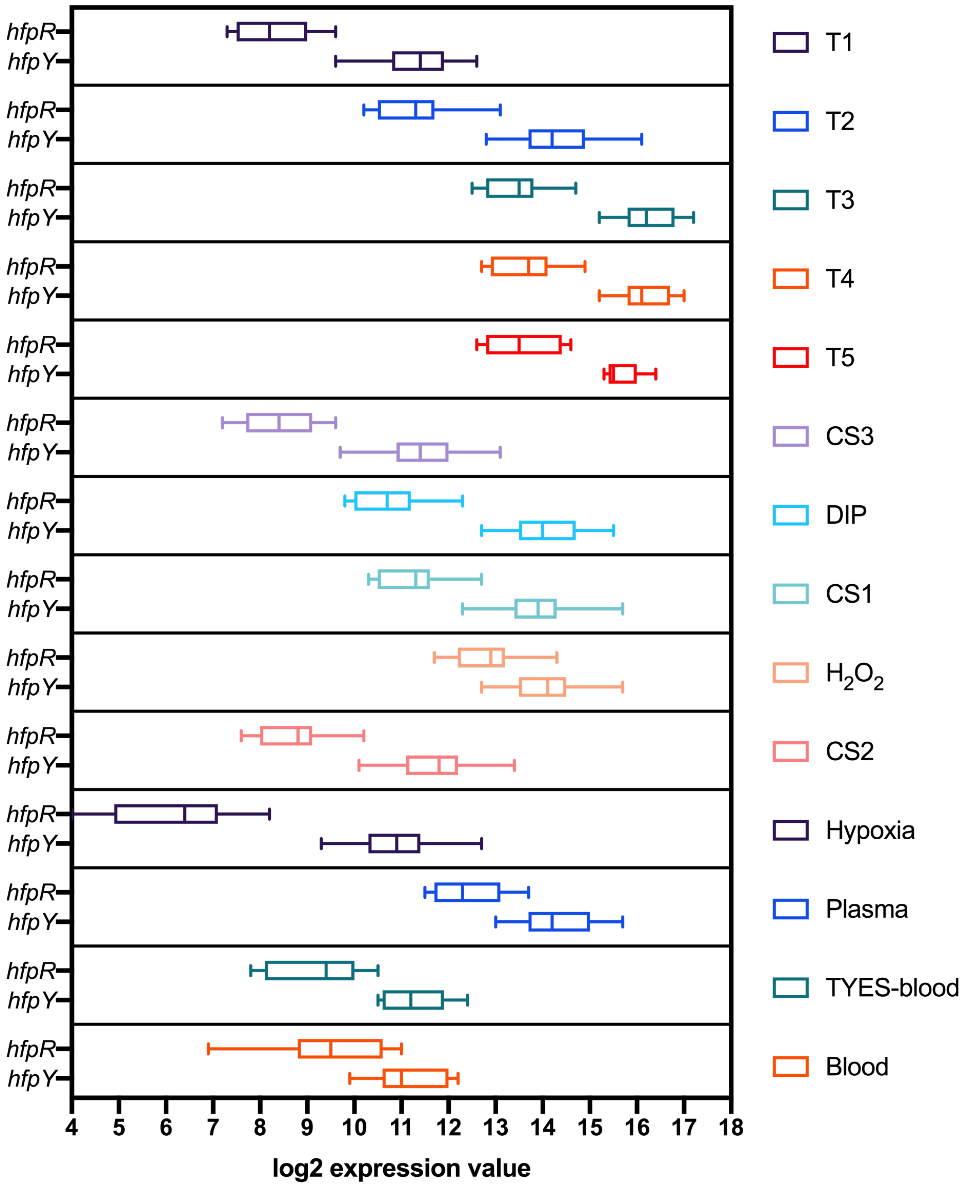

**F8. THC0290_1813 encoding a predicted inner membrane protein of unknown function is not required for growth on hemoglobin as an iron source.**

The THC0290_1813 mutant produced as part of a Tn*4351* mutant library in strain OSU THCO2-90 was used. Growth was compared to wild-type and Δ*hfpY* by optical density measurement at 600nm in 20 µM EDDHA iron-depleted TYES broth supplemented with 1.25 µM hemoglobin (▲) or not (♦). A reduced growth rate was observed only for Δ*hfpY* indicating that THC0290_1813 is not required under these conditions and that the Tn*4351* insertion does not block the expression of *hfpY* downstream gene. Wild-type: black, Tn4351::*1813*: pink, Δ*hfpY*: blue. Values represent the mean and standard deviation of three independent experiments.

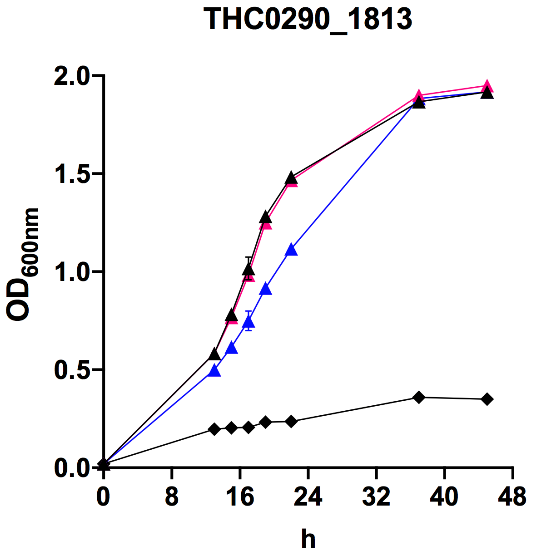

**F9. MSA of HfpY-like protein of *F. psychrophilum* with 5 characterized HmuY homologs.**

Multiple sequence alignment was constructed using PROMALS3D [4] with 5 sequences of HmuY-like proteins from *Porphyromonas gingivalis* (accession number: CAM31898), *Tannerella forsythia* (accession number: WP_046825712.1), *Prevotella intermedia* (accession numbers: AFJ07542, AFJ08449) and *Bacteroides vulgatus* (accession numbers: ABR39853) and the crystal structure of holo-HmuY from *Porphyromonas gingivalis* (PDB: 3H8T). Resulting alignment is provided below (*see* PROMALS3D web server for detailed output information). Sequences are colored according to predicted secondary structures (red: α-helix, blue: β-strand). `consensus_ss`: consensus secondary structure (α-helix: h; β-strand: e); `consensus_aa`: conserved amino acids are in bold and uppercase letters.

```
Conservation:             99                    9           6                       6 6
HmuY_Pg                1  MKKIIFSALCALPLI--VSLTSCGKK-----KDEPNQPSTP---------EAVTKTVTIDA--------S   46
3h8t_chainA_p001       1  --------------------------------------------------EAVTKTVTIDA--------S   12
Bvu_Bv                 1  MKRY-LSII-TILGMMLLPFSACDGILEG-IYDSPAASDSNELGFIRTDPSTHSGTIYIDA--------T   59
Tfo_Tf                 1  MKMRNVMTLALVALS--LAFVGC---------DKKDDVKET---------IKKS--KTIDA--------T   40
PinA_Pi                1  MKFKSFMAL-SCLTV--LLFSSCSNDDPT--PKPKPEQPKD---------VPTKTLSFITQEKAF----Q   52
PinO_Pi                1  MKTK-IFAV-ACLAT--LLFTSCSKDNN----DDPNPKPET---------TQVKHFETLM---------P   44
HfpY_Fp                1  MKNH-FFKVAVLALT--IFVSSCSKDEPATAVLTPQPQITT---------LEVKTVSNLNAPQVGMSFGP   58
Consensus_aa:             MK...h.hl.hhhhh..l.hotC.........cp.s...ps...........hpph.hlsA........s
Consensus_ss:               hhhhhhhhhhhhh  hhhhh                                 eeee

Conservation:              6  6    6       6   6                                      96 9 6  6
HmuY_Pg               47  KYETWQYFSFSKGEV--VNVTD--------------------------------YKNDLNWDMALHRYDV   82
3h8t_chainA_p001      13  KYETWQYFSFSKGEV--VNVTD--------------------------------YKNDLNWDMALHRYDV   48
Bvu_Bv                60  DYRRWTFIDFHTQKVDSVNVTD-----------------------------SEQ-KEPEEWDIAVHRYDV   99
Tfo_Tf                41  KYEMWTYVNLETGQT--ETHRDFSE--WHVMKN--------GKLLETIPAKGSEADIKIKWHIAIHRFDI   98
PinA_Pi               53  GYDKWVYVNLETGET--VMKDDVSEQEWRTYSDAGKKKDLFGKYDVTKTVEGKPSNAPDKWHLAFHVFDV  120
PinO_Pi               45  GYDSWIYIDLETGKF--EQQAELGKREFRKYKSM---MDPNYEVVGTEPAKGTDADLPKKWDIAFHITDA  109
HfpY_Fp               59  VSGEFTKFSFSENAV--VTN--------------------------------------DNWDVAFRGTTI   88
Consensus_aa:             .YcpWh@hshppsph...p.sD.................................c.s.pWclAhH.hDl
Consensus_ss:                 eeeee hh  e  ee                         ee               eeeeeeeee

Conservation:               9 66                        6           9       9            6
HmuY_Pg               83  RLNCGESGKGKGGAVFSG-----------KT---EMDQATTVPTDG-YTVDVL----GRITVKYEMGPDG  133
3h8t_chainA_p001      49  RLNCGESGKGKGGAVFSG-----------KT---EMDQATTVPTDG-YTVDVL----GRITVKYEMGPDG   99
Bvu_Bv               100  KTNAGAV-------LETG-----------FTGFSALRNADAMPEGA-YVEDVWTT--AKIAIDMSGMMDG  148
Tfo_Tf                99  RTNEGEA-------IATK-----------ET---EFSKVTGLPAGD-YKKDVEIKDKMLVGFNMADMMKS  146
PinA_Pi              121  RTNGAEA-------CMTD-----------TT---DIETIKTLPTNVKWVSDIK----AYLIYDMTGMMKN  165
PinO_Pi              110  RTNNGEV-------LMTG-----------ET---DLNKINALPAGN-YVADAP----ADIVVDMSRMQSE  153
HfpY_Fp               89  LVNGGAA-------IGIAGEPSRTGMGAVSIANNTLSGVTAFPAADTFRQDAN----SSYAIP-IGSGN-  145
Consensus_aa:             +hNsG.h.......l.os...........pT...phpphshhPhss.@..Dh.....t.lhhsh....ps
Consensus_ss:             ee   ee       ee                   hhhhhe      eeee      eeeee

Conservation:                        6                9                        96 96  69  9
HmuY_Pg              134  H-QMEYEEQGFSEVITGKKNAQGFASGGWLEFSHGPAGP-TYKLSK------RVFFVRGADGNIAKVQFT  195
3h8t_chainA_p001     100  H-QMEYEEQGFSEVITGKKNAQGFASGGWLEFSHGPAGP-TYKLSK------RVFFVRGADGNIAKVQFT  161
Bvu_Bv               149  N--IVYMESYYNEELS-----------KWLNVDKSNMPP-TYTLSN------KVYMVKLKDGTYAAVRLT  198
Tfo_Tf               147  K-FTVAGMAKVNPVLK-----------TWIVENPMGKAP-VLS--K------SVFVVKFKDGSYAKIKFT  195
PinA_Pi              166  PVVMGYMKSYVNMGLY-----------YWMHKVKGTMGEYALTMSKSDPKKAPVFLVKFKDGSYAVIQFT  224
PinO_Pi              154  G-VLGMVKTMLNGEMG-----------KWVKSN--GMGK-PKTVMG------NVFAVKFKNGNAALIKFK  202
HfpY_Fp              146  ---------------------------GWYNYN--GATN-IVTPLAG-----KVFVVKTHNGKYAKFEIL  180
Consensus_aa:             ...h.h.cp.hs..l.............Whp.s..shs..hho.........pVFhVKh+sGshA.lphh
Consensus_ss:                                         eeee        ee           eeeee     eeeeeee

Conservation:                             6   66 6 6
HmuY_Pg              196  DYQDA--------ELKKGVITFAYTYPVK-----  216
3h8t_chainA_p001     162  DYQDA--------ELKKGVITFTYTYPVK-----  182
Bvu_Bv               199  NYMNA--------SGVKGFMTIDYIYPFEL----  220
Tfo_Tf               196  DATND--------KQEKGHVSFNYEFQPK-----  216
PinA_Pi              225  GLKDA--------TGKKKEVSFKYKFVKKN----  246
PinO_Pi              203  DNLDK--------TGKKKAVSFDYKFIKKAK---  225
HfpY_Fp              181  SYYKDTPANPDAATSVGRHYTFKFVYQANSTTSF  214
Consensus_aa:             shbss........p..K.hhoFpY.@..c.....
Consensus_ss:             eee               eeeeeeeee
```

**F10. Kaplan-Meier survival curves of rainbow trout after infection with wild-type, mutant and complemented strains.**

Rainbow trout fry (Sy*Aut line, 5 g average weight) were infected by intramuscular injection. Each group (n=10) was challenged at a dose set to correspond to 1 LD50 (left column) or 10 x LD50 (right column) of the wild-type strain (black line). Exact viable cell counts were determined by plating bacterial suspensions in TYES agar and the results expressed as CFU/fish are indicated on the graphs. Comparison of wild-type and **(A)** Δ*hfpR* (red plain line) and ecto::*hfpR* (red dashed line); **(B)** Δ*hfpY* (blue plain line) and ecto::*hfpY* (blue dashed line); **(C)** Δ*bfpR* (green plain line) and ecto::*bfpR* (green dashed line); **(D)** Δ*bfpR*Δ*hfpR* (brown plain line) and ecto::*bfpR-hfpR* (brown dashed line). Survival curves were compared between groups of fish infected with mutant and wild-type using the Mantel-Cox log-rank test and the corresponding pvalue is indicated.

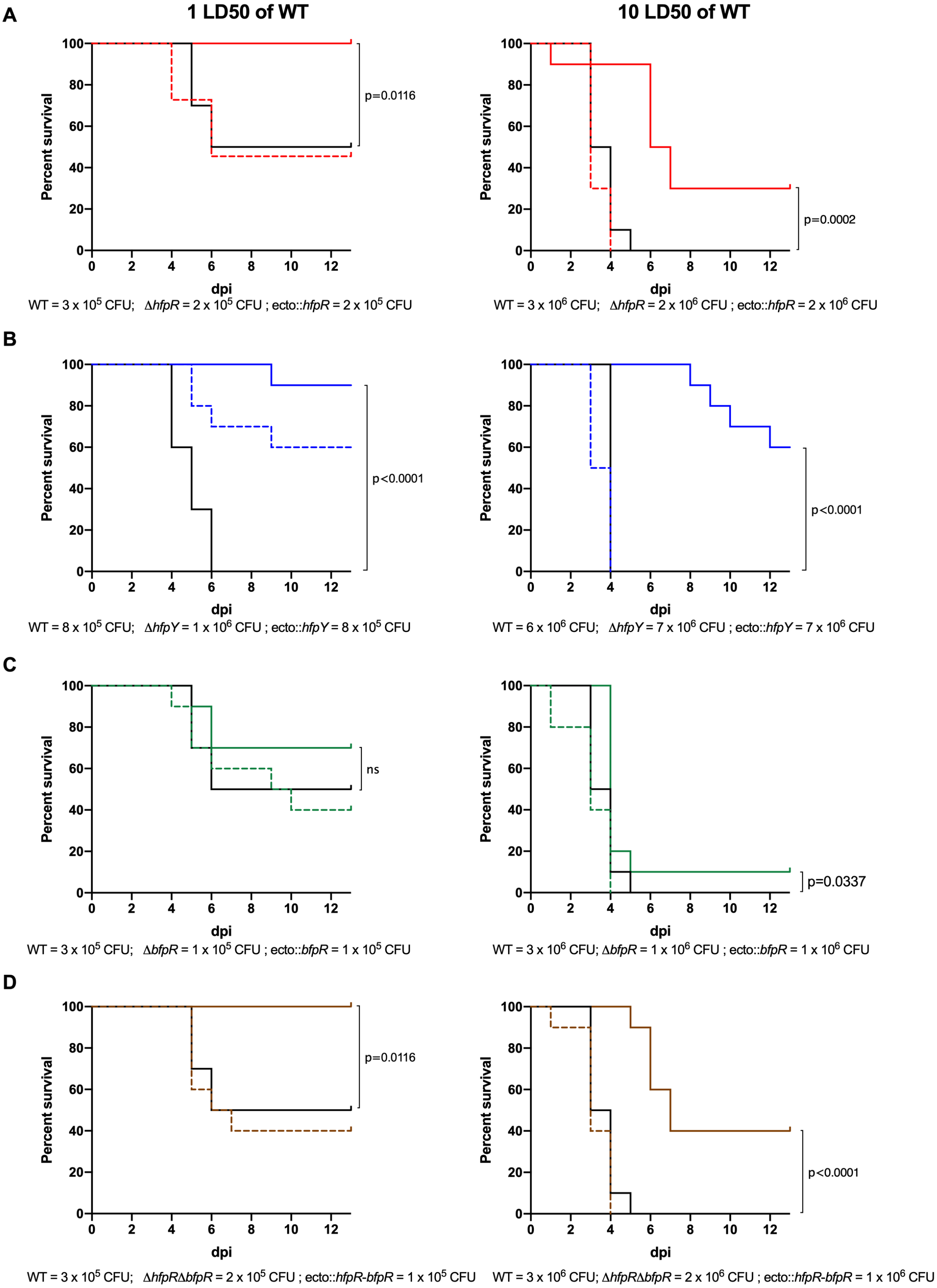

#### F11. MSA of HfpR and BfpR with other hemoglogin/heme TBDRs

Multiple sequence alignment was constructed using PROMALS3D [4] with 6 sequences of hemoglobin/heme TBDRs from Gram-negative bacteria and the crystal structure of ShuA from *Shigella dysenteriae* (PDB: 3fhh.1). Resulting alignment is provided below (*see* PROMALS3D web server for detailed output information). Sequences are colored according to predicted secondary structures (red: α-helix, blue: β-strand). `consensus_ss`: consensus secondary structure (α-helix: h; β-strand: e); `consensus_aa`: conserved amino acids are in bold and uppercase letters. DOMAINS: predicted protein family domains retrieved from the MicroScope annotation platform (Carboxypeptidase-like, regulatory domain: IPR008969; TonB-dependent receptor Plug domain: IPR012910; β-barrel domain: IPR000531). The predicted 22 transmembrane segments of the β-barrel are underlined and the surface-exposed loops are numbered L1 to L11. Alignment results for *F. psychrophilum* BfpR and HfpR **(A)** and for THC0290_0681 and THC0290_0369 protein sequence **(B)**.

**A.** BfpR and HfpR display conserved residues within the surface-exposed loop L7.

```
Conservation:           9               5          7
ShuA_Sd              1  MSRPQFTSLRLSLLALAVSATLPTF---------------------------------------------------------------------------   25
3fhh_chainA_p001        ----------------------------------------------------------------------------------------------------
PhuR_Pa              1  MPLSPP--FALRPC-LALLLLSPSLALAG-----------------------------------------------------------------------   26
HutA_Vc              1  MY-------KKSLLSSAIMLALVPSAYAD-----------------------------------------------------------------------   22
HpuB_Nm              1  MPIPF----KPVLAVAAIAQAFPAF-AADPAP--------------------------------------------------------------------   27
HgbA_Hd              1  MKAN-----KLSAITLCILGYAHTV-YAESNM--------------------------------------------------------------------   26
HmuR_Pg              1  MKSVVT---KQALIGLLFFSISIYSHAANPPAQPT-----------------------------------------------------------------   32
HfpR_Fp              1  MVI------NKAFGLHLLLLSFVGF--SQE-KQL------------------------------------------------------------------   25
BfpR_Fp              1  MK--------SFLVALLLGFAIFCN--AQN-KITGAITNSENKILETVLVSIPELNKETISDSNGNFLFLDLPKGKFKMSFSLVGYDEKLVSVFINKAEI   89
Consensus_aa:           M.......p..hhshhl..h..s...Ap........................................................................
Consensus_ss:                    hhhhhhhhhhhhhhhh
DOMAINS                 |   SIGNAL PEPTIDE   |     |   CARBOXYPEPTIDASE-LIKE, REGULATORY DOMAIN   (IPR008969)              .

Conservation:                       5   565 5                    5         5 5    595     5   5      79 5   55  579  5     6
ShuA_Sd             26  ---------AFATETMTVTATGNARSSFEAPMMVSVIDTSAPENQTATSATDLLRHV-PGITLDGTGR-TNGQDINMRGYDHRGVLVLVDGIRQGTDTG-  113
3fhh_chainA_p001     1  ------------TETMTVTATGNARSSFEAPMMVSVIDTSAPENQTATSATDLLRHV-PGITLDGTGR-TNGQDINMRGYDHRGVLVLVDGIRQGTDTG-   85
PhuR_Pa             27  --------NAVPLTPTTITATRTEQAVDSVPSTVSVQTREQLDRQNVNNIKELVRYE-PGVSVGGAGQRAGITGYNIRGIDGNRILTQIDGVELPNDFFS  117
HutA_Vc             23  --------DYASFDEVVVSTTRLNTQITDSAASVAVINASDIEQQMAEDIEGLFKYT-PGVTLTTNSR-QGVQGINIRGIEGNRIKVIVDGVAQPNQFDS  112
HpuB_Nm             28  -------QSAQTLNEITVTGTHKT-----QKLGEEKIRRKTLDKLLVNDEHDLVRYD-PGISVVEGGR-AGSNGFTIRGVDKDRVAINVDGLAQAESRSS  113
HgbA_Hd             27  --------QTEKLETIVVSSEDDS--VHNKNVGEIKKNAKALSKQQVQDSRDLVRYE-TGVTVVEKGR-FGSSGYAIRGVDENRVAVVVDGLHQAETISS  114
HmuR_Pg             33  ---DTIVSGNIALEDIVVTGSRTARLLKDVPVPTKVFKAKDIKAIAPSSFIDVLQYILPGIEFTKHGS---RDQLNAQGFDESSILFLVDGELISTGSTS  126
HfpR_Fp             26  ----SDSVKVATLNDVVITATRTERQLSSLPLPISIITSADISKAGISRLNEILTEQ-TGLITVPDFG--GAEGIQVQGLDAAYTLILIDGVPLVGRSAG  118
BfpR_Fp             90  NLNIRLESSSNHLEEVKIAATIFNKIQSQNVMKIEHKSIRELQQKGAATLMEGLASI-AGVSQVSTGT--SIGKPVIRGLSGNRVLVYTQGVRLENQQFG  186
Consensus_aa:           ........ps..hpphhloto...p..pp.sh....bp.p.lppb.hps..-llp...sGlphs..tp..t.p.hshpGh-.s.lhh.lDGl.bspp..t
Consensus_ss:                          eeeeee     hhh   eeeee hhhhhh     hhhhhhh    eeee         eeeeeee    eeeee  ee
DOMAINS                 ~~~~~~~~~~~~~|                     TONB-DEPENDENT RECEPTOR PLUG DOMAIN                              .

Conservation:                                      557 696 7 5 99 7679 5   5  5 7            5  5 5             5   5
ShuA_Sd            114  -------------HLNGTFLDPALIKRVEIVRGPSALLYGSGALGGVISYDTVDAKDLLQEGQSSGFRVFGTGGTGDHSLGLGASAFGRTENLDGIVAWS  200
3fhh_chainA_p001    86  -------------HLNGTFLDPALIKRVEIVRGPSALLYGSGALGGVISYDTVDAKDLLQEGQSSGFRVFGTGGTGDHSLGLGASAFGRTENLDGIVAWS  172
PhuR_Pa            118  GP---------YAQTHRNYVDPDIVKRVEILRGPASALYGSNAIGGAVSYFTLDPSDIIKDGKDVGARLKAGYESASHSWLTSATVAGRADDFDGLLHYG  208
HutA_Vc            113  GN--------SFLNSSRVDIDTDMVKSVEIVKGAASSLQGSDAIGGIVAFETKDPADILK-GRNMGGYAKLNYSSSDKTFSESIALANKSGDLESLVAYT  203
HpuB_Nm            114  EAFQELFGAYGNFNANRNTSEPENFSEVTITKGADSLKSGSGALGGAVNYQTKSASDYVSEDKPYHLGIKGGSVGKNSQKFSSITAAGRLFGLDALLVYT  213
HgbA_Hd            115  QGFKELFEGYGNFNNTRNGVEVENLKQAVIQKGADAIRTGSGSLGGTVSFESKDARDYLI-DKNYHFGYKTGYSSADNQKLHSVTAAGRYSDFDLLAVHT  213
HmuR_Pg            127  -------------GIDFERINPDDIERIEVLRGASSALYGSNAIGGVINIITRTAK------DPFRVSASARYDSRD-GQKYDVAAGVKRGIFTSQSGVQ  206
HfpR_Fp            119  -------------TLDLSRISVGNIDRIEIVKGASSSLYGSEALAGVINIITKKPK-----SDHFSGNLSYRYASFN-THDATANFVWKKKKFSGSFFTN  199
BfpR_Fp            187  -------------EEHGLGLNESGIESVEVIKGPASLLYGSDALGGVLYFNPEKFAK----PNSYLADFSQKYFSNTNGSNTSFGLKSSSENWKFLIRGA  269
Consensus_aa:           .............p.p...lps..lcplplh+Gsst.b.GSsAlGGhlsh.o.ps.D....spsh...hp....o.sp....shsh.s+...hp.bh.hs
Consensus_ss:                                hhhh eeeeee             eeeeeee          eeeeeeeeeeee    eeeeeeeee     eeeeeeee
DOMAINS                 ~~~~~~~~~~~~~~~~~~~~~~~~~~~~~~~~~~~~~~~~~~~~~~~~|      β-BARREL DOMAIN         L1                     .

Conservation:            5                                                           7         5
ShuA_Sd            201  SRDRGDLRQSNGE-----------------------------------TAPNDESINNMLAKGTWQIDSAQSLSGLVRYYNNDAREPKNPQTVEAS----  261
3fhh_chainA_p001   173  SRDRGDLRQSNGE-----------------------------------TAPNDESINNMLAKGTWQIDSAQSLSGLVRYYNNDAREPKNPQTVEAS----  233
PhuR_Pa            209  YRQGHETESNGGHGG--------------------------TGLSRSEANPEDADSYSLLGKLGWNYAEGSRFGLVFEKYKSDVDTDQKSAYGGPYDKGK  282
HutA_Vc            204  RRDGQEIQNFGSP------------------------------------DQQDNNANNLLVKLQYQLNPKHRLEFSGNYIRNKNDLENLEFS--------  259
HpuB_Nm            214  RRFGKETKNRSTEGNVEIKNDGYVYNPTDTGGPSKYLTYVATGVARSQPDPQEWVNKSTLFKLGYNFNDQNRIGWIFEDSRTDRFTNELSNLWTGTTTS-  312
HgbA_Hd            214  QRHGNELRNYGYRH----------------------YDGSVVRKEREKADPYKITKQSSLIKIGYQLNDTNRFTLGYDDSRNTSRGTDWSNAFTSYNGG-  290
HmuR_Pg            207  YRADKSYILADQFE--------------------------------QELNVAGNTTWNINQKFTISPTENLSFNLTGLVNLRKQHWTD------------  262
HfpR_Fp            200  YFDTNGYDLDKKNS---------------------------------LQTVEPYYNTTIQPKLYYDFSDKLKLIVGTRFYHQKQDNKTIINNK-------  259
BfpR_Fp            270  YNNHGDYKMPSDSL----------------------------------VTNTRFEEADVKTALGFSNS-KFSSVLRYNLNHLNLGIPEQIGLQTT-----  329
Consensus_aa:          ..ph.h..p..ppp....pb...........
Consensus_ss:           eee   ee                                              eeeeeeeeeeeee   eeeeeeeeeee
DOMAINS                 ~~~~~~~~~~~~~~~~~~~~~~~~~~~~~~~ L2 ~~~~~~~~~~~~~~~~~~~~~~~~~~~~~~~~~~~~~~~~~~~~~~~~~~~~~~~~ L3 ~~~~~~~~~~.

Conservation:
ShuA_Sd            262  -----------ESSNPMVDRSTIQRDAQLSYKLAPQGNDWLNADAKIYWSEVRINAQNTGS---------------------------------------  311
3fhh_chainA_p001   234  -----------ESSNPMVDRSTIQRDAQLSYKLAPQGNDWLNADAKIYWSEVRINAQ-------------------------------------------  279
PhuR_Pa            283  PAIPPSMLPGGMYQWRKGNDTLTRERYGLEHHFLLDSQVADRIQWSLNYQLAKTDQATREFYYPIT----------------------------------  348
HutA_Vc            260  -----------GYKNASGTDETTQYQLGIKHIWDAEFSLADRITWQFDVVGKEETGITDRTSKS------------------------------------  312
HpuB_Nm            313  ---------AATGDYRHRQDVSYRRRSGVEYKNELEHGPWDSLKLRYDKQRIDMNTWTWDIPKNY-----------------------------------  368
HgbA_Hd            291  ---------PFLKDVRHTNDQSNRKNISFVYENFDTNDFWDTLKITHNHQKIKLKARLDEYCDVNGEIDCPAIANPSGLYINDKGIFLDKHDGEITHKKE  381
HmuR_Pg            263  ----------------KIDFLYNSYDVKAGANWRISETSDLDISYHYDKYSRDTCLIKTEN---------------------------------------  307
HfpR_Fp            260  --------------NYQGDAVINEWNSQIKLEHNWNSK--INSEYEIYLTNYKTDAFLNNSK--------------------------------------  305
BfpR_Fp            330  -----------KRDPDYPRQGVVNHILSLHNNFYFRN---SKFDADFGYISNSRKEFADSTI--------------------------------------  377
Consensus_aa:           .............p....p....p.p..h.hp...p.p.h.phphph.....c.p...pp........................................
Consensus_ss:                               eeeeeeeeeeeeee    eeeeeeeeeeeeeeeeeee
DOMAINS                 ~~~~~~~~~~~~~~~~~~~~~~~~~~~~~~~~~~~~~~~~~~~~~~~~~~~~~~~~~~~~~~~~~~~~~~~~~~~~ L4 ~~~~~~~~~~~~~~~~~~~~~.
```

```
Conservation:
ShuA_Sd                     ----------------------------------------------------------------------------------------------------
3fhh_chainA_p001            ----------------------------------------------------------------------------------------------------
PhuR_Pa              349  ------------------------------------------------------------------------------------------------RKVL  352
HutA_Vc              313  ------------------------------------------------------------------------------------------------NGNI  316
HpuB_Nm              369  -------------------------------------------------------------------------------------------DLRGINSEV  377
HgbA_Hd              382  GEFNNYFDSKGKEVRVKGFNVDSILINCDQYDCSKPMQLLSSTNNGYGGSPNKYIYKTYELFEKTMNNGNGKYAVLEIRSSGHEKFSRVYLPSEKGYVEN  481
HmuR_Pg                     ----------------------------------------------------------------------------------------------------
HfpR_Fp              306  --------------------------------------------------------------------------------------------------NI  307
BfpR_Fp                     ----------------------------------------------------------------------------------------------------
Consensus_aa:             ....................................................................................................
Consensus_ss:
DOMAINS

Conservation:                                            5  9                                                             5
ShuA_Sd              312  --SGEYREQITKGARLENRSTLFADSFASHLLTYGGEYYRQEQHPGGAT--------------------------------------------TGFPQAK  365
3fhh_chainA_p001     280  ----EYREQITKGARLENRSTLFADSFASHLLTYGGEYYRQEQ--GSHT--------------------------------------------TGFPQAK  329
PhuR_Pa              353  RTRDTTYKERLWVFDSQLDKSFAI-GETEHLLSYGINLKHQKVTGMRSGTGTNLDT-------------------------GADSPRDALERSSDFPDPT  426
HutA_Vc              317  QKKDYLYSDKGFSFDSQLDKSFMV-SNTEHYIVYGFSLSDKDIENTNQEFNSI----------------------------------GKNNVIFYIPNAS  381
HpuB_Nm              378  YHSFRHIRQNTAQWTADFEKQLDF-SKAVWAAQYGLGGGRGDNANSDYSYFVKLY--------------------------DPKILTSNQAKITMLIENR  450
HgbA_Hd              482  QWKDRDLNTDTQQYNIDLTKSFKL-KSVEHNATYGGLYSEVKKSMTNRAGYEAYNRQWWANIFFGKENNKPNKCQPYNGNSFTTLCSHEDRLFSFLIPVK  580
HmuR_Pg              308  QKKLPIFDEDMHHLRAQYNLNLA----EVHLLNVGLEYIHDNVASPRLSSP------------------------------------------NDPGEKS  361
HfpR_Fp              308  LFEESFYDQWLIRPEIRTTFSI-----NNNKLTAGIGINHETLDRTYFGI-----------------------------------------------NAN  355
BfpR_Fp              378  --PVLDMNLNTLNYDLKYYLPNI----GKLEAIIGVQGMSQSNKNLGSEL--------------------------------------------LIPDAK  427
Consensus_aa:             ..p...hpp.h..hp.phpbsh.....s.h.h.hG....c.p...................................................s.h.psp
Consensus_ss:                 eeeeeeeeeeeeeeeeeee     eeeeeeeeeeeeeeeee                                                      e
DOMAINS                                                                                         L5

Conservation:              5      5  5            7 959                                                     5       5
ShuA_Sd              366  IDFSSGWLQDEITL-RDLPITLLGGTRYDSYRGSSDGY--------------------------------------------KDVDADKWSSRAGMTINP  420
3fhh_chainA_p001     330  IDFSSGWLQDEITL-RDLPITLLGGTRYDSYRGSSDGY--------------------------------------------KDVDADKWSSRAGMTINP  384
PhuR_Pa              427  VKTYALFAQDSISW---NDWTFTPGLRYDYTRMEPHITDEFLRTMK---------------------------QSQNTAVDESDKKWHRVSPKFGVTYDF  496
HutA_Vc              382  EKRYGFFIQDEIAF---DNLIVTPGIRFDSFETKPGDTSANPSL--------------------------------NDASEYKKYSDSALTARLGTVYKL  446
HpuB_Nm              451  SKYKFAYWNNVFHLGGNDRFRLNAGIRYDKNSSSAKDDPKYTTAI-----------------------------RGQIPHLGSERAHAGFSYGTGFDWRF  521
HgbA_Hd              581  TKTGALYVTDKIKL--NDKVNLDVAYRYDRIKHDPKYIPGTTPKLPTDLILGRFIEFKPKNTYATQDEKNENAEKNAVYLASKKTKFSANSYSATFSFDP  678
HmuR_Pg              362  VNNKILYGQYIYKV--TPKLVLSYGGRLDKHSGFG----------------------------------------------------LYYTSRLSAMYKC  407
HfpR_Fp              356  FNSQYGFLQYDYNP--SKKWNILAGFRYDNHNQYK----------------------------------------------------SQLSPKIAINYKV  401
BfpR_Fp              428  TNDIGALLTTNYEW---KSNVIQAGFRYDTRKINTESHGIIGDE---------------------------------GYFEAISKKYTSFNASVGYKTKL  491
Consensus_aa:             .p....@hp..hph....ph.l.sGhR@Dp.p..sc.h............................................pc.ph..ho.phth.hc.
Consensus_ss:             eeeeeeeeeeeeee     eeeeeeeeeeeeeeee                                                     eeeeeeeeeeee
DOMAINS                                                                                       L6

Conservation:                      55 77759   5 5  5 6                                       7  9559 5
ShuA_Sd              421  TNWLMLFGSYAQAFRAPTMGEMYNDSKHFSIGRFYTNYWV--------------------------PNPNLRPETNETQEYGFGLRFDDLMLSNDALEFK  494
3fhh_chainA_p001     385  TNWLMLFGSYAQAFRAPTMGEMYNDSKHFSIGRFYTNYWV--------------------------PNPNLRPETNETQEYGFGLRFDDLMLSNDALEFK  458
PhuR_Pa              497  AQHYTWYGQYAQGFRTPTAKALYGRFENLQA----GYHIE--------------------------PNPNLKPEKSQSFETGLRGKFD-------EGSFG  559
HutA_Vc              447  NQENRLFAQISQGFRAPDFQELYYSFGNPAH----GYVFK--------------------------PNPNLEAEDSVSYELGWRYNAD-------SVSNE  509
HpuB_Nm              522  TKHLHLLAKYSTGFRAPTSDETWLLFPHP------DFYLK--------------------------ANPNLKAEKAKNWELGLAGSGK-------AGNFK  582
HgbA_Hd              679  MDFLKIQAKYATGFRAPTSDEIYFVFQHP------SFSIY--------------------------PNLYLKAERSKNKEVAITLHKQ-------KSFLT  739
HmuR_Pg              408  S-HVTNRLSYAEGYRAPSLQEMYFFFNHG------AFFIY--------------------------GNPDLKPEKSRMLSYSAEAHWN-------KLTLM  467
HfpR_Fp              402  TDHLSLKTAVGYGFKAPDFRQLYFDFTNSSV----GYTVLGYNVADTRLNALDNQGQILKRIKDISFDNPLKPESSLNFNLGIFYQRS-------KLKIE  490
BfpR_Fp              492  TDNLSFRLNVASGFRSPNLAELTSNGVHEGS----NRYEI--------------------------GNSNLKNEQNIQTDLNLEYGNS-------HFEFF  554
Consensus_aa:             sp.hph..phtpGFRtPs..ph@....p.......s.hhb..........................sNssL+sEps.pbphth..p.p.........ph.
Consensus_ss:                eeeeeeeeeeee   hhhh               ee                                  eeeeeeeeeeeee          eeee
DOMAINS                                                  L7

Conservation:              5 9 5   6 7                                   5   5   9 7                            5
ShuA_Sd              495  ASYFDTKAKDYISTTVDFA---------------------AATTMSYNVPNAKIWGWDVMTKYTTDLFS-------LDVAYNRTRGKDTDT---------  557
3fhh_chainA_p001     459  ASYFDTKAKDYISTTVDFA---------------------AATTMSYNVPNAKIWGWDVMTKYTTDLFS-------LDVAYNRTRGKDTDT---------  521
PhuR_Pa              560  VAVFYNKYRDFIDEDALNTDST---------------GGNGQTFQSNNIERAVIKGVELKGRLELGAFGA-PQGLYTQGSVAYAYGRNKDN---------  634
HutA_Vc              510  LSIFYSDYDNFIDSQIVSGSF----------------KTRDAVHQSINIDKATIKGIELSNQFFWDRFMP-IVGFSSRIAAAYTEGKDGN----------  582
HpuB_Nm              583  LSGFKTKYRDFIELTYMGVSSDNPNKPTYAPLSDGTALVSSPVWQNQNRSSAWVKGLEFNGTWNLDSIGL-PQGTHAGVNVSYIKGKAKQTNG-------  674
HgbA_Hd              740  VNLFQTDYKDFLDLAYLKKGSLP--------YGNGGSQLETLLYQNVNRDKARVKGLEVNSKLHLGDVWRTLDGFNLSYKLSLQKGRMSSKVGEEGKQRD  831
HmuR_Pg              468  GNVFFNHVRNRIDFTYKGT----------------------DLIYTNVSKEMRIFGFEGQANVVLPYGFG------FRASYSYTYDRWKATDKAG-----  534
HfpR_Fp              491  VNAFYNAISNLIDTRAIAIKY-----------------TGQNVFSYYNVGKIFTYGTEFNSAYSFSNNLK------ISLGYQYLIAKDKSVIDDFETHQY  567
BfpR_Fp              555  ANGFYNNISNYIFIAPNGSIM-----------------ADNDVYDYVQ-ANAKLYGGEFGIHFHPHPLDW------LHITSSFETVTGLKE---------  621
Consensus_aa:             hs.FbsphpshIp..h........................s.hh...s..ph.lbGhEh.sph.hs..........hphshsh..s+..pp.........
Consensus_ss:             eeeeeeeeeeeeeee                           eeeeeee  eeeeeeeeeeeeee   ee    eeeeeeeeeeeeeee
DOMAINS                                           L8

Conservation:                                 5    5        5                                                 65  55
ShuA_Sd              558  ----------------GEYISSINPDTVTSTLNIPIAHS----GFSVGWVGTFADRSTHISS--------------SYSKQPGYGVNDFYV--SYQGQQA  621
3fhh_chainA_p001     522  ----------------GEYISSINPDTVTSTLNIPIAHS----GFSVGWVGTFADRSTHIS-----------------SKQPGYGVNDFYV--SYQGQQA  582
PhuR_Pa              635  ----------------GEPINSVNPLTGVFGLGYDEADG----NYGGLLSWTLVKRKDRVDDSTFHTPD----GTASQFKTPGFGVLDLSA--YYRLS--  706
HutA_Vc              583  ----------------GKPLNSVSPWNAVTGINYDSEN-----NWGTAVNLTYTAKKKASEIN------------GDYQPISSATVIDVTA--YYKPI--  645
HpuB_Nm              675  ---------------QETPINALSPWSAVYNLGYDAPSK----RWGINAYATRTAAKKPSDTVHSNDDL--NNPWPYAKHSKAYTLFDLSA--YLNIG--  749
HgbA_Hd              832  TNK------------LDTPMNAIQPQTHVVGVGYEHPQE----KFGVDMYLTHASAKKEKDTFNMFYDGKDQKDQHIKWRSDRYTLVDLIA--YVKPV--  911
HmuR_Pg              535  ---------------TEMKLSNTRPHAATATLSYGHNFSKNY-RLASNFSTRFLSDLKTGRMNS--------NELFEEITYPGYTICRLDI--ENHFY--  606
HfpR_Fp              568  IRDSQTLETIKINKVAYFGLFNRSKHTANVKFLYMIPKI----KTDINLRVFYRSKYGLFDTNNNQ------ILDVYDDFVKGYFITNVSI--SKQIK--  653
BfpR_Fp              622  ---------------NGEYLPLIPANKWNNNIRTEFNLNNWIKKGFATLNIESIFKQNNNST--------------FETKSKDYTLVNVGLGGNININ--  690
Consensus_aa:             ...................lsshps.shs.sl.h..s......phs.sh.hp.h..bp..p.....................s@slhsl.h....p....
Consensus_ss:                                      eeeeeeeeeee       eeeeeeeeeeeeeeeee                  eee  eeeeeeee  eeee
DOMAINS                                 L9                                                     L10

Conservation:              5   5      9  6  5                                79
ShuA_Sd              622  LK-GMTTTLVLGNAFDKEYWSPQGI-----------------------PQDGRNGKIFVSYQW  660
3fhh_chainA_p001     583  LK-GMTTTLVLGNAFDKEYWSPQGI-----------------------PQDGRNGKIFVSYQW  621
PhuR_Pa              707  -K-DLTLNAGLYNLTDKKYWLWDDVRGYDSVGEASALAP---ANIDRLSQPGRNFAVNLVWDI  764
HutA_Vc              646  -K-DLTLRAGVFNLTDEEYYNWNDVRGLPS-------------EDKDKTQAKRNFGITAKYEF  693
HpuB_Nm              750  -K-QVTLRAAAYNITNKQYYTWESLRSIREFGTVNRVDNKTHAGIQRFTSPGRSYNFTIEAKF  810
HgbA_Hd              912  -K-NVTLRAGVYNLTNREYGTWDSIRSIRPFGTTNLINQETGKGIKRFNAPGRNFRVNAEITF  972
HmuR_Pg              607  -R-NYTLHIGAENLFDYKPKALAFN---------------------SPTSPGRIIYGLVRITF  646
HfpR_Fp              654  -E-KFSFQVGANNLLDYTNPKEI------------------------SNLSGRQIFGKLQFNF  690
BfpR_Fp              691  -KLKFDISINANNMLNKSYISHLSRLK-----------------TDRISNIGRNIVLGVSFFL  735
Consensus_aa:             .+.phohphsh.Nhhsbp....s.........................s.sGRph...hphph
Consensus_ss:                 eeeeeeeee     eee                               eeeeeeeeee
DOMAINS                                               L11
```

**B.** The FRAP/NxxL motifs identified in loop L7 of heme/hemoglobin TBDRs are not conserved in THC0290_0681 and THC0290_0369 sequences.

```
Conservation:                     5 666 8   6                                              6  9558
ShuA_Sd           415  GMTINPTNWLMLFGSYAQAFRAPTMGEMYNDSKHFSIGRFYTNYWVP--------------------------NPNLRPE  468
3fhh_chainA_p001  379  GMTINPTNWLMLFGSYAQAFRAPTMGEMYNDSKHFSIGRFYTNYWVP--------------------------NPNLRPE  432
PhuR_Pa           491  GVTYDFAQHYTWYGQYAQGFRTPTAKALYGRFENLQA----GYHIEP--------------------------NPNLKPE  540
HutA_Vc           441  GTVYKLNQENRLFAQISQGFRAPDFQELYYSFGNPAH----GYVFKP--------------------------NPNLEAE  490
HpuB_Nm           516  GFDWRFTKHLHLLAKYSTGFRAPTSDETWLLFPHP------DFYLKA--------------------------NPNLKAE  563
HgbA_Hd           673  TFSFDPMDFLKIQAKYATGFRAPTSDEIYFVFQHP------SFSIYP--------------------------NLYLKAE  720
THC0290_0369      657  NLKYGVSDKANIRFATSKTYTKPVIMEAFPLTYLNAD----GTSIQG--------------------------NSLLKNS  706
HmuR_Pg           402  SAMYKCS-HVTNRLSYAEGYRAPSLQEMYFFFNHG------AFFIYG--------------------------NPDLKPE  448
HfpR_Fp           396  AINYKVTDHLSLKTAVGYGFKAPDFRQLYFDFTNSSV----GYTVLGYNVADTRLNALDNQGQILKRIKDISFDNPLKPE  471
BfpR_Fp           486  GYKTKLTDNLSFRLNVASGFRSPNLAELTSNGVHEGS----NRYEIG--------------------------NSNLKNE  535
THC0290_0681      396  HARYNPWEGGVLRFSAGRGKRSANIFAENQQLFGSSR----SFNILESS----------------------GKIYGLNPE  449
Consensus_aa:          shphp..p...h...htpt@+sPsh..hh............s..hbs..........................s..LcsE
Consensus_ss:          eeeeee   eeeeeeeeeeee   hhhh               ee                                  e
DOMAINS                L7
```

### *Supplementary Tables*

#### T1. Strains and plasmids.

| Strain or plasmid | Description[(a)] | Source or reference |
|---|---|---|
| **Strains** | | |
| *E. coli* DH5α Z1 | Strain used for cloning | Expressys |
| *E. coli* MFD*pir* | Strain used for conjugation | [5] |
| *E. coli* Top10 | Strain used for heterologous protein expression | Invitrogen |
| *F. psychrophilum* strain OSU THCO2-90 | Wild type | [6] |
| *F. psychrophilum* strain TRV311 | Δ*hfpR* in strain OSU THCO2-90 | This study |
| *F. psychrophilum* strain TRV347 | Δ*hfpY* in strain OSU THCO2-90 | This study |
| *F. psychrophilum* strain TRV405 | Δ*bfpR* in strain OSU THCO2-90 | This study |
| *F. psychrophilum* strain TRV403 | Δ*hfpR*Δ*bfpR* in strain OSU THCO2-90 | This study |
| *F. psychrophilum* strain TRV452 | Δ*hfpR* ecto::*hfpR* in strain OSU THCO2-90 | This study |
| *F. psychrophilum* strain TRV458 | Δ*bfpR* ecto::*bfpR* in strain OSU THCO2-90 | This study |
| *F. psychrophilum* strain TRV459 | Δ*hfpR*Δ*bfpR* ecto::*bfpR-hfpR* in strain OSU THCO2-90 | This study |
| *F. psychrophilum* strain TRV491 | Δ*hfpY* ecto::*hfpY* in strain OSU THCO2-90 | This study |
| *F. psychrophilum* strain TRV421 | ecto::$P_{rpsL}$-*mChFp* in strain OSU THCO2-90 | This study |
| *F. psychrophilum* strain TRV422 | ecto::$P_{rpsL}$-FP0506 in strain OSU THCO2-90 | This study |
| **Plasmids** | | |
| pCP*Gm*$^r$ | *E. coli–F. psychrophilum* shuttle plasmid; ColE1 *ori* (pCP1 ori), Ap$^r$ (Gm$^r$) | [7] |
| pCP*Gm*$^r$-$P_{less}$-mCh (pGi47) | pCP*Gm*$^r$ carrying a promoter less codon-adapted *mCherry* gene; Ap$^r$ (Gm$^r$) | [2] |
| pGi48 | pGi47 derivative with *mCherry* under the transcriptional control of *hfpR* promoter region (240 bp) | This study |
| pGi49 | pGi47 derivative with *mCherry* under the transcriptional control of *bfpR_1* promoter region (284 bp) | This study |
| pGi74 | pGi47 derivative with *mCherry* under the transcriptional control of *bfpR_2* promoter region (609 bp) | This study |
| pGi80 | pGi47 derivative with *mCherry* under the transcriptional control of *bfpR_3* promoter region (950 bp) | This study |
| pYT313 | Suicide vector carrying *sacB*; Ap$^r$ (Em$^r$) | [1] |
| pGi27 | pYT313 derivative used to delete *hfpR* | This study |
| pGi29 | pYT313 derivative used to delete *hfpY* | This study |
| pGi38 | pYT313 derivative used to delete *bfpR* | This study |
| pGi37 | pYT313 derivative used for chromosomal integration of $P_{rpsL}$-mChFp reporter fusion at the neutral intergenic site (between 1121879 and 1124880 of OSU THCO2-90 genome) | This study |
| pGi39 | pYT313 derivative used for cloning gene into the chromosomal expression platform, carrying 2-kb fragments upstream and downstream of the neutral intergenic site and a K7 composed of $P_{rpsL}$, a multiple cloning site (NotI, SpeI, SacI, SalI, XmaI) and an intrinsic terminator | This study |
| pGi42 | pGi39 derivative used for chromosomal integration of $P_{rpsL}$-FP0506 | This study |
| pGi44 | pGi39 derivative used for chromosomal integration of $P_{hfpR}$-*hfpR* | This study |
| pGi45 | pGi39 derivative used for chromosomal integration of $P_{bfpR\_1}$-*bfpR* | This study |
| pGi46 | pGi39 derivative used for chromosomal integration of $P_{bfpR\_1}$-*bfpR*-$P_{hfpR}$-*hfpR* | This study |
| pGi55 | pGi39 derivative used for chromosomal integration of $P_{hfpR}$-*hfpY* | This study |
| pMAL-c4X | Maltose binding protein (MBP) vector, P*tac* promoter, M13 *ori*, Ap$^r$ | New England Biolabs |
| pMBP-HfpY | Expression of N-terminal MBP-tagged HfpY cloned into pMAL-c4X | This study |

[(a)] Antibiotic resistance markers: ampicillin, Ap$^r$ is expressed in *E. coli* and erythromycin (Em$^r$) and gentamicin (Gm$^r$) in *F. psychrophilum*. The replication origin in parentheses is active in *F. psychrophilum*.

### T2. List of oligonucleotides.

| Name | Sequence 5'->3' | PCR Amplification |
|---|---|---|
| **Plasmids allowing mutant construction** | | |
| TRO186 | AAATGTGCGCGGAACCCCTA | pYT313 vector |
| TRO187 | CGGCACATAACAAACAAATTGGC | pYT313 vector |
| TRO336 | TAGGGGTTCCGCGCACATTTCAGCACAGCCCAGAAATTATC | Upstream region of *hfpR* for pGi27 |
| TRO337 | GAAAATTTGGTGCAAACCAAAAGCTTTATTTATA | Upstream region of *hfpR* for pGi27 |
| TRO338 | GCTTTTGGTTTGCACCAAATTTTCGGAAAATTGCAA | Downstream region of *hfpR* for pGi27 |
| TRO339 | GCCAATTTGTTTGTTATGTGCCGTTCGTCCGAATTATAGCCTGTT | Downstream region of *hfpR* for pGi27 |
| TRO358 | TAGGGGTTCCGCGCACATTTCGCTAGCTTTAACACACAT | Upstream region of *hfpY* for pGi27 |
| TRO359 | CTCACGCCACATTTATATTACATAATGAAAGTGTTAAAATATAAATG | Upstream region of *hfpY* for pGi27 |
| TRO360 | CATTTATATTTTAACACTTTCATTATGTAATATAAATGTGGCGTGAGATAG | Downstream region of *hfpY* for pGi27 |
| TRO361 | GCCAATTTGTTTGTTATGTGCCGGAGGCAATATAGGAACTGCTG | Downstream region of *hfpY* for pGi27 |
| TRO408 | TAGGGGTTCCGCGCACATTTGTCGAGTTTCACATCGTTATTGA | Upstream region of *bfpR* for pGi38 |
| TRO409 | CTCCCAAAACCATAGTATAGGGTAATTTTTATAGCGTTC | Upstream region of *bfpR* for pGi38 |
| TRO410 | ACCCTATACTATGGTTTTGGGAGTTAGTTTCTTCTTATAA | Downstream region of *bfpR* for pGi38 |
| TRO411 | CAATTTGTTTGTTATGTGCCGTGGCAAAGAATTTTTACGGATTA | Downstream region of *bfpR* for pGi38 |
| **Plasmids allowing chromosomal gene integration** | | |
| TRO390 | ATGCAAGCTTGGCGTAATC | pGi37 vector |
| TRO419 | CCGCTGCATAACCCTGCTT | pGi37 vector |
| TRO393 | GGTGCGAGGATCCTCCTAACGCAAAATACTGTAATTAGTA | Upstream of insertion site for pGi37 |
| TRO420 | GAAGCAGGGTTATGCAGCGGTCAAATTACATCATCATAAGTCCA | Upstream of insertion site for pGi37 |
| TRO396 | AAGCGGGCTTATTGCTGCAGATTGAAGTGAAAATCATTTTTTTTAAG | Downstream of insertion site for pGi37 |
| TRO397 | GATTACGCCAAGCTTGCATAGAAACACACGATAAAAAAGAAATAAC | Downstream of insertion site for pGi37 |
| TRO394 | GTTAGGAGGATCCTCGCACCACACTTTTTCAAAC | $P_{rpsL}$-mChFp K7 for pGi37 |
| TRO395 | CTGCAGCAATAAGCCCGCTTTTGCG | $P_{rpsL}$-mChFp K7 for pGi37 |
| TRO245 | CTCCTTTGCTTACCATAATTAAATACTAAAAATTACTTGTTTATAAA | pCP23 fragment for pGi19 |
| TRO246 | caaataggcctagctgactttcc | pCP23 fragment for pGi19 |
| TRO247 | ATTTTTAGTATTTAATTATGGTAAGCAAAGGAGAAGAAGATAACATG | mChFp gene for pGi19 |
| TRO248 | AAAGTCAGCTAGGCCTATTTGTATAATTCATCCATTCCTCCG | mChFp gene for pGi19 |
| TRO125 | GTCAGCTAGGCAATTAAATACTAAAAATTACTTGTTTATAAAAT | pCP23 fragment for pCP23-$P_{rpsL}$ |
| TRO126 | TTAGTATTTAATTGCCTAGCTGACTTTCCGC | pCP23 fragment for pCP23-$P_{rpsL}$ |
| TRO108 | GTGTGGTGCGATGAACTATCGCTTTGCGTCG | $P_{rpsL}$ fragment for pCP23-$P_{rpsL}$ |
| TRO111 | AAAAGCGGGCTTATTGCTCGGTCTTGCCTTGCTC | $P_{rpsL}$ fragment for pCP23-$P_{rpsL}$ |
| TRO443 | GCCTAGCTGACTTTCCGC | plasmid fragment for pGi39 |
| TRO444 | AATTAAATACTAAAAATTACTTGTTTATAAAATTCC | plasmid fragment for pGi39 |
| TRO421 | gcggccgccgactagtgagctcgtcgacccggggcctagctgactttccgc | MCS sequence for pGi39 |
| TRO422 | CCCGGGTCGACGAGCTCACTAGTCGGCGGCCGCAATTAAATACTAAAAATTACTTGTTTATAAAATTCC | MCS sequence for pGi39 |
| TRO434 | AATTAAATACTAAAAATTACTTGTTTATAAAATTCCCGC | plasmid fragment for pGi42 |
| TRO435 | GCCTAGCTGACTTTCCGCC | plasmid fragment for pGi42 |
| TRO436 | ATAAACAAGTAATTTTTAGTATTTAATTATGACAACAACCAAAAAAACCA | FP0506 fragment for pGi42 |
| TRO437 | GGGCGGAAAGTCAGCTAGGCTTATTGTAGATTAAAGTATTCTTGATCAGAC | FP0506 fragment for pGi42 |
| TRO404 | GCGATTTTGTTCTAATCACCCCAAA | chromosomal structure analysis |
| TRO405 | TGAGGTTTGGGTTTAAGAAAGCCAAA | chromosomal structure analysis |
| TRO402 | GAGCATGATAATTTAGTAGCAGAAGCTGGAG | chromosomal structure analysis |
| TRO403 | TCCTTAAGATGTACAGGCGCAAGAACC | chromosomal structure analysis |
| **pGi39 derivative plasmids allowing ectopic complementation of mutants** | | |
| TRO443 | GCCTAGCTGACTTTCCGC | pGi39 vector |
| TRO445 | GGATCCTCCTAACGCAAAATAC | pGi39 vector |
| TRO446 | GTATTTTGCGTTAGGAGGATCCTCGCTAAAGTTAAACTTTACTTCAT | $P_{hfpR}$-*hfpR* for pGi44 |

| | | |
|---|---|---|
| TRO447 | GGCGGAAAGTCAGCTAGGCTTAAAAATTAAATTGCAATTTTCCGAA | $P_{hfpR}$-*hfpR* for pGi44 and pGi46 |
| TRO448 | GTATTTTGCGTTAGGAGGATCCTCCTCCTAGTTTTATGTTTTAATTTATTT | $P_{bfpR\_1}$-*bfpR* for pGi45 and pGi46 |
| TRO449 | GGCGGAAAGTCAGCTAGGCTTATAAGAAGAAACTAACTCCCAAAAC | $P_{bfpR\_1}$-*bfpR* for pGi45 |
| TRO452 | GTTTAACTTTAGCGATTATAAGAAGAAACTAACTCCCAAAAC | $P_{hfpR}$-*hfpR* for pGi46 |
| TRO453 | GTTAGTTTCTTCTTATAATCGCTAAAGTTAAACTTTACTTCAT | $P_{bfpR\_1}$-*bfpR* for pGi46 |
| TRO519 | CATTCTTAATTATTTTCTGCAAATATATGTCC | $P_{hfpR}$ for pGi55 |
| TRO520 | TCATTTTAAGCCTAGCTGACTTTCCG | $P_{hfpR}$ for pGi55 |
| TRO521 | GGACATATATTTGCAGAAAATAATTAAGAATGAAAAATCATTTTTTTAAAGTTGCAG | *hfpY* for pGi55 |
| TRO522 | GAAAGTCAGCTAGGCTTAAAATGATGTCGTGCTATTAGC | *hfpY* for pGi55 |

**Construction of pCP$Gm^r$ derivative replicative plasmides**

| | | |
|---|---|---|
| TRO454 | ACATCGACACCAGAACTATCGCTTTGCGTCGTTC | pCP$Gm^r$ vector |
| TRO455 | ATTATACAAATAGGAAAAGGCTCCTGTTTTTGAGC | pCP$Gm^r$ vector |
| TRO456 | GCGATAGTTCTGGTGTCGATGTAAGAAATCAAGTT | *mCherry* gene for pGi47 |
| TRO457 | CTCGAGACTAGTCTTAAGCGGCCGCTAGCCACTGTTTGCTAAGTGAGCTAGAC | *mCherry* gene for pGi47 |
| TRO458 | CCGCTTAAGACTAGTCTCGAGAAAAATTATGGTAAGCAAAGGAGAAGAAGATAA | Rho independent terminator for pGi47 |
| TRO459 | AACAGGAGCCTTTTCCTATTTGTATAATTCATCCATTCCTCCG | Rho independent terminator for pGi47 |
| TRO460 | CTCGAGAAAAATTATGGTAAGCAAAGG | pGi47 vector |
| TRO461 | ACTAGTCTTAAGCGGCCGCTA | pGi47 vector |
| TRO462 | TAGCGGCCGCTTAAGACTAGTTCGCTAAAGTTAAACTTTACTTCAT | $P_{hfpR}$ for pGi48 |
| TRO463 | CCTTTGCTTACCATAATTTTTCTCGAGTCTTAATTATTTTCTGCAAATATATGTCC | $P_{hfpR}$ for pGi48 |
| TRO464 | TAGCGGCCGCTTAAGACTAGTTCCTCCTAGTTTTATGTTTTAATTTATTT | $P_{bfpR\_1}$ for pGi49 |
| TRO465 | CCTTTGCTTACCATAATTTTTCTCGAGAGTATAGGGTAATTTTTATAGCGTTC | $P_{bfpR\_1}$ for pGi49 |
| TRO789 | TAGCGGCCGCTTAAGACTAGTCTCATTATTAGCGATGGTAGTATTG | $P_{bfpR\_2}$ for pGi74 |
| TRO790 | CCTTTGCTTACCATAATTTTTCTCGAGAGTATAGGGTAATTTTTATAGCGTTC | $P_{bfpR\_2}$ for pGi74 and $P_{bfpR\_3}$ for pGi80 |
| TRO824 | TAGCGGCCGCTTAAGACTAGTGGTGTTGTGGTACAAGATAATAATGATAC | $P_{bfpR\_3}$ for pGi80 |

**RT-qPCR**

| | | |
|---|---|---|
| TRO497 | GGTCTACGCACTTGCGTTCT | Housekeeping gene *rpsA* forward |
| TRO498 | GTCCAAGGGTCTTGCGTCAT | Housekeeping gene *rpsA* reverse |
| TRO499 | TAGTGTTCCGCCACTTACCG | Housekeeping gene *frr* forward |
| TRO500 | CCTATTTTCGCGTCTTCGGC | Housekeeping gene *frr* reverse |
| TRO591 | GCTGGCTTTAGGTACGATAATCA | *hfpR* forward |
| TRO592 | TCTGCCACGTTATACCCCAA | *hfpR* reverse |
| TRO593 | ACGTTGAGGTTTTGAAGATGGA | *THC0290_1813* forward |
| TRO594 | AAACTTCGACCAACTCTGCG | *THC0290_1813* reverse |
| TRO595 | GTAAACGGAGGTGCTGCAAT | *hfpY* forward |
| TRO596 | CCAACCATTACCGCTCCCTA | *hfpY* reverse |
| TRO597 | TCATGCAACTTCAAGATTGGTTT | *THC0290_1811* forward |
| TRO598 | GAATTGGCTTCGATTTTAAATTTCAAAA | *THC0290_1811* reverse |
| TRO599 | CGGGAATCGTGTTTTGGTGT | *bfpR* forward |
| TRO600 | CCAACGCATCAGAGCCATAC | *bfpR* reverse |
| TRO863 | GCGATGGTAGTATTGTTTTCGAT | *THC0290_1678* forward |
| TRO864 | GTGGGTAATTTGAGAACCAGAA | *THC0290_1678* forward |

**3'-5' RACE experiments**

| | | |
|---|---|---|
| TRO806 | CCCGCACTTCGCCCCACAA | Reverse transcription *hfpR* |
| TRO807 | GGGAGCGGTAATGGTTGGTACAA | PCR1 *hfpR* and *hfpY* forward |
| TRO808 | CCCCACAAGTGGCACACCAT | PCR1 *hfpR* reverse |
| TRO809 | CTCCTCTTGCCGGAAAAGTATTTGT | PCR2 *hfpR* and *hfpY* forward |
| TRO810 | CCTTCTGCGCCACCAAAATCT | PCR2 *hfpR* reverse |
| TRO811 | CCCGATACAGGTCCAAACGACATA | Reverse transcription *hfpY* |
| TRO812 | CGACATACCTACTTGTGGTGCATT | PCR1 *hfpY* reverse |

| | | |
|---|---|---|
| TRO813 | GTGGTTGCGGAGTAAGAACAG | PCR2 *hfpY* reverse |
| TRO814 | GCCAATCCCTCCATGAGCGTT | Reverse transcription *bfpR* |
| TRO815 | CGAAACCGTAACTGGTCTGAAAGA | PCR1 *bfpR* forward |
| TRO816 | GCGGCTCCTTTTTGTTGTAATTCT | PCR1 *bfpR* reverse |
| TRO817 | CGCTGGTAAATGTAGGTTTAGGTGGAA | PCR2 *bfpR* forward |
| TRO818 | CTCAGGAATAGAAACCAAGACAGTTTCT | PCR2 *bfpR* reverse |
| TRO825 | CACCCGAACAAACAAAACCACCT | PCR1 *THC0290_1678* reverse |
| TRO826 | GGTGTTGGTGGGTAATTTGAGAACCA | PCR2 *THC0290_1678* reverse |
| TRO827 | GCCCATTCCAGAGAAGTTTCTCCA | PCR1 *asd* reverse |
| TRO828 | CTCCACCAGCAGAAAACAAAGCAA | PCR2 *asd* reverse |
| **OVERLAPPING RT-PCR** | | |
| TRO672 | GGGCATAGCGAAGCGG | *asd* forward |
| TRO673 | GCTCATAACTATGCACCGATTG | *THC0290_1678* reverse |
| TRO674 | GTTCGGGTGTTTCTTGTTTTC | *THC0290_1678* forward |
| TRO675 | CCACTTCAATCCCTCACGC | *bfpR reverse* |
| TRO676 | GAAAAGCCCGACCCGTAG | *upstream of bfpR forward* |
| TRO677 | ATTCCCGCTTAACCCACG | *bfpR* reverse |

**T3. Genomic organization and predicted protein functions of the iron-induced gene cluster encoding the Hfp system in strain OSU THCO2-90.**

| TU [a] | Locustag | Name | Annotation | DIP [b] | Plasma [b] | Blood [b] |
|---|---|---|---|---|---|---|
| | THC0290_1795 | | Di-heme oxidoreductase family lipoprotein | 1.2 | | -0.4 |
| ↑ | THC0290_1796 | *irpA* | Imelysin-like domain-containing lipoprotein | 1.7 | 1.0 | -2.3 |
| | THC0290_1797 | | Protein of unknown function precursor containing a T9SS-dependent Type A C-terminal domain | 1.8 | 2.3 | -2.8 |
| | THC0290_1798 | | Conserved exported ankyrin repeat-containing protein | 2.1 | 2.8 | -3.2 |
| | THC0290_1799 | | Putative lipoprotein | 2.1 | 2.8 | -2.5 |
| | THC0290_1800 | | Putative lipoprotein | 2.0 | 2.7 | -2.2 |
| | THC0290_1801 | | DUF3570 protein; putative ApbE substrate (DOI: 10.7554/eLife.66878) | 2.0 | 3.7 | -3.7 |
| | THC0290_1802 | | DUF4266 protein; putative redox cycling activity (DOI: 10.7554/eLife.66878) | 2.4 | 3.3 | -3.1 |
| | THC0290_1803 | *apbE* | Membrane-associated ApbE enzyme catalyzing the flavinylation of FMN-binding proteins | 1.9 | 3.5 | -2.9 |
| ↑ | THC0290_1804 | | Thioredoxin family protein | 2.0 | 3.3 | -3.6 |
| ↓ | THC0290_1805 | | Protein of unknown function | 1.8 | 1.9 | -2.4 |
| | THC0290_1806 | *irpA* | Imelysin-like domain-containing lipoprotein | 1.1 | 1.5 | -2.1 |
| | THC0290_1807 | *ggcX* | Vitamin K-dependent gamma-glutamyl carboxylase | 1.0 | 2.3 | -1.6 |
| | THC0290_1808 | *fecA* | Iron(III) dicitrate TonB-dependent receptor | 0.9 | 1.0 | -1.2 |
| | THC0290_1809 | | Acyl-CoA hydrolase family protein | | | |
| ↑ | THC0290_1810 | | PAS-domain protein, putative signaling protein | | | |
| | THC0290_1811 | | Putative hypothetical protein | 2.6 | 4.3 | -3.7 |
| | THC0290_1812 | *hfpY* | Heme binding lipoprotein, HmuY-like family | 2.7 | 0.7 | -3.0 |
| | THC0290_1813 | | Probable transmembrane protein of unknown function | 2.8 | 3.1 | -3.4 |
| ↑ | THC0290_1814 | *hfpR* | TonB-dependent outer membrane receptor | 2.2 | 4.5 | -3.0 |
| ↓ | THC0290_1815 | | Protein of unknown function precursor | 2.8 | 1.7 | -3.5 |
| | THC0290_1816 | | Protein of unknown function | 2.8 | 1.5 | -3.1 |

[(a)] Gene orientation on the chromosome (↑, forward strand; ↓ reverse strand) and [(b)]differential expression data were retrieved from Guérin *et al* [2] (https://fpeb.migale.inrae.fr) that provides microarray expression level along the genome, transcription start sites (TSSs) and differential expression analyses for diverse biological conditions. [(a)] Arrows indicate the transcriptional orientation of each gene cluster (TU); [(b)] Genes significantly upregulated (yellow) and down-regulated (blue) in iron-limited condition ('**DIP**'; TYES 2,2'-dipyridyl 25 µM), under rainbow trout plasma exposure ('**Plasma**') or in the presence of blood ('**Blood**'; TYES agar supplemented with 10% horse blood) are indicated. Values are log2-fold changes of the test condition compared to the relevant control condition.

### *Supplementary references*

1. Zhu Y, Thomas F, Larocque R, et al. Genetic analyses unravel the crucial role of a horizontally acquired alginate lyase for brown algal biomass degradation by *Zobellia galactanivorans*. Environ Microbiol. 2017 Jun;19(6):2164-2181.
2. Guérin C, Lee B-H, Fradet B, et al. Transcriptome architecture and regulation at environmental transitions in flavobacteria: the case of an important fish pathogen. ISME Communications. 2021 2021/07/07;1(1):33.
3. Rochat T, Pérez-Pascual D, Nilsen H, et al. Identification of a novel elastin-degrading enzyme from the fish pathogen *Flavobacterium psychrophilum*. Applied and environmental microbiology. 2019 Jan 11.
4. Pei J, Kim BH, Grishin NV. PROMALS3D: a tool for multiple protein sequence and structure alignments. Nucleic acids research. 2008 Apr;36(7):2295-300.
5. Ferrieres L, Hemery G, Nham T, et al. Silent mischief: bacteriophage Mu insertions contaminate products of *Escherichia coli* random mutagenesis performed using suicidal transposon delivery plasmids mobilized by broad-host-range RP4 conjugative machinery. Journal of bacteriology. 2010 Dec;192(24):6418-27.
6. Bertolini JM, Wakabayashi H, Watral VG, et al. Electrophoretic detection of proteases from selected strains of *Flexibacter psychrophilus* and assesment of their variability. Journal of aquatic animal health. 1994;6:224-233.
7. Pérez-Pascual D, Rochat T, Kerouault B, et al. More than gliding: Involvement of GldD and GldG in the virulence of *Flavobacterium psychrophilum*. Frontiers in microbiology. 2017;8:2168.
